# Middle-aged mice treated with GHK-Cu peptide administered intraperitoneally or intranasally show behavioral rescue but divergent hippocampal aging programs

**DOI:** 10.64898/2026.04.09.717524

**Authors:** Jordan Mazzola, Manuela Rosenfeld, Matthew Tucker, Jackson Wezeman, Warren Ladiges, Gerald Yu Liao

## Abstract

Age-related cognitive decline (ARCD) is driven by conserved biological mechanisms of aging, yet no gerotherapeutic directly targets these processes in the brain. Glycyl-L-histidyl-L-lysine complexed with copper (GHK-Cu) is an endogenous peptide with regenerative and anti-inflammatory properties that declines with age. Whether its effects on cognitive aging depend on delivery route or exposure duration remains unclear. Aged C57BL/6J mice (20-21 months) received GHK-Cu (15 mg/kg) via short-term intraperitoneal (IP; 5 days) or longer-term intranasal (IN; 8 weeks) administration. Hippocampal-dependent escape learning was assessed using a spatial navigation task. Molecular effects were evaluated using hippocampal immunohistochemistry and bulk RNA sequencing. Differential gene expression was analyzed using DESeq2 with false discovery rate (FDR) correction, and pathway-level changes were assessed via gene set enrichment analysis (GSEA). IN GHK-Cu improved escape latency across Trials 2-4 in both sexes (*P* < 0.05), whereas IP dosing produced a transient improvement in males during Trial 2 (*P* < 0.05) without sustained effects or improvement in females. IN treatment increased synaptophysin in females (*P* < 0.001) and decreased GFAP in both sexes (*P* < 0.01), while IP treatment reduced TGF-β, GFAP, and MCP-1 in males (*P* < 0.05) and decreased p21 in females (*P* < 0.0001). Transcriptomic analysis revealed distinct molecular programs. IN GHK-Cu induced coordinated suppression of oxidative phosphorylation (male NES -5.44, female NES -4.20; FDR < 0.0001) and MYC target pathways (female NES -4.31, FDR < 0.0001), with additional attenuation of PI3K-AKT-mTOR signaling in females (NES -3.15, FDR = 0.062). In contrast, IP treatment activated oxidative phosphorylation (female NES 4.97, FDR < 0.001), DNA repair (NES 5.58, FDR < 0.001), and MYC targets (NES 4.34, FDR = 0.002), indicating engagement of acute stress-response and repair pathways. GHK-Cu improves hippocampal-dependent learning in aged mice through distinct biological modes: IP exposure activates repair and stress-response pathways, whereas IN delivery induces sustained suppression of growth and mitochondrial metabolic signaling associated with aging biology. These findings demonstrate that functional cognitive improvement can arise from divergent molecular states and identify administrative route and exposure duration as key determinants of gerotherapeutic response.

## Introduction

Population aging is a major biomedical challenge, with age-related cognitive decline (ARCD) affecting nearly two-thirds of individuals by age 70 with increasing vulnerability to neurodegenerative disease [1-3]. ARCD reflects the cumulative impact of conserved aging hallmarks, including mitochondrial dysfunction, deregulated nutrient sensing, chronic inflammation, and cellular senescence, on neuronal integrity and synaptic plasticity, particularly within the hippocampus [4-5]. Despite its prevalence, no gerotherapeutic drug directly targets the biological drivers of brain aging. While lifestyle interventions remain foundational [6-7], scalable pharmacologic strategies that modulate core aging pathways are needed. Rodent models enable such efforts through integrated behavioral, histologic, and transcriptomic interrogation [8-9].

Glycyl-L-histidyl-L-lysine complexed with copper (GHK-Cu) is an endogenous tripeptide that declines with age and exhibits regenerative, anti-inflammatory, and antioxidant properties [10-11]. It modulates transforming growth factor-β (TGF-β) signaling and suppresses senescence-associated pathways including p21 and p53, positioning it as a candidate regulator of aging biology [12-13]. Evidence of blood-brain barrier (BBB) permeability and oxidative stress modulation further support its relevance to cognitive aging [14]. However, its central effects may depend on delivery kinetics: intraperitoneal (IP) administration enables rapid systemic exposure but may limit sustained brain concentrations [15-16], whereas intranasal (IN) delivery bypasses the BBB, enhancing central nervous system (CNS) uptake but potentially requiring prolonged administration to achieve stable remodeling [17-19].

Whether GHK-Cu efficacy is governed by pharmacokinetic route or by duration-dependent biological remodeling remains unknown. We hypothesized that short-term IP delivery activates an acute hippocampal repair program, whereas prolonged IN administration induces sustained homeostatic remodeling, with both converging on improved hippocampal-dependent escape learning. To test this, we integrated behavioral performance with hippocampal immunohistochemistry and transcriptomic profiling to determine whether functional rescue arises from shared or distinct molecular programs.

## Materials and Methods

### Animals

C57BL/6J mice (50 male, 50 female), 18 months of age, were obtained from the National Institute on Aging Aged Rodent Colony (Charles River Laboratories) and housed under specific pathogen-free conditions (14:10-hour light-dark cycle, *ad libitum* food and water) at the University of Washington. Animals were group housed (≤5/cage), provided environmental enrichment, and acclimated for three weeks prior to experimentation. Mice were randomized to IP or IN treatment paradigms. Health monitoring followed institutional standards, and all procedures were approved by the University of Washington Institutional Animal Care and Use Committee.

### Treatment and Route of Delivery

GHK was administered as a copper complex (GHK-Cu; Active Peptide, Cambridge, MA) at 15 mg/kg, selected based on efficacy and safety relative to known copper toxicity thresholds [20-21]. Solutions were prepared in sterile 0.9% saline under aseptic conditions.

For IP delivery, mice received once-daily injections (200 µL) for five consecutive days with weight-adjusted dosing. For IN delivery, mice received 15 mg/kg in 20 µL saline once daily for eight consecutive weeks via a custom atomization device to enhance nasal deposition and CNS uptake [21]. Control groups received volume-matched saline under identical conditions. All administrations were performed at consistent times of day.

### Box Maze

Spatial learning was assessed using the Box Maze paradigm [22]. The apparatus consisted of a brightly lit square enclosure with eight floor-level openings, one of which connected to a dark escape chamber. Mice were given up to 120 seconds to locate the escape tunnel; failure to escape was recorded as 120 seconds. Each mouse completed four consecutive trials with 30-second inter-trial intervals. Escape latency was the primary outcome. The apparatus was cleaned with 70% ethanol between trials.

### Immunohistochemistry

Formalin-fixed, paraffin-embedded brain sections (4 µm) were deparaffinized, rehydrated, and subjected to heat-mediated antigen retrieval. Following blocking, sections were incubated overnight at 4°C with primary antibodies against synaptophysin (1:250, Invitrogen MA5-16402), PSD95 (1:250, Abcam ab18258), phospho-SMAD2 (1:50, Invitrogen 44-244G), MCP-1 (1:800, Novus NBP1-07035), p21 (1:200, Abcam ab188224), TGF-β (1:50, Abcam ab215715), and GFAP (1:1500, Invitrogen: PA1-10019). Detection was performed using HRP-conjugated secondary antibodies and DAB chromogen (Abcam). Sections were imaged at 20×-40× magnification, and hippocampal staining was quantified as percent area positivity using QuPath (v0.4.3) [23].

### RNA Sequencing

Flash-frozen hippocampal tissue was processed for bulk RNA sequencing. Samples (Intraperitoneal: N = 12; Intranasal: N = 24) were homogenized in TRIzol, and RNA was extracted, purified (Qiagen RNeasy Mini Kit), and quality-checked (RIN ≥ 7.3). Libraries were sequenced on the Illumina NovaSeq X-Plus platform (Novogene Co., Ltd.). Reads were trimmed (Trim Galore), aligned to the mouse reference genome (mm10) using STAR, and analyzed for differential expression using DESeq2 with Wald testing and false discovery rate (FDR) correction (0.1). Gene set enrichment analysis (GSEA) was performed using hallmark gene sets (FDR < 0.1). Analyses were conducted using SciDAP (https://github.com/datirium).

### Data Analyses

Statistics were performed using GraphPad Prism (v10.2.3). Two-group comparisons were analyzed using unpaired two-tailed *t*-tests; multi-group comparisons used one- or two-way analysis of variance (ANOVA) with Geisser-Greenhouse correction as needed, followed by appropriate post hoc testing. A threshold of *P* < 0.05 was used. RNA-seq analyses were conducted as described above using DESeq2 and GSEA (FDR < 0.1).

## Results

### IN GHK-Cu for 8 weeks produced sustained improvement in escape learning across sexes, whereas IP administration for 5 days yielded transient, sex-dependent effects

IN GHK-Cu (8 weeks) improved escape performance in both sexes following initial exposure, with reduced escape latency across Trials 2-4 (*P*’s < 0.05) (Figure 1A-B). In contrast, short-term IP dosing (5 days) improved escape latency in males during Trial 2 (*P* < 0.05) (Figure 1C) but did not sustain improvement through Trials 3-4 and did not improve performance in females (*P*’s > 0.05) (Figure 1C-D).

**Figure 1.**
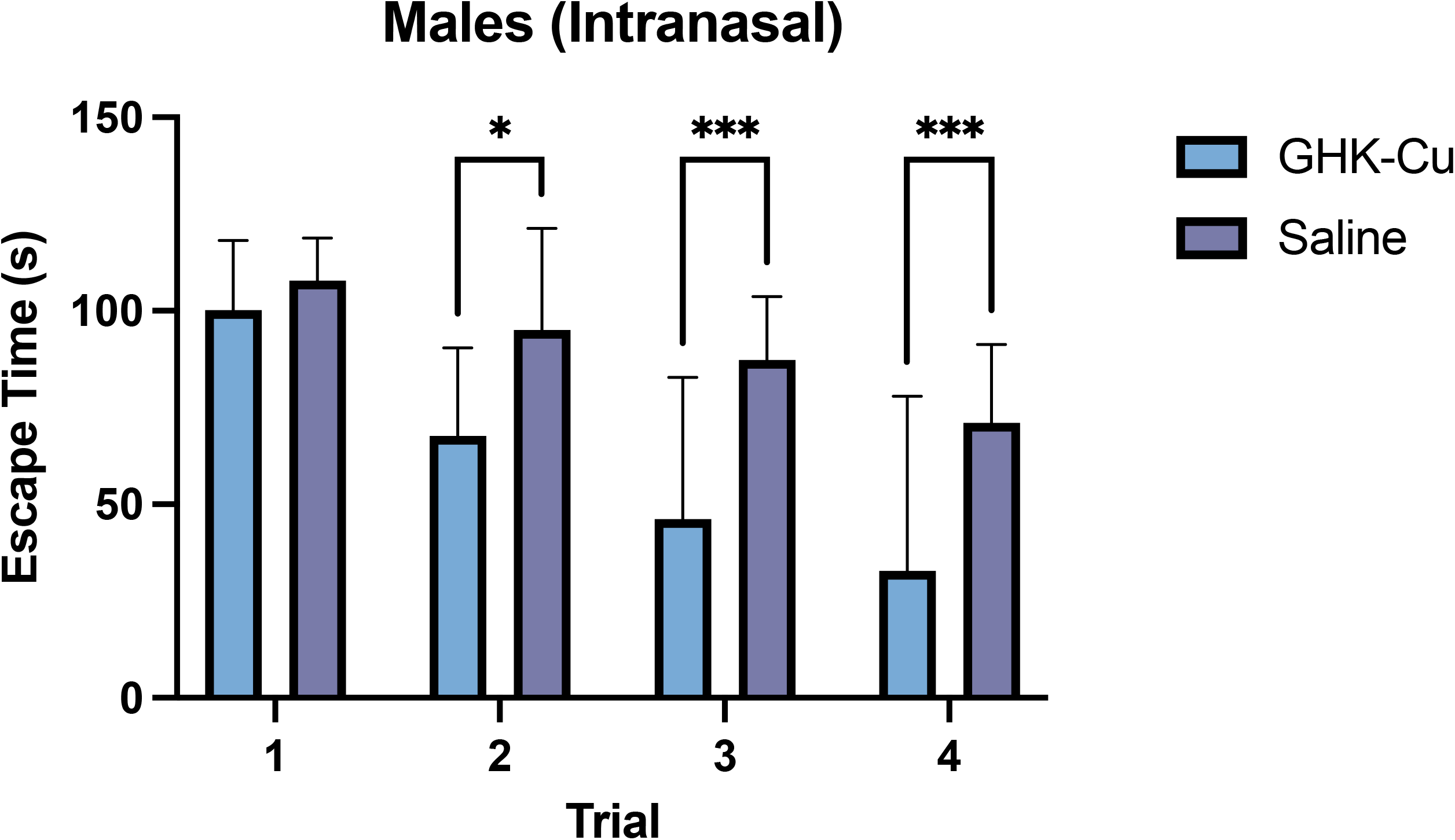
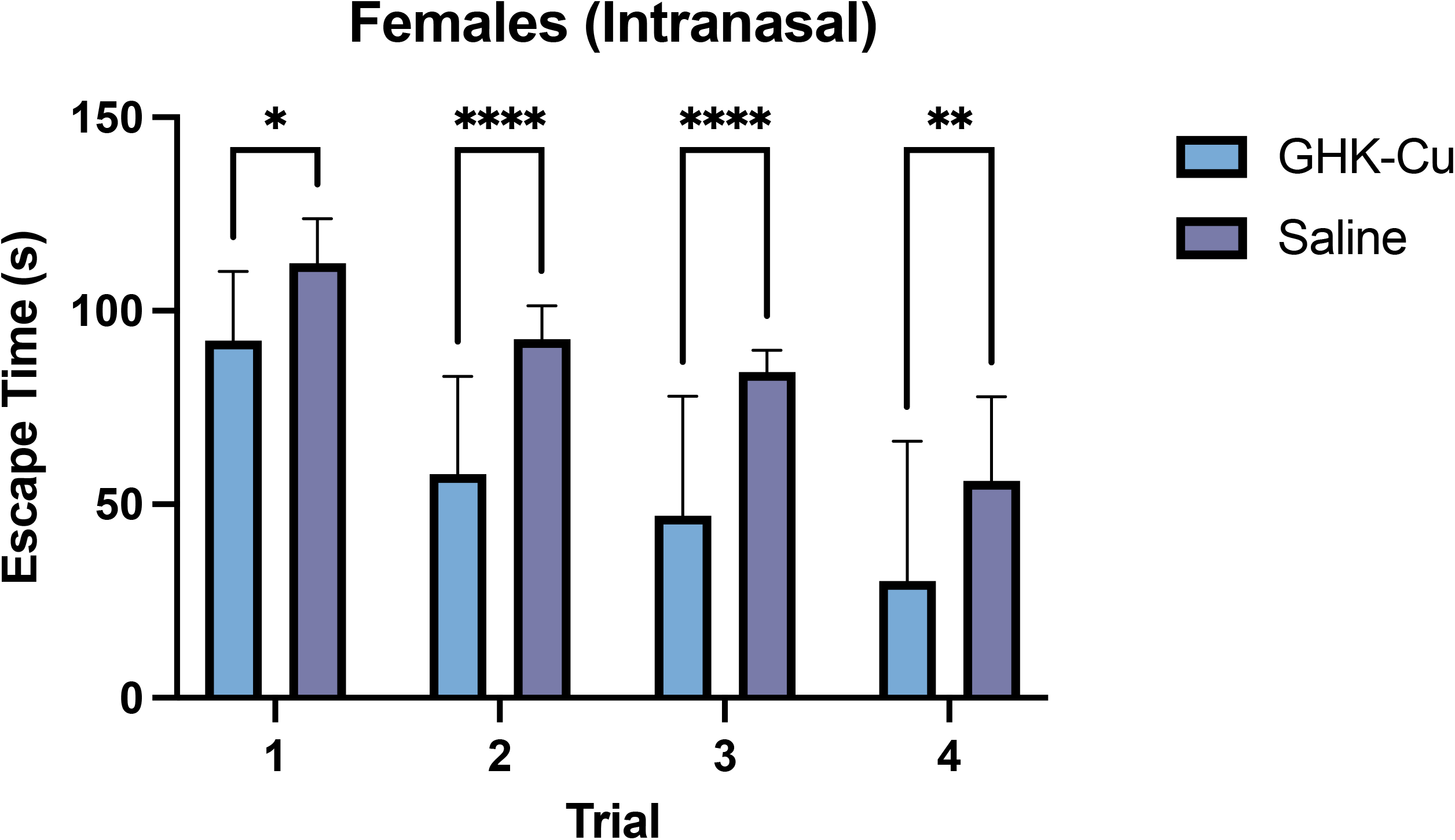
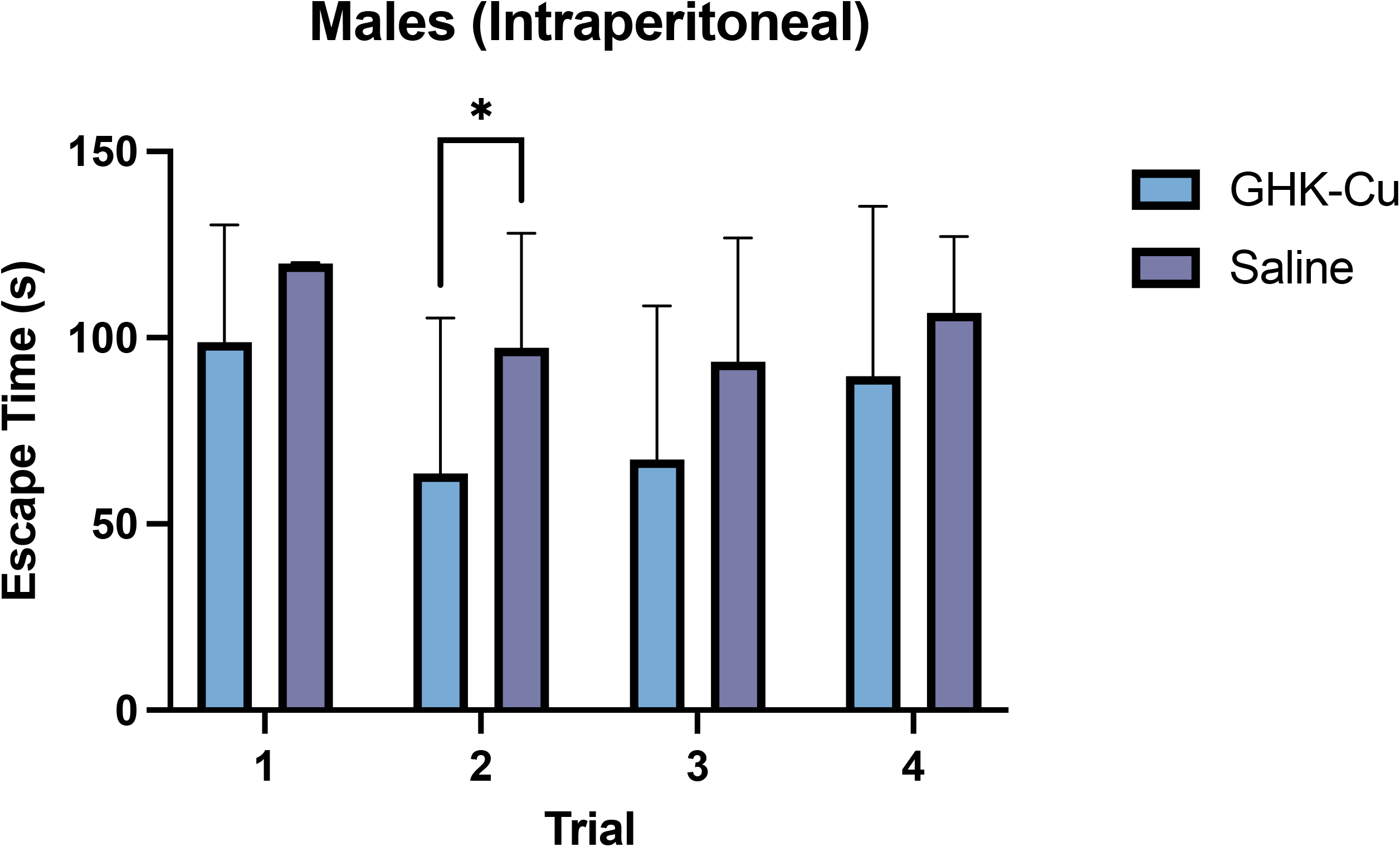
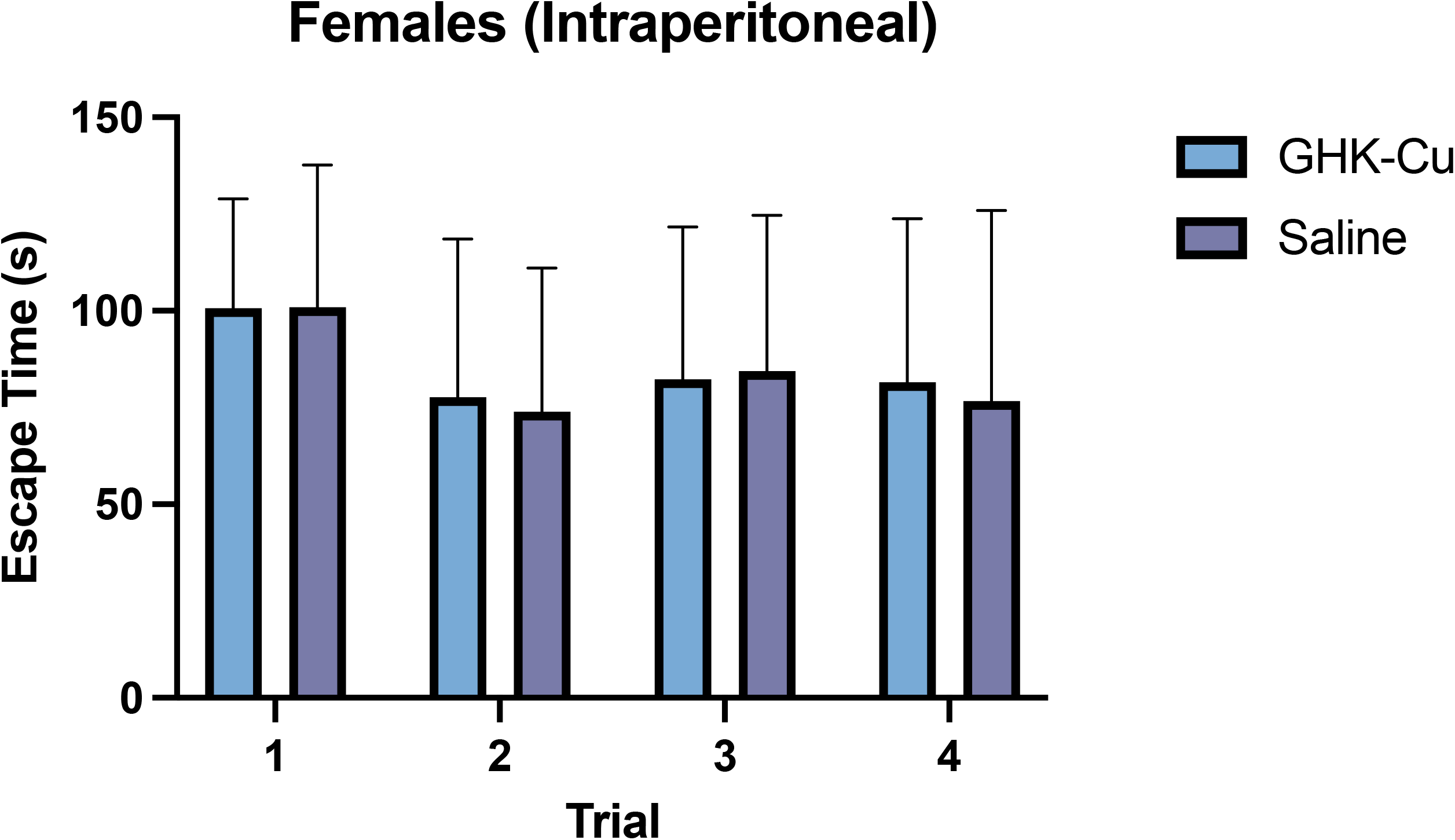
Box-Maze Escape Times Separated By Sex And GHK-Cu Induction Method. (A) Male and (B) Female Intranasal GHK-Cu escape times. (C) Male and (D) Female Intraperitoneal GHK-Cu escape times (*P < 0.05, \*\**P* < 0.01, \*\*\**P* < 0.001, \*\*\*\**P* < 0.0001).

### IN GHK-Cu engaged hippocampal chemokine suppression and synaptic remodeling programs

In the hippocampus, IN treatment increased synaptophysin in females (*P* < 0.001) (Figure 2A), while both sexes demonstrated modulation of GFAP (*P*’s < 0.01) (Figure 2B). No changes were observed in MCP-1, PSD95, or TGF-β (*P*’s > 0.05) (Figure 2C-F).

**Figure 2.**
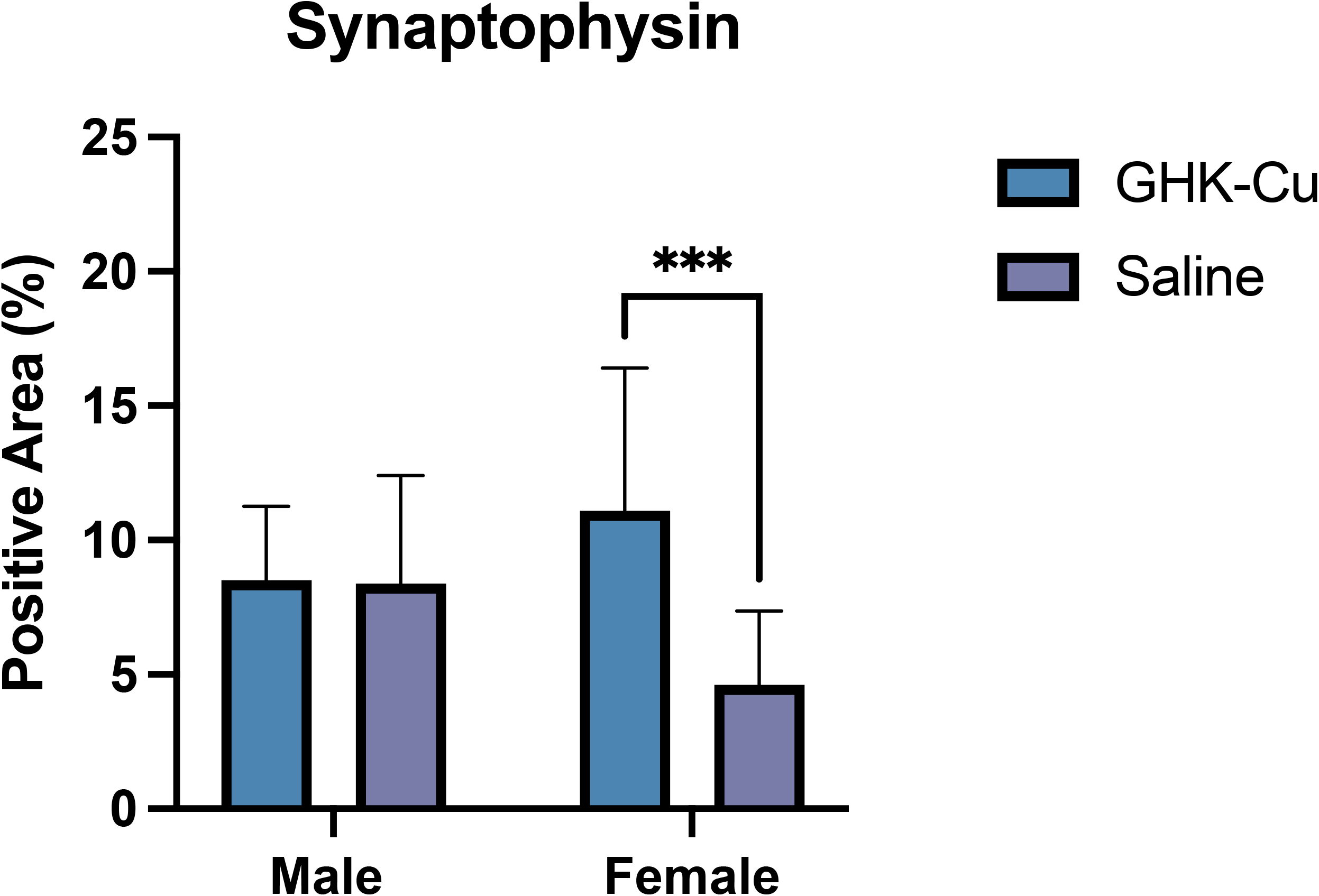
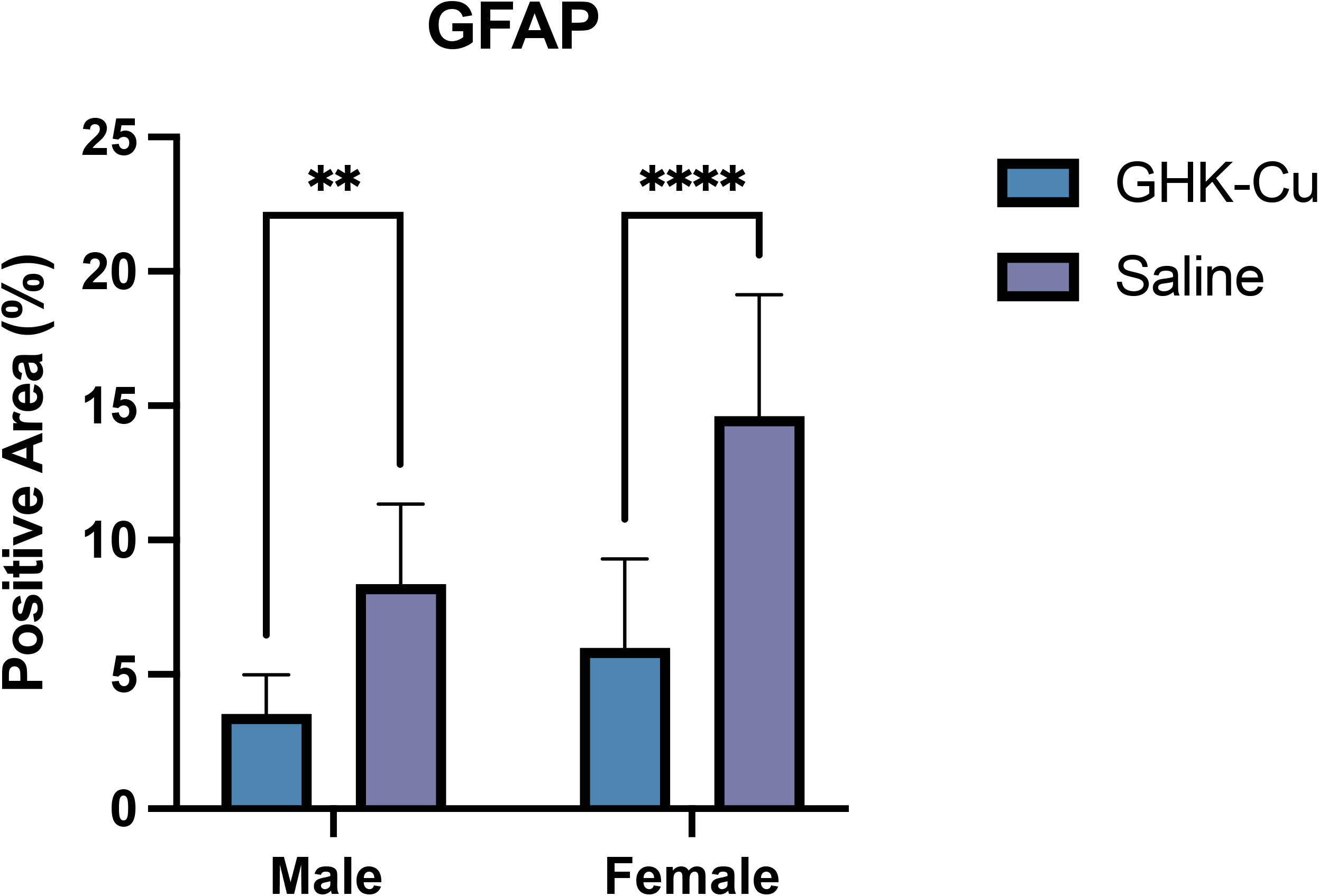
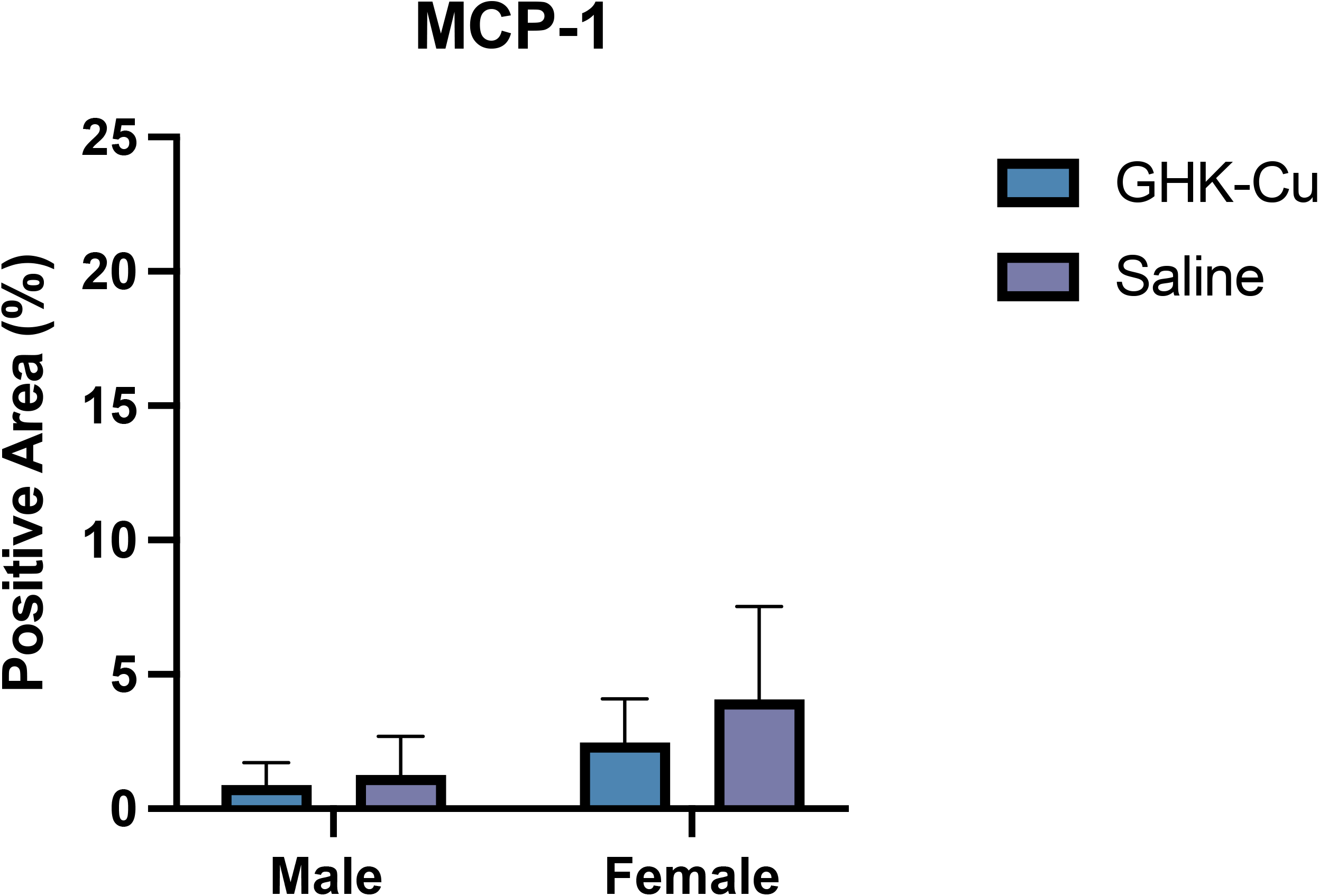
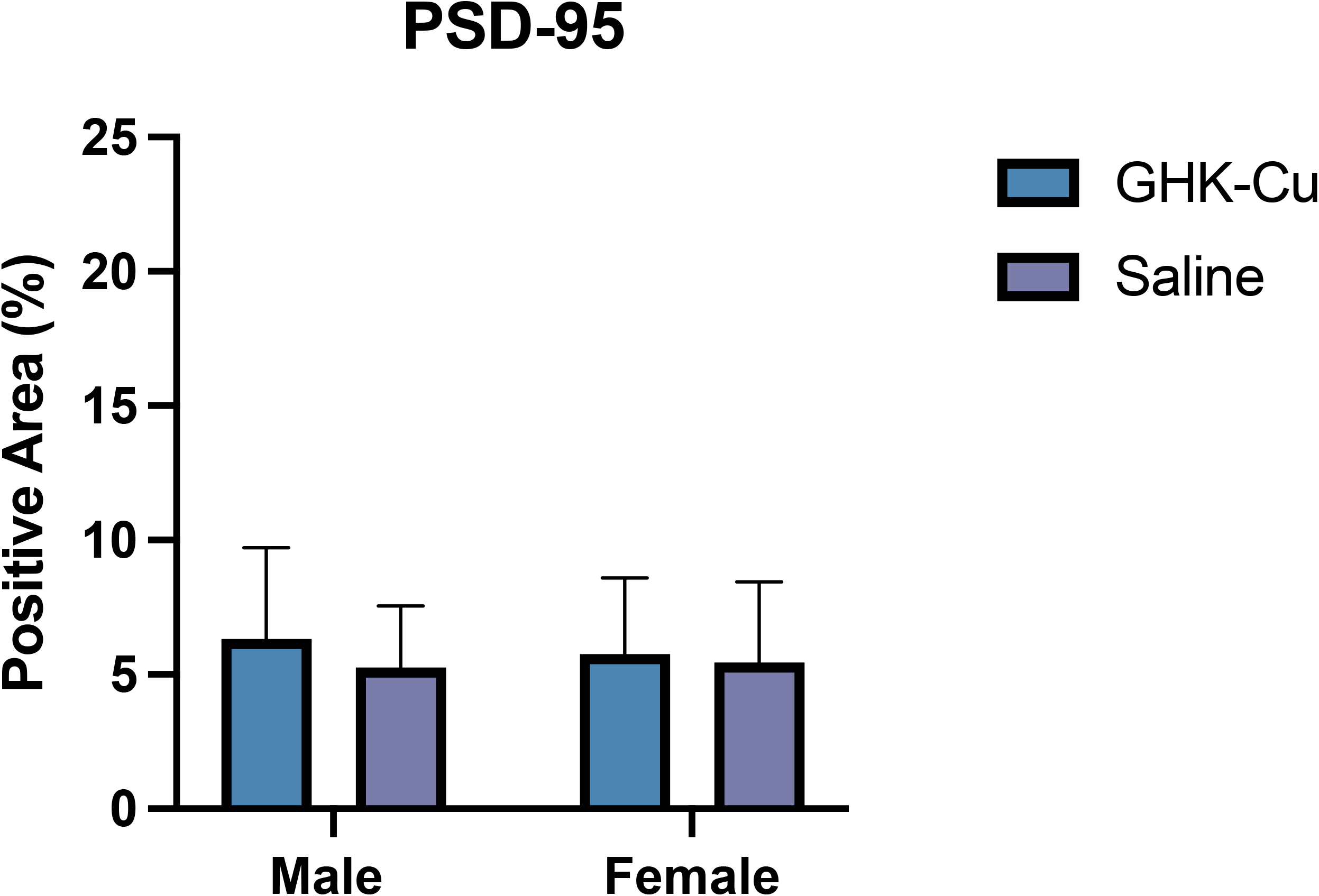
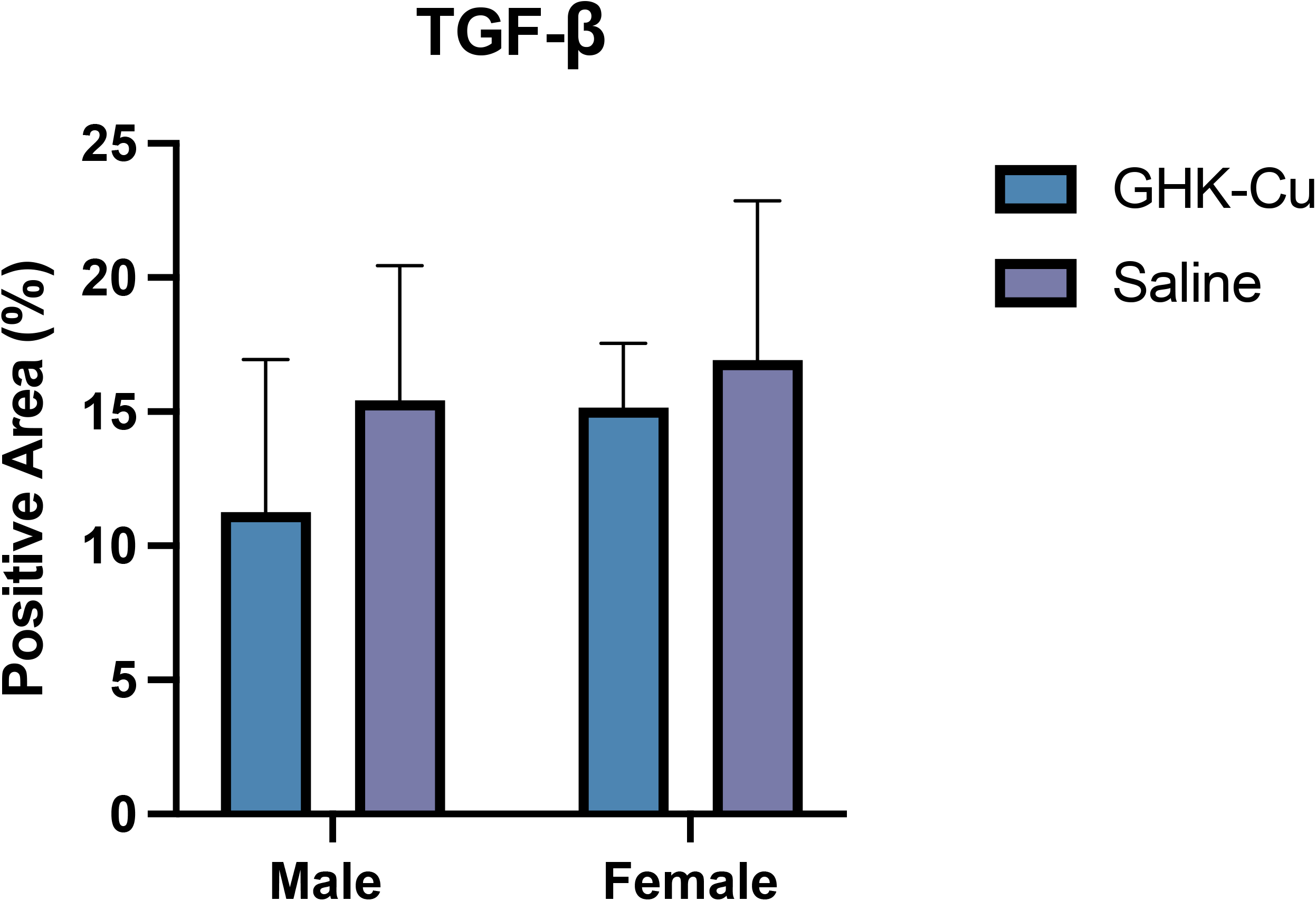
Positive area-staining by antibody in intranasally-treated GHK-Cu mice of both sexes for (A) Synaptophysin, (B) GFAP, (C) MCP-1, (D) PSD-95, and (E) TGF-β. \*\**P* < 0.01, \*\*\**P* < 0.001, \*\*\*\**P* < 0.0001.

### IP GHK-Cu engaged sex-specific hippocampal repair and stress-response molecules

Short-term IP dosing decreased TGF-β, GFAP, and MCP-1 in the male hippocampus (*P*’s < 0.05) (Figures 3A-C). p21 expression decreased in the female hippocampus (*P* < 0.0001) (Figure 3D). No changes were observed in synaptophysin, PSD-95, or pSMAD-2 (*P*’s > 0.05) (Figures 3E-G).

**Figure 3.**
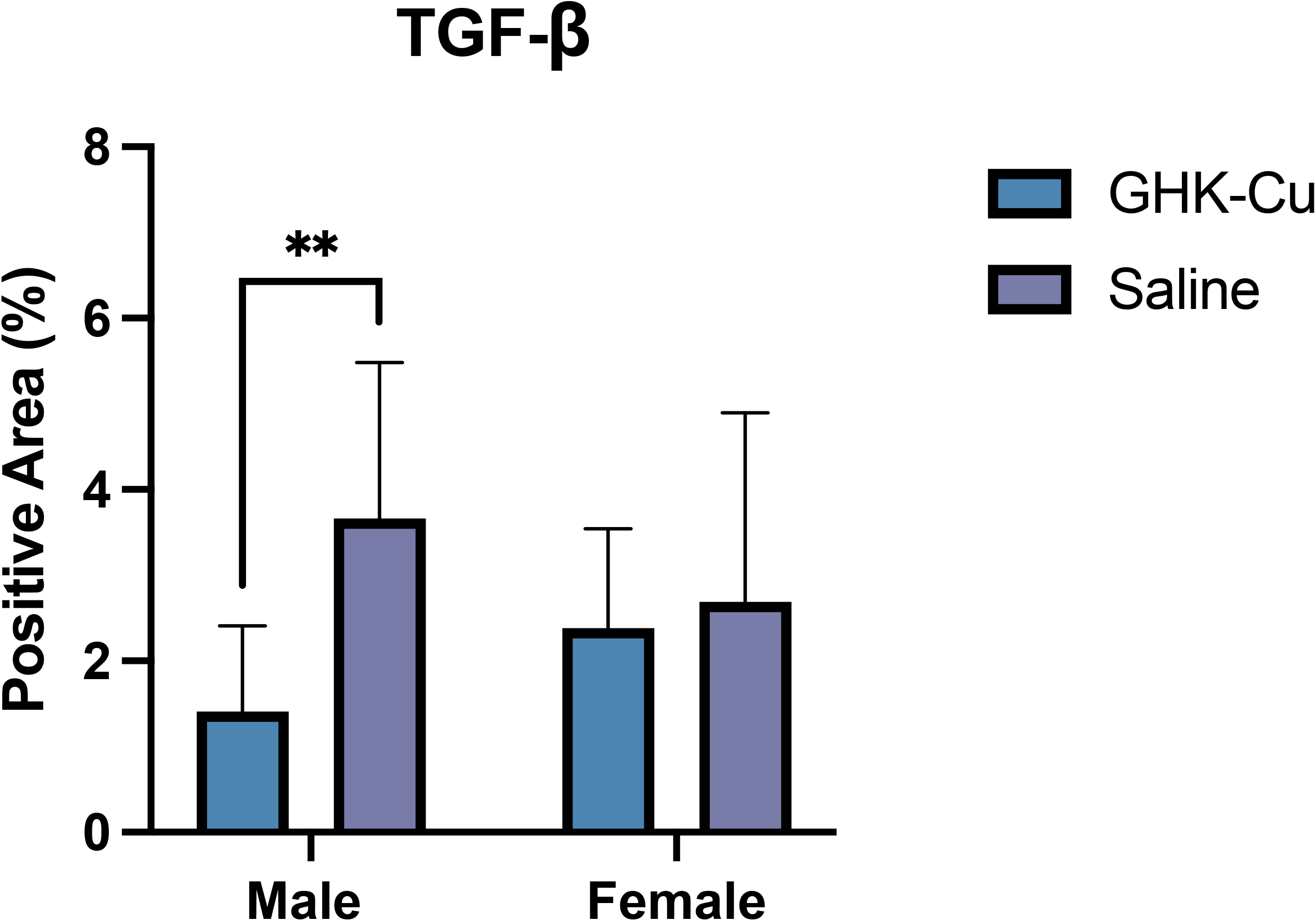
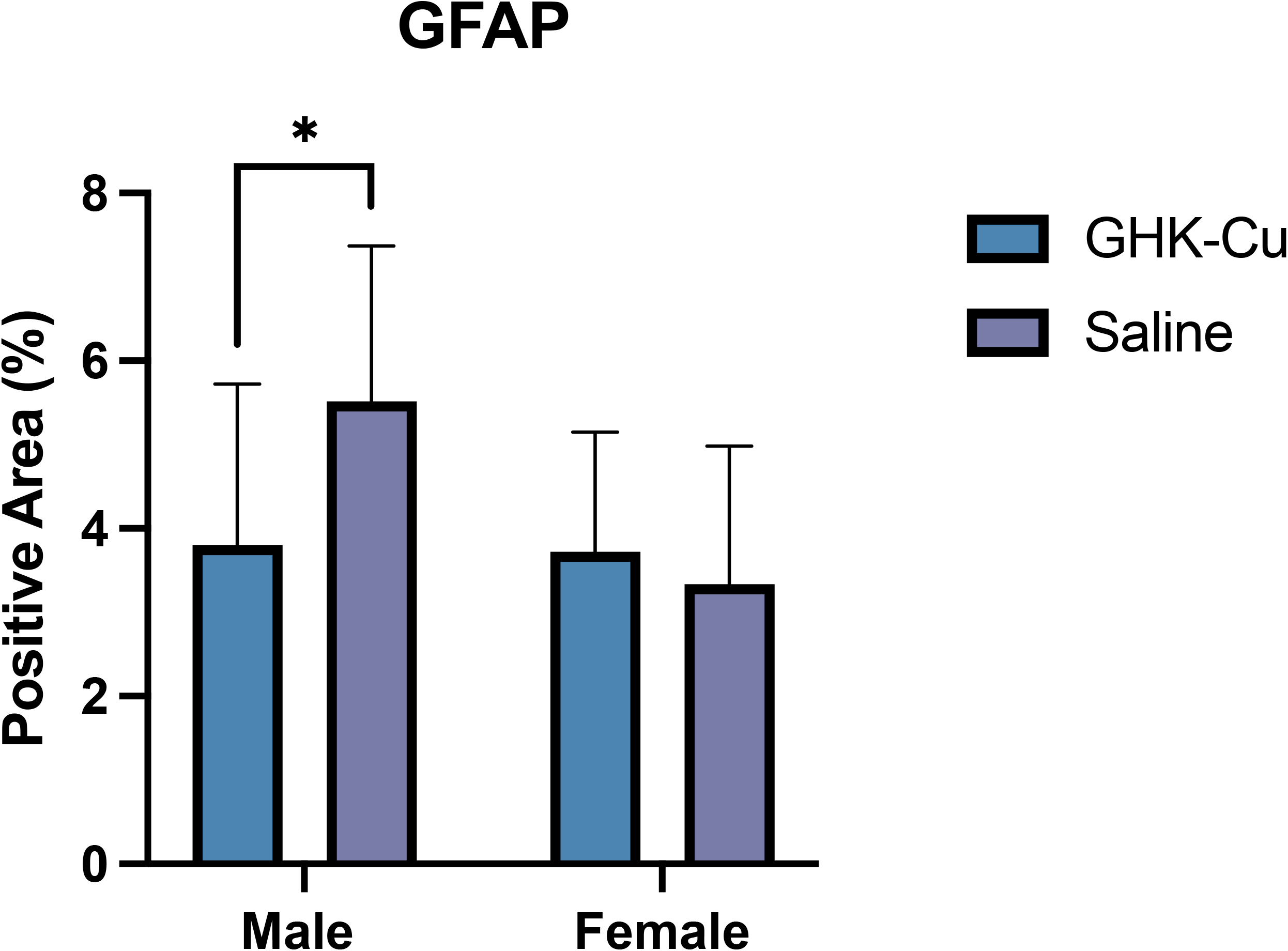
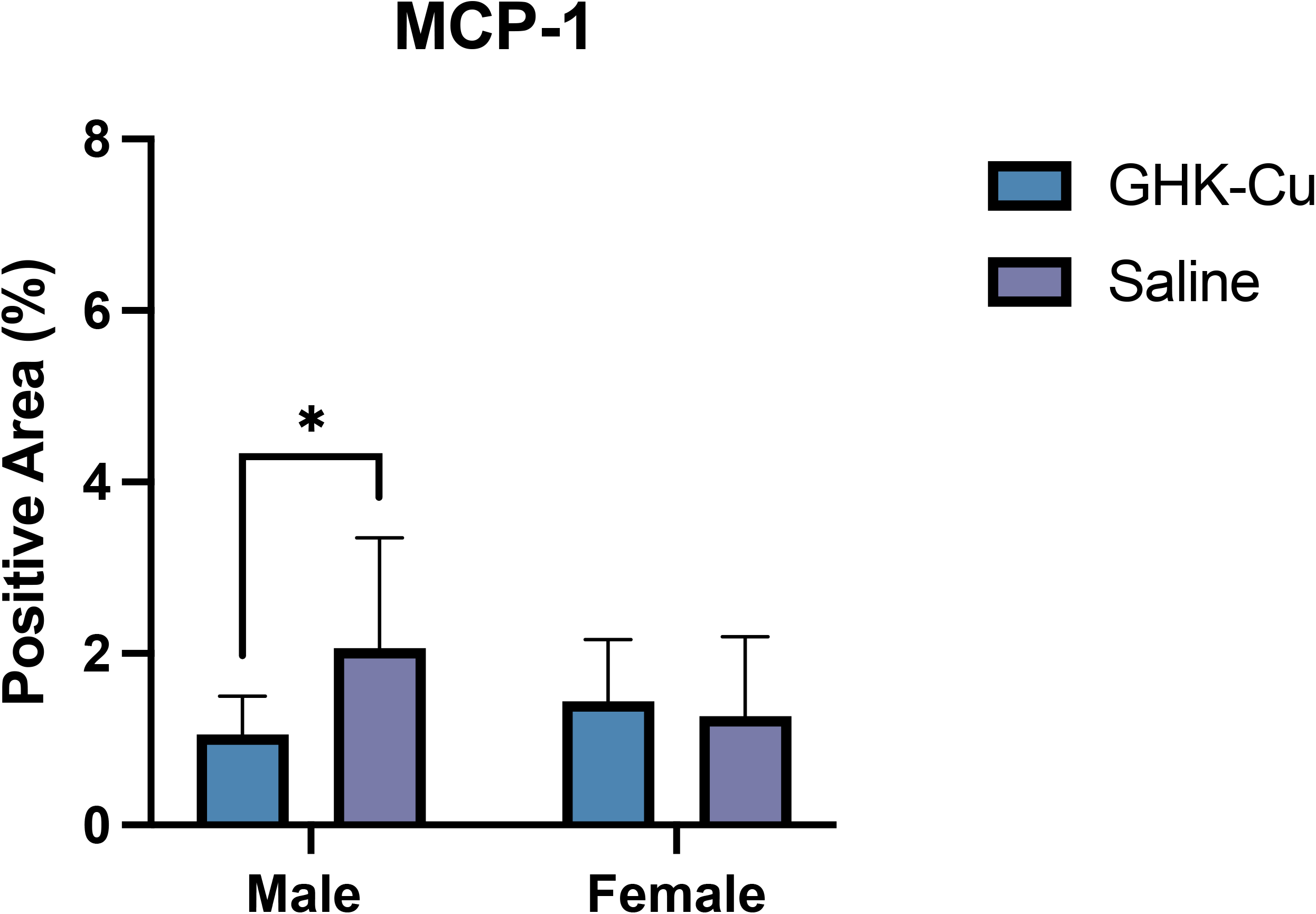
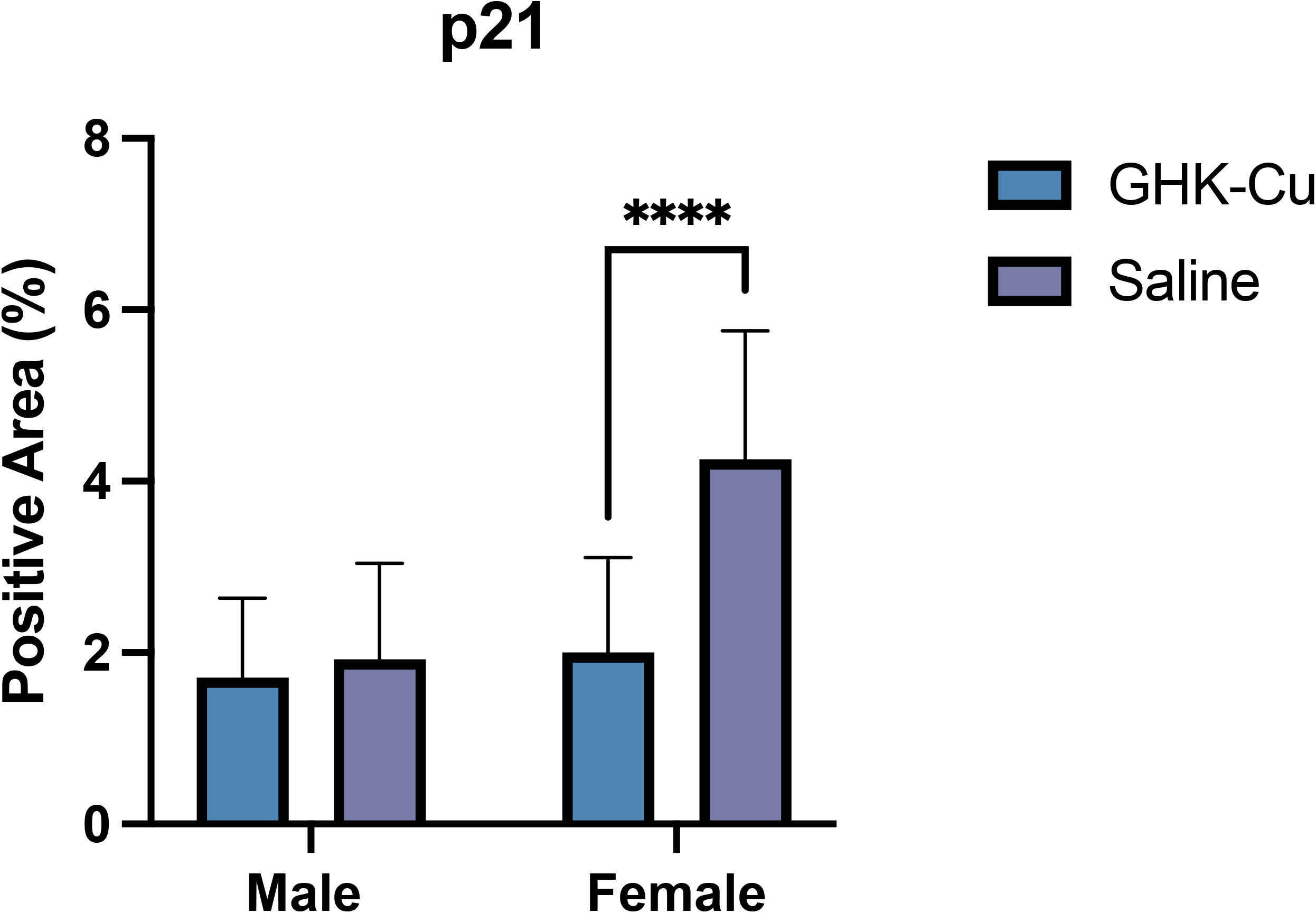
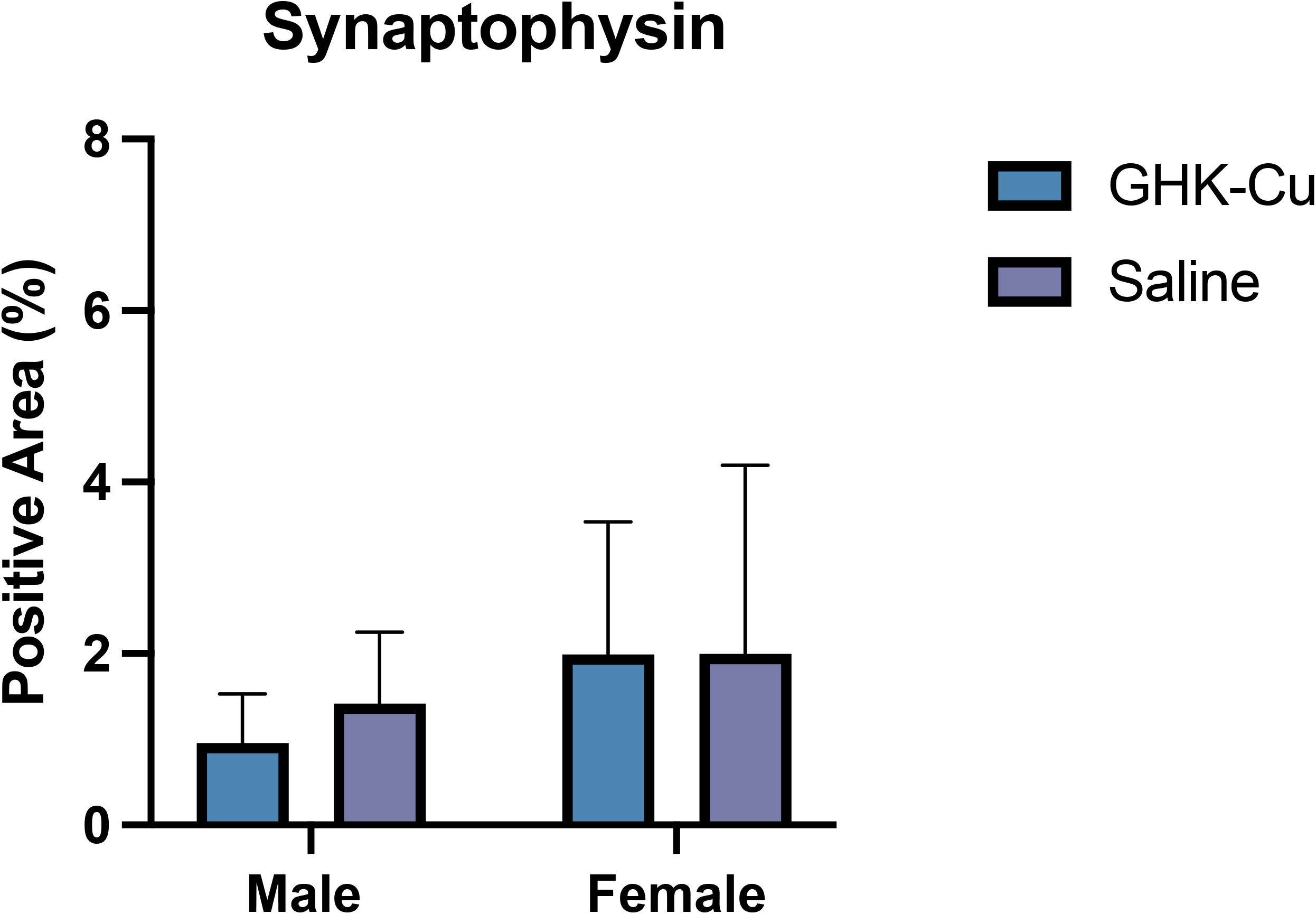
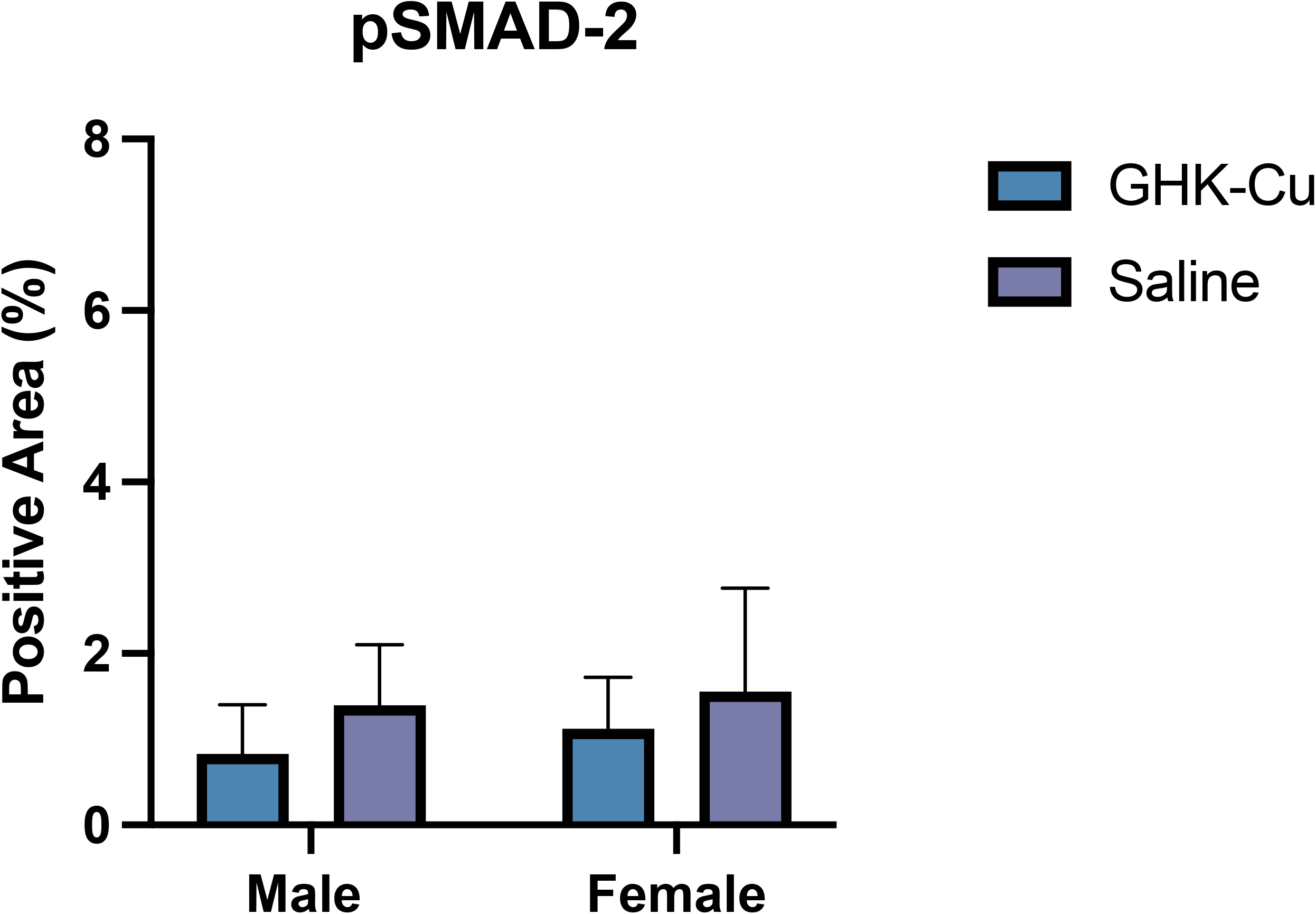
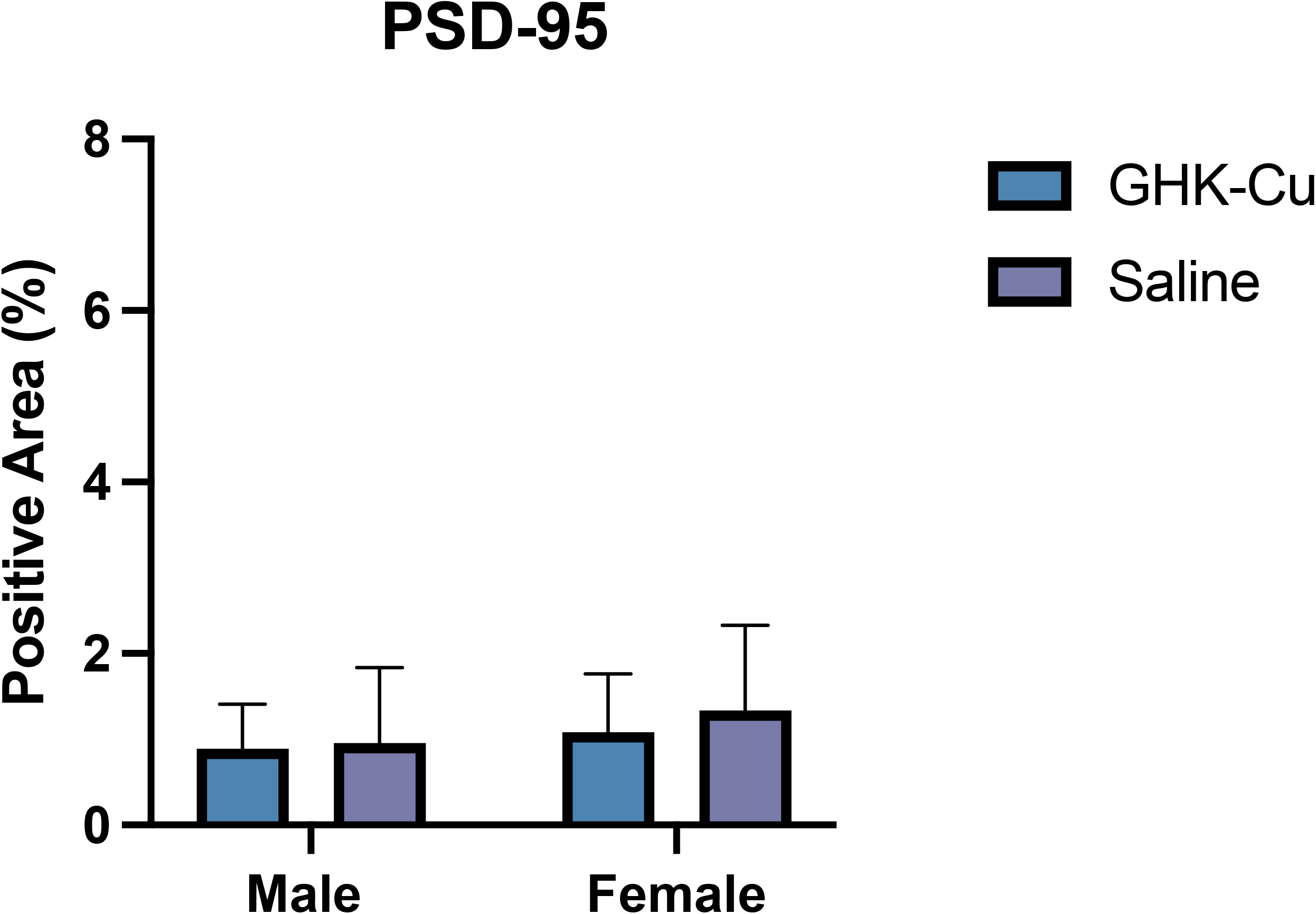
Positive area-staining by antibody in intraperitoneally-treated GHK-Cu mice of both sexes for (A) TGF-β, (B) GFAP), (C) MCP-1, (D) p21, (E) Synaptophysin, (F) pSMAD-2, and (G) PSD-95. \**P* < 0.05, \*\**P* < 0.01, \*\*\*\**P* < 0.0001.

### IN GHK-Cu induced sex-divergent but convergent suppression of growth, mitochondrial, and nutrient-sensing programs consistent with modulation of aging biology in the hippocampus

GSEA was performed to assess transcriptional pathway changes in male and female hippocampi following IN GHK-Cu treatment.

Pathways related to mitochondrial function and metabolic regulation showed prominent signals across both sexes (Table 1, Figure 4). Oxidative phosphorylation displayed negative normalized enrichment scores in males (NES -5.44, FDR < 0.0001) and females (NES - 4.20, FDR < 0.0001) (Table 1, Fig. 4A-B,G-H). MYC target genes also showed negative scores in both sexes, (Female: NES -4.31, FDR < 0.0001; Male: NES -2.81, FDR = 0.104). Reactive oxygen species signaling were negatively enriched in males (NES -3.41, FDR = 0.015) but not females (NES -2.03, FDR = 0.635). DNA repair pathways also had a negative score in males (NES -4.08, FDR < 0.0001) but not females (NES -2.66, FDR = 0.251). In females, nutrient-sensing pathways showed negative scores for PI3K-AKT-mTOR signaling (NES -3.15, FDR = 0.062) and mTORC1 signaling (NES -2.64, FDR = 0.229), whereas these pathways showed weaker signals in males (PI3K-AKT-mTOR NES -1.98, FDR = 0.673; MTORC1 NES -1.81, FDR = 0.805). Androgen response pathways were positively enriched in both sexes (Male: NES 3.23, FDR = 0.032; Female: NES 2.79, FDR = 0.058) (Supplementary Table S2, Figures S3-4). Additional metabolic pathways showed negative scores in males, including adipogenesis (NES -3.09, FDR = 0.050), fatty acid metabolism (NES -2.81, FDR = 0.092), cholesterol homeostasis (NES -2.80, FDR = 0.083), and xenobiotic metabolism (NES -3.05, FDR = 0.052). In females, xenobiotic metabolism (NES 3.01, FDR = 0.027) and bile acid metabolism (NES 3.10, FDR = 0.024) showed positive scores.

**Table 1.** GSEA Profiles of Core Longevity Pathways (Long-term Intranasal GHK-Cu).

| Pathway | Sex | Enrichment Score | NES | $P$-value | FDR | Regulation |
| --- | --- | --- | --- | --- | --- | --- |
| MYC Targets V1 | Male | -0.287 | -2.809 | 0.006 | 0.104 | ↓ |
|  | Female | -0.472 | -4.312 | <0.0001 | <0.0001 | ↓ |
| Oxidative Phosphorylation | Male | -0.548 | -5.438 | <0.0001 | <0.0001 | ↓ |
|  | Female | -0.451 | -4.203 | <0.0001 | <0.0001 | ↓ |
| PI3K-AKT-mTOR | Male | -0.219 | -1.983 | 0.482 | 0.673 | NS |
|  | Female | -0.318 | -3.150 | 0.013 | 0.062 | ↓ |
| MTORC1 | Male | -0.191 | -1.806 | 0.785 | 0.805 | NS |
|  | Female | -0.284 | -2.637 | 0.035 | 0.229 | ↓ |
| Reactive Oxygen Species | Male | -0.372 | -3.409 | 0.006 | 0.015 | ↓ |
|  | Female | -0.213 | -2.028 | 0.611 | 0.635 | NS |
| DNA Repair | Male | -0.456 | -4.076 | <0.0001 | <0.0001 | ↓ |
|  | Female | -0.259 | -2.663 | 0.122 | 0.251 | NS |

**Figure 4.**
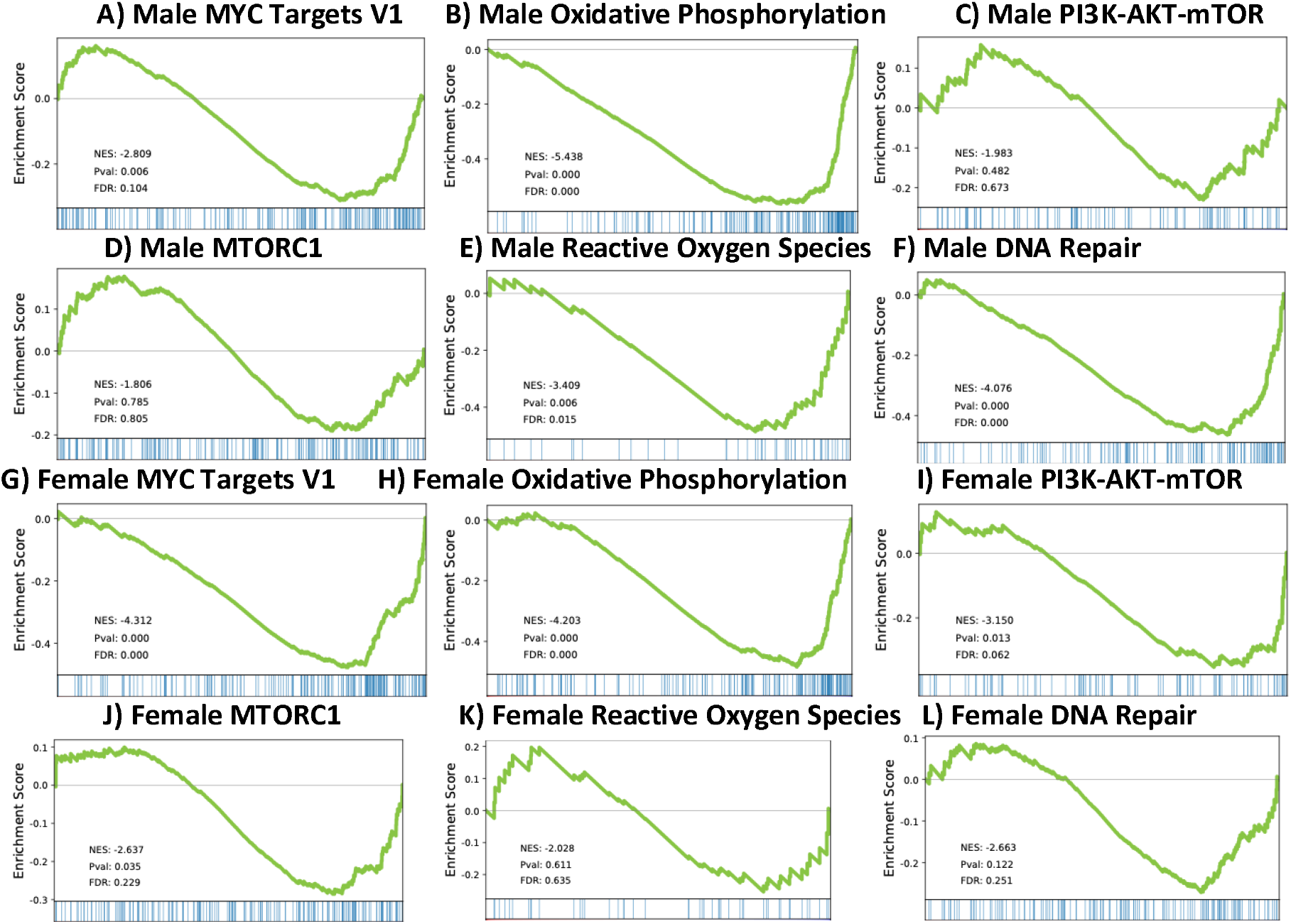
GSEA Graphs of Core Longevity Pathways (Long-Term Intranasal GHK-Cu).

Immune-associated pathways showed elevated signals in female hippocampi (Supplementary Table S1, Figure S2). Interferon α response (NES 3.55, FDR = 0.010), interferon γ response (NES 3.50, FDR = 0.007), and IL2-STAT5 signaling (NES 3.56, FDR = 0.018) were among the strongest positive scores. IL6-JAK-STAT3 signaling also showed a positive score (NES 2.78, FDR = 0.055), and coagulation pathways showed a similar pattern (NES 2.86, FDR = 0.047). These pathways did not show comparable signals in males (FDR’s > 0.1) (Supplementary Table S1, Figure S1).

Pathways related to cell cycle regulation and tissue remodeling also differed by sex (Supplementary Table S3, Figures S5-6). In males, G2M checkpoint (NES 3.17, FDR = 0.024), mitotic spindle (NES 3.41, FDR = 0.024), hedgehog signaling (NES 3.19, FDR = 0.028), and UV response DN (NES 4.93, FDR < 0.0001) showed positive scores, whereas UV response UP showed a negative score (NES -2.77, FDR = 0.090). In females, epithelial-mesenchymal transition (NES 3.43, FDR = 0.007), TGF-β signaling (NES 3.11, FDR = 0.028), and the p53 pathway (NES 2.73, FDR = 0.062) showed positive scores. Several additional hallmark pathways including angiogenesis, apoptosis, KRAS signaling, glycolysis, unfolded protein response, and multiple estrogen response pathways, did not show strong signals in either sex (FDR’s > 0.1) (Supplementary Tables S1-S4, Figures S1-S8).

### IP GHK-Cu induces sex-dependent activation of mitochondrial, growth, and stress-response pathways rather than the metabolic suppression observed with longer-term IN treatment

GSEA was performed to assess transcriptional pathway changes in male and female hippocampi following short-term IP GHK-Cu treatment.

Pathways related to mitochondrial metabolism and growth signaling showed positive normalized enrichment scores, particularly in females (Table 2, Figure 5). Oxidative phosphorylation showed a strong positive score in females (NES = 4.97, FDR < 0.001) and a weaker signal in males (NES = 2.94, FDR = 0.247). MYC target genes also showed positive scores in females (NES 4.34, FDR = 0.002) and males (NES = 2.47, FDR = 0.409). The reactive oxygen species pathway showed a positive score in females (NES = 3.40, FDR = 0.037) but not in males (NES = 2.47, FDR = 0.449). Additional pathways with positive scores in females included DNA repair (NES = 5.58, FDR < 0.001), E2F targets (NES = 3.79, FDR = 0.013), p53 signaling (NES = 3.75, FDR = 0.012), and mTORC1 signaling (NES = 3.00, FDR = 0.136) (Table 2, Figure 5). In males, these pathways showed weaker signals and higher FDR values (Table 2, Figure 5). Within metabolic pathways, androgen response showed a negative score in males (NES = -2.81, FDR = 0.129), while xenobiotic metabolism showed a positive score in females (NES = 2.97, FDR = 0.136) (Supplementary Table S6, Figures S11-12).

**Table 2.**
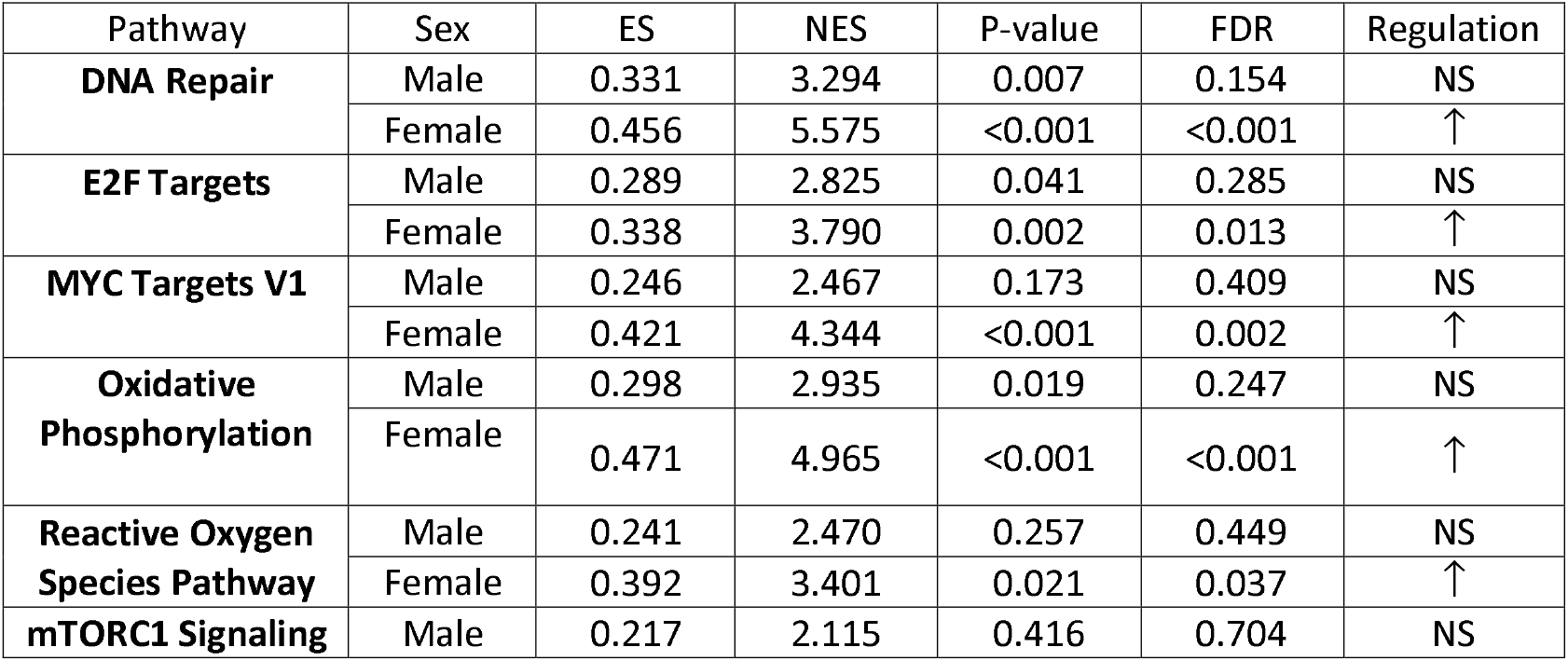
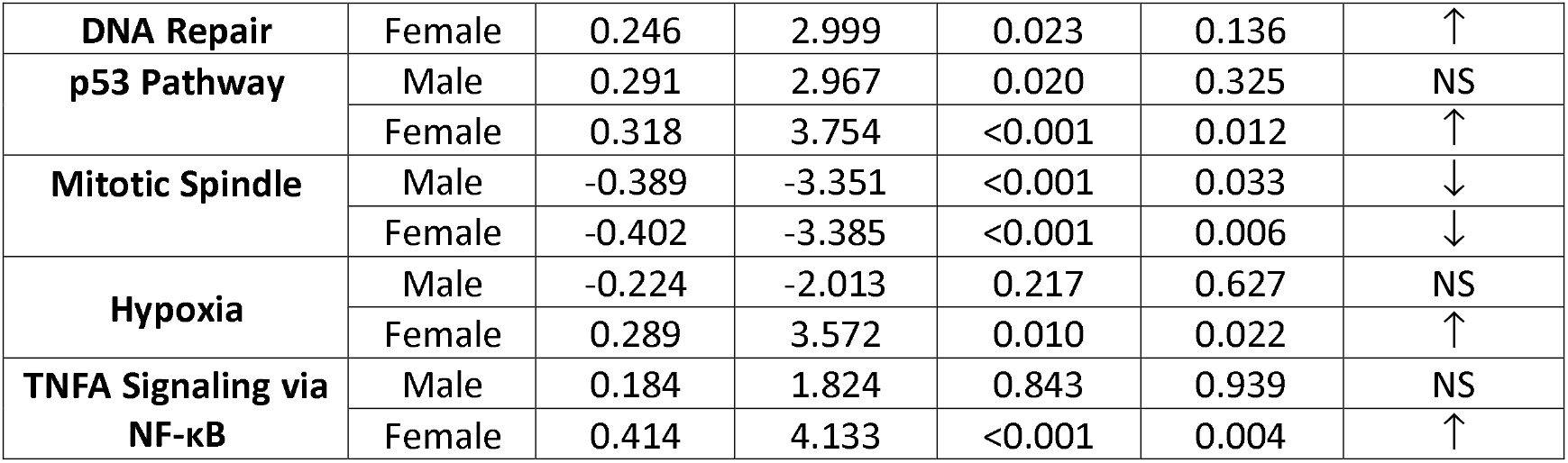
GSEA Profiles of Core Longevity Pathways (Short-Term Intraperitoneal GHK-Cu).

| Pathway | Sex | ES | NES | P-value | FDR | Regulation |
| --- | --- | --- | --- | --- | --- | --- |
| **DNA Repair** | Male | 0.331 | 3.294 | 0.007 | 0.154 | NS |
|  | Female | 0.456 | 5.575 | <0.001 | <0.001 | ↑ |
| **E2F Targets** | Male | 0.289 | 2.825 | 0.041 | 0.285 | NS |
|  | Female | 0.338 | 3.790 | 0.002 | 0.013 | ↑ |
| **MYC Targets V1** | Male | 0.246 | 2.467 | 0.173 | 0.409 | NS |
|  | Female | 0.421 | 4.344 | <0.001 | 0.002 | ↑ |
| **Oxidative Phosphorylation** | Male | 0.298 | 2.935 | 0.019 | 0.247 | NS |
|  | Female | 0.471 | 4.965 | <0.001 | <0.001 | ↑ |
| **Reactive Oxygen Species Pathway** | Male | 0.241 | 2.470 | 0.257 | 0.449 | NS |
|  | Female | 0.392 | 3.401 | 0.021 | 0.037 | ↑ |
| **mTORC1 Signaling** | Male | 0.217 | 2.115 | 0.416 | 0.704 | NS |
| **DNA Repair** | Female | 0.246 | 2.999 | 0.023 | 0.136 | ↑ |
| **p53 Pathway** | Male | 0.291 | 2.967 | 0.020 | 0.325 | NS |
|  | Female | 0.318 | 3.754 | <0.001 | 0.012 | ↑ |
| **Mitotic Spindle** | Male | -0.389 | -3.351 | <0.001 | 0.033 | ↓ |
|  | Female | -0.402 | -3.385 | <0.001 | 0.006 | ↓ |
| **Hypoxia** | Male | -0.224 | -2.013 | 0.217 | 0.627 | NS |
|  | Female | 0.289 | 3.572 | 0.010 | 0.022 | ↑ |
| **TNFA Signaling via NF-κB** | Male | 0.184 | 1.824 | 0.843 | 0.939 | NS |
|  | Female | 0.414 | 4.133 | <0.001 | 0.004 | ↑ |

**Figure 5.**
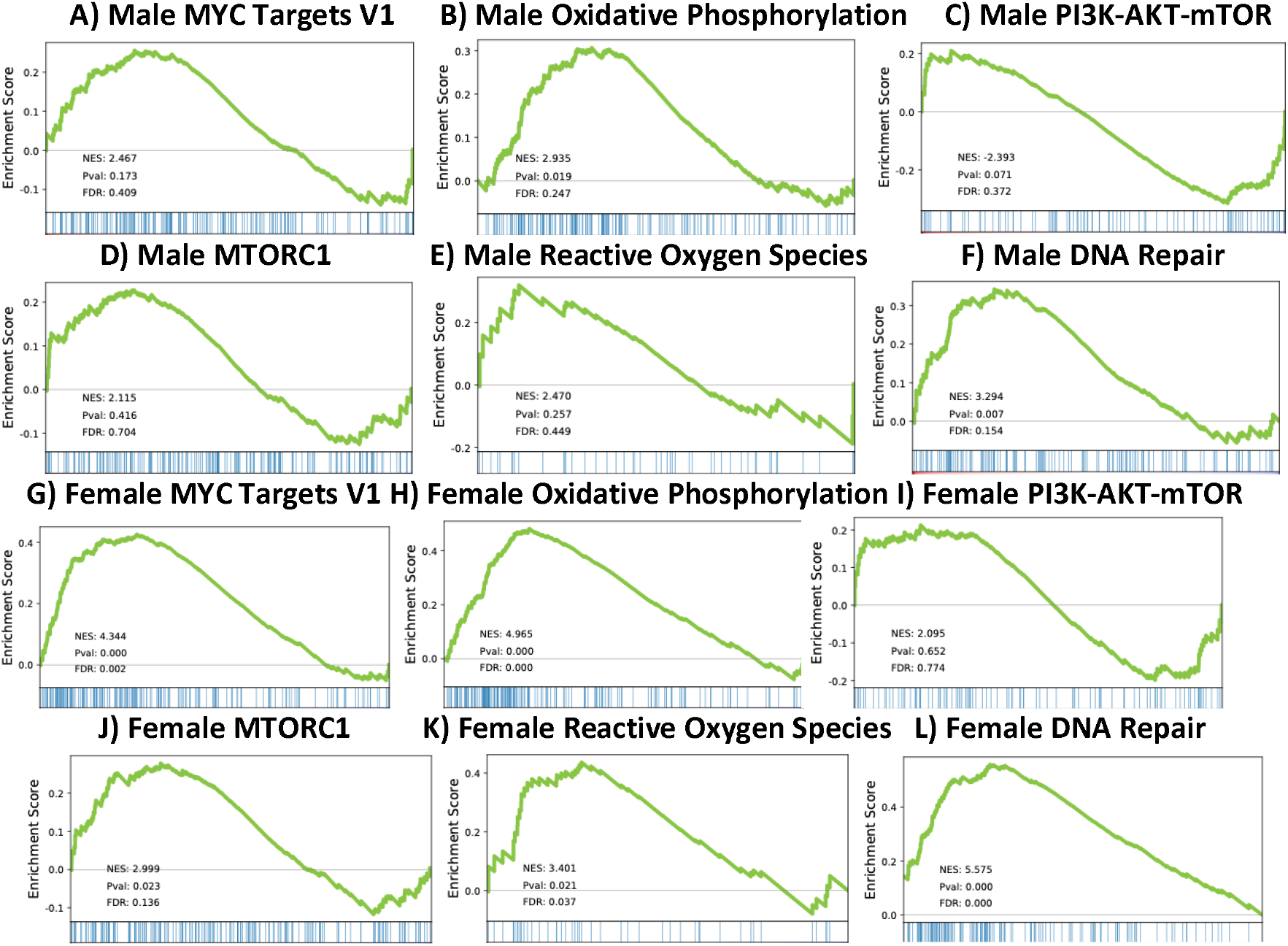
GSEA Graphs of Core Longevity Pathways (Short-Term Intraperitoneal GHK-Cu).

Immune-associated pathways were limited overall but showed elevated scores in females (Supplementary Table S5, Figures S9-10). TNFα signaling via NF-κB showed a strong positive score in females (NES = 4.13, FDR = 0.004), and hypoxia signaling also showed a positive score (NES = 3.57, FDR = 0.022). Other immune pathways, including interferon α response, interferon γ response, IL2-STAT5 signaling, IL6-JAK-STAT3 signaling, complement, inflammatory response, and coagulation, did not show strong signals in either sex (FDR’s > 0.1) (Supplementary Table S5, Figures S9-10).

Cell cycle and genome stability pathways showed additional sex-specific patterns (Supplementary Table S7, Figures S13-14). In females, DNA repair, E2F targets, and p53 signaling showed positive scores, whereas mitotic spindle pathways showed negative scores in both sexes (male NES = -3.35, FDR = 0.033; female NES = -3.39, FDR = 0.006). UV response DN pathways also showed negative scores in both sexes (male NES = -2.72, FDR = 0.141; female NES = -3.03, FDR = 0.024), while UV response UP showed a positive score in females (NES 3.22, FDR = 0.069). Additional pathways, including epithelial-mesenchymal transition, TGF-β signaling, KRAS signaling, and myogenesis, did not show strong signals (FDR’s > 0.1) (Supplementary Table S7, Figures S13-14).

Several additional hallmark pathways of aging, including estrogen response programs, angiogenesis, apoptosis, glycolysis, unfolded protein response, and cellular structure pathways, did not show strong signals in either sex (FDR’s > 0.1) (Supplementary Tables S5-8, Figures S9-16).

## Discussion

Both short-term IP and longer-term IN administration of GHK-Cu improved hippocampal-dependent escape learning in middle-aged mice, yet these effects arose from distinct transcriptional states. Short-term IP exposure engaged cellular repair and stress response pathways, including DNA repair, E2F signaling, p53 pathways, oxidative phosphorylation, and NF-κB-associated inflammation in females, features consistent with canonical responses to metabolic stress or tissue injury [4,24]. In contrast, longer-term IN delivery produced coordinated suppression of MYC targets, oxidative phosphorylation, and nutrient-sensing signaling pathways, including PI3K-AKT-mTOR signaling in females. MYC-dependent networks regulate ribosomal biogenesis, mitochondrial metabolism, and cellular growth, and their suppression is a feature of longevity-associated interventions [4,15]. Concurrent downregulation of oxidative phosphorylation suggests reduced mitochondrial demand and oxidative stress [25]. Together, these findings indicate that similar behavioral outcomes can emerge from divergent molecular programs, supporting the existence of multiple temporal modes of gerotherapeutic peptide action.

Sex-specific responses were evident across both paradigms. Short-term IP dosing elicited stronger transcriptional activation in females despite behavioral improvement being restricted to males, whereas longer-term IN treatment improved performance in both sexes despite divergent molecular responses. Females showed greater modulation of nutrient-sensing pathways, while males exhibited stronger suppression of oxidative stress-related programs, consistent with known sex-dependent differences in metabolic and neuroimmune regulation in aging [26-27]. Longer-term IN GHK-Cu further aligned with core hallmarks of aging biology, including a MYC-low transcriptional state and reduced mitochondrial metabolic load [4,25,28]. In females, activation of interferon- and cytokine-related pathways may reflect immune recalibration rather than generalized inflammation, consistent with emerging models of neuroinflammation as altered intercellular communication among neural cell types [5, 29-30]. Signals involving G2M checkpoint and mitotic spindle pathways should be interpreted cautiously, as they may reflect shifts in cellular composition or glial activation rather than neuronal proliferation [5].

The divergence between treatment paradigms appears primarily driven by exposure duration rather than route alone. IP administration produces rapid systemic exposure, whereas IN delivery enables sustained central access via olfactory pathways and bypass of the BBB [19]. Accordingly, short-term exposure captures transient repair responses, while prolonged delivery permits stable metabolic and transcriptional remodeling of aging neural circuits. These findings suggest distinct therapeutic roles, with short-term IP dosing potentially supporting transient performance enhancement and sustained IN delivery promoting longer-term remodeling of aging-associated pathways.

Several limitations should be considered. First, bulk RNA sequencing cannot resolve cell-type-specific transcriptional changes and may reflect shifts in cellular composition [31-32]. Single-cell or spatial transcriptomic approaches would provide greater mechanistic resolution [33-34]. Second, pharmacokinetic measurements of brain GHK-Cu concentrations were not performed, limiting interpretation of route-dependent exposure differences [19,35]. Without direct brain concentration measurements, it remains unclear whether the transcriptional differences observed between treatment paradigms reflect pharmacokinetic differences, exposure duration, or both. Third, transcriptomic profiling was conducted at a single time point, precluding evaluation of temporal dynamics in molecular responses. Aging-related pathways and stress-response programs often evolve over time, with early adaptive responses differing from longer-term metabolic or inflammatory remodeling [4]. Finally, this study did not include young reference cohorts, limiting the ability to determine whether GHK-Cu restores transcriptional profiles toward a youthful state or instead induces alternative compensatory programs in aged hippocampi. Future studies integrating longitudinal, pharmacokinetic, and cell-type-resolved approaches will be required to more precisely define the mechanisms underlying these effects.

In summary, GHK-Cu improves cognitive performance through distinct molecular modes via acute repair versus sustained metabolic suppression, highlighting exposure duration as a key determinant of gerotherapeutic peptide response.

## Supporting information

Publication License for Graphical Abstract

## Acknowledgments

This work was supported by the National Institute on Aging of the National Institutes of Health under award number R01 067193 (Ladiges, PI). The funder had no role in study design, data collection and analysis, decision to publish, or preparation of the manuscript.

## Availability of Data and Materials

The datasets generated during the current study are available from the corresponding author on reasonable request.

---

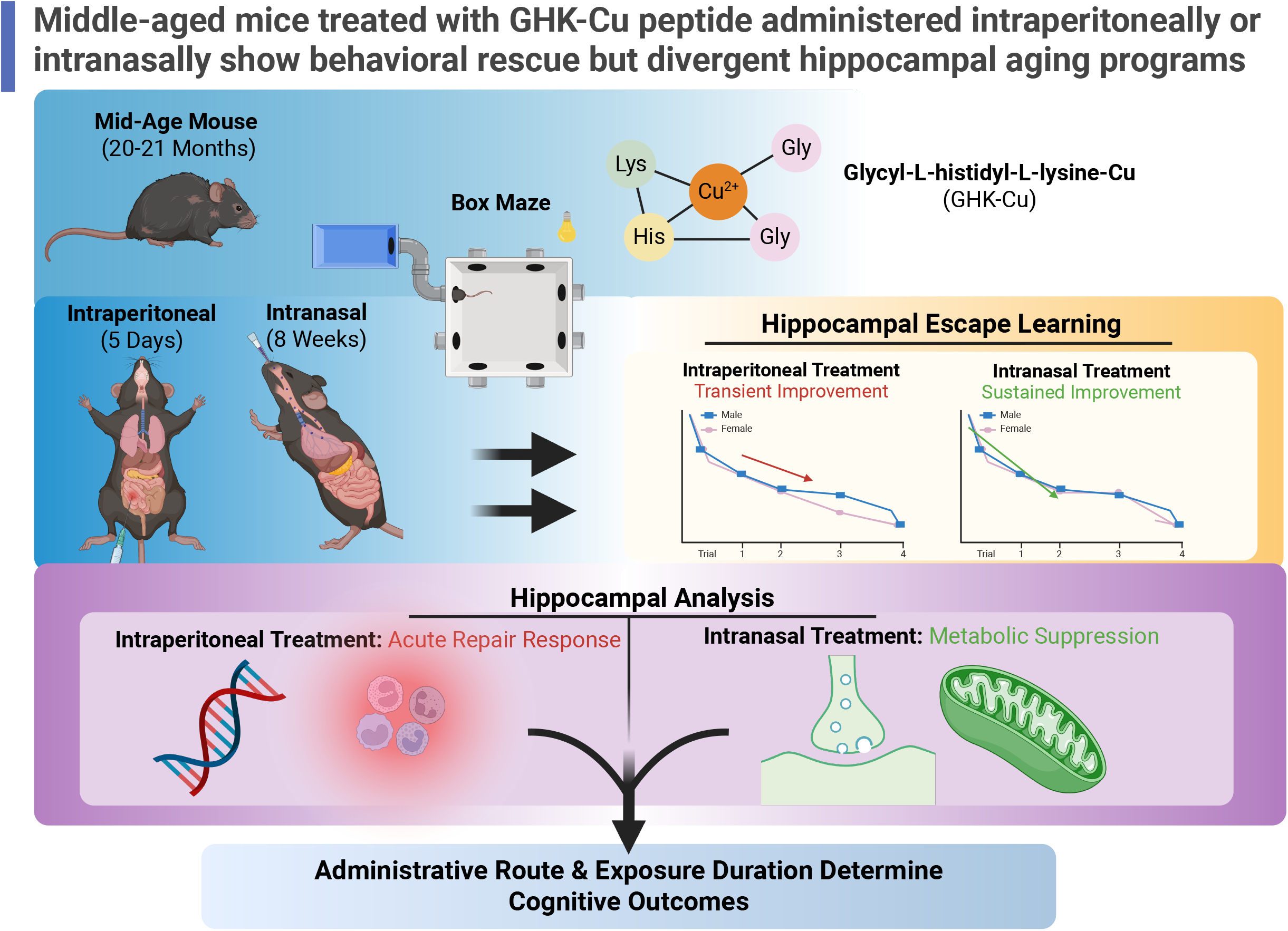

