## Supplementary material for "Middle-aged mice treated with GHK-Cu peptide administered intraperitoneally or intranasally show behavioral rescue but divergent hippocampal aging programs": Supp. Table 1-2 & Fig. 1-4

### **Supplementary Table S1. Immune & Inflammatory Pathways GSEA Profiles (Intranasal GHK-Cu).**

| **Pathway** | **Sex** | **ES** | **NES** | **P-value** | **FDR** | **Regulation** |
| --- | --- | --- | --- | --- | --- | --- |
| **Interferon α Response** | Male | -0.292 | -2.429 | 0.164 | 0.270 | NS |
|  | Female | 0.462 | 3.546 | <0.0001 | 0.010 | ↑ |
| **Interferon γ Response** | Male | -0.257 | -2.344 | 0.100 | 0.279 | NS |
|  | Female | 0.401 | 3.499 | <0.0001 | 0.007 | ↑ |
| **IL2-STAT5 Signaling** | Male | 0.186 | 1.845 | 0.724 | 0.858 | NS |
|  | Female | 0.412 | 3.562 | <0.0001 | 0.018 | ↑ |
| **IL6-JAK-STAT3 Signaling** | Male | -0.231 | -2.149 | 0.237 | 0.446 | NS |
|  | Female | 0.347 | 2.784 | 0.010 | 0.055 | ↑ |
| **Complement** | Male | -0.213 | -1.871 | 0.564 | 0.800 | NS |
|  | Female | 0.286 | 2.540 | 0.031 | 0.128 | ↑ |
| **Inflammatory Response** | Male | -0.171 | -1.563 | 0.947 | 0.899 | NS |
|  | Female | 0.287 | 2.422 | 0.031 | 0.178 | ↑ |
| **Coagulation** | Male | -0.314 | -2.831 | 0.020 | 0.110 | ↓ |
|  | Female | 0.342 | 2.864 | 0.006 | 0.047 | ↑ |
| **TNF-α Signaling via NFκB** | Male | 0.182 | 1.686 | 0.887 | 1.000 | NS |
|  | Female | 0.226 | 2.140 | 0.342 | 0.333 | NS |
| **Allograft Rejection** | Male | -0.247 | -2.222 | 0.222 | 0.374 | NS |
|  | Female | 0.222 | 1.909 | 0.504 | 0.543 | NS |
| **Angiogenesis** | Male | -0.118 | -1.609 | 0.737 | 0.893 | NS |
|  | Female | 0.366 | 2.661 | 0.106 | 0.079 | NS |
| **Hypoxia** | Male | 0.153 | 1.520 | 0.972 | 1.000 | NS |
|  | Female | 0.262 | 2.398 | 0.093 | 0.184 | NS |
| **NOTCH Signaling** | Male | 0.176 | 1.425 | 0.931 | 1.000 | NS |
|  | Female | 0.241 | 2.242 | 0.302 | 0.268 | NS |
| **WNT β-catenin signaling** | Male | 0.262 | 2.123 | 0.379 | 0.625 | NS |
|  | Female | 0.281 | 2.526 | 0.138 | 0.126 | NS |
| **Apoptosis** | Male | -0.204 | -1.823 | 0.584 | 0.812 | NS |
|  | Female | 0.203 | 1.778 | 0.678 | 0.665 | NS |

**Supplemental Figure S1.** **Male Immune & Inflammatory Pathways GSEA Graphs (Intranasal GHK-Cu).**

**A)** **Interferon α Response	B) Interferon γ Response	C) IL2-STAT5 Signaling**

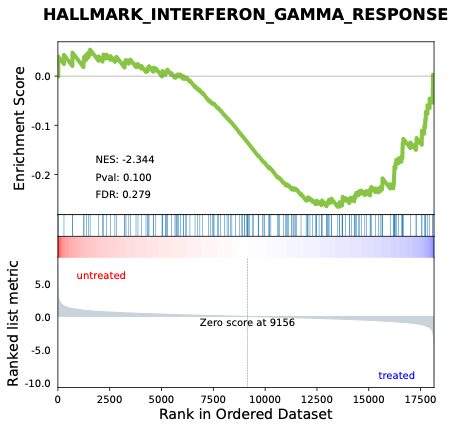

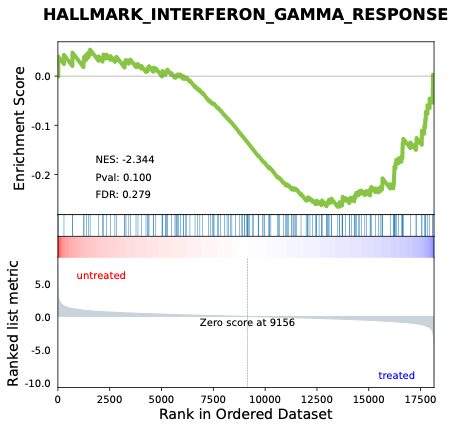

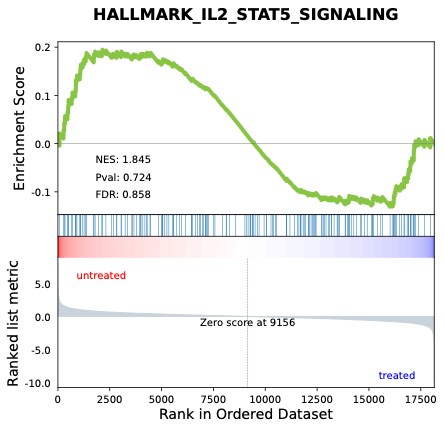

**D)** **IL6-JAK-STAT3 Signaling	E) Complement	F) Inflammatory Response**

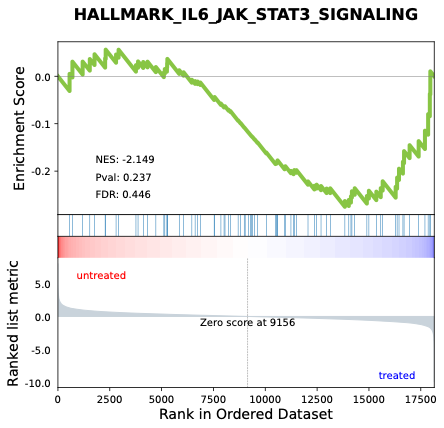

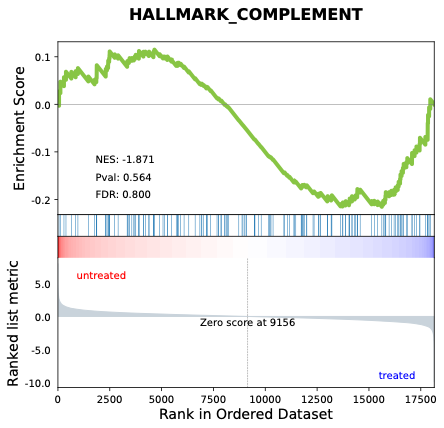

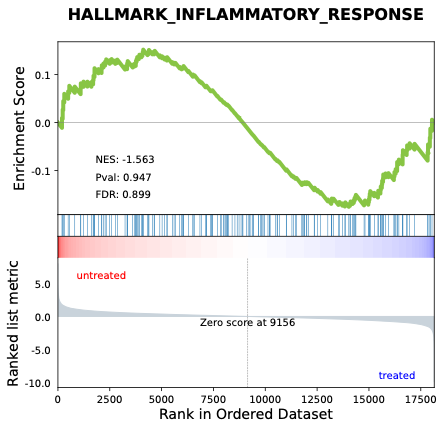

**G) Coagulation	H) TNF-α Signaling via NFκB	I) Allograft Rejection**

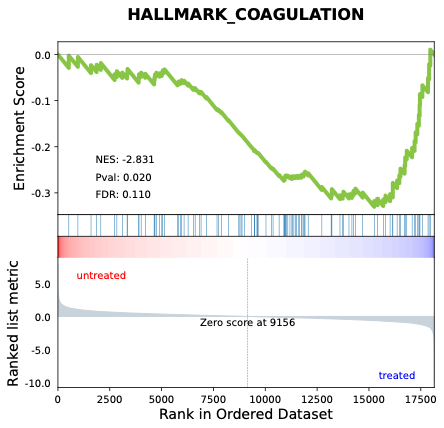

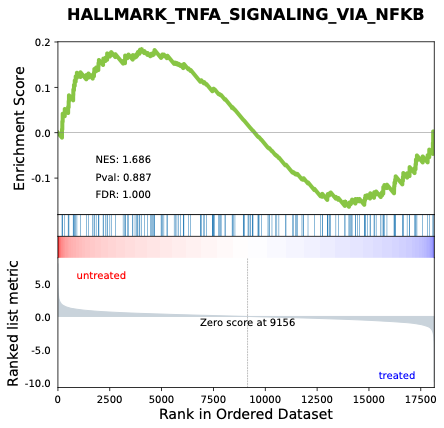

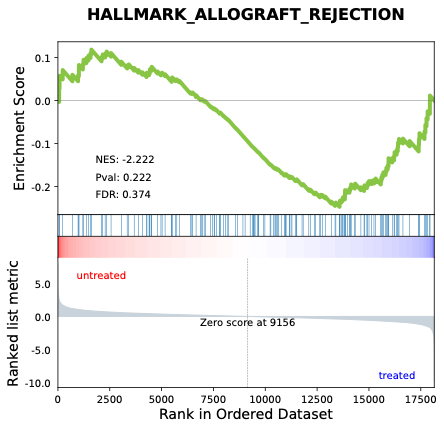

**J) Angiogenesis	K) Hypoxia	L) NOTCH Signaling**

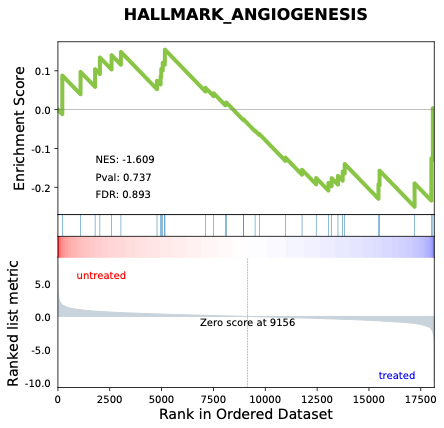

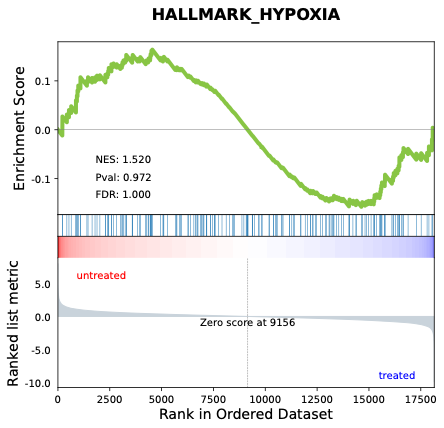

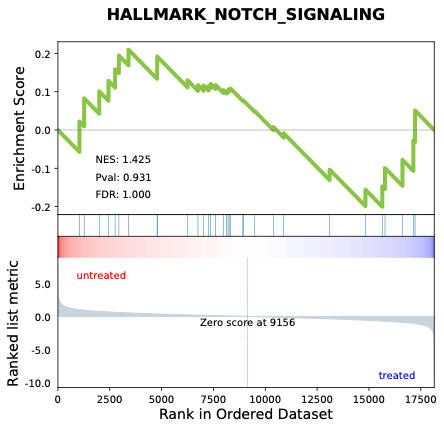

**M) WNT β-catenin signaling	N) Apoptosis**

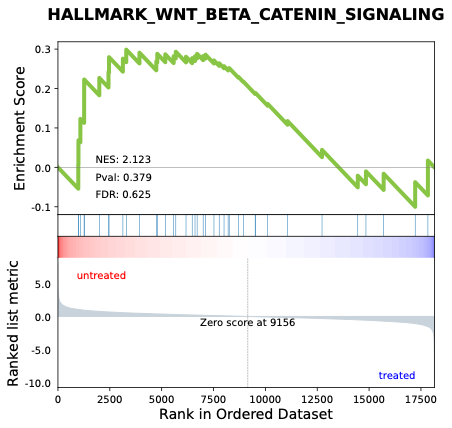

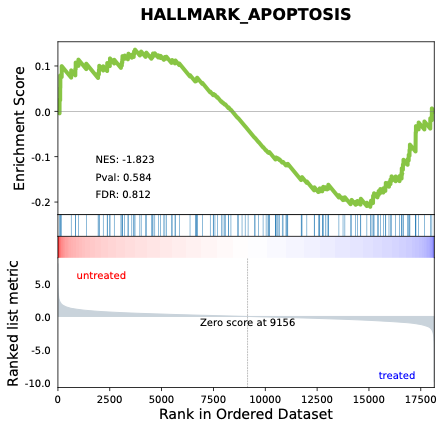

**Supplemental Figure S2.** Female **Immune & Inflammatory Pathways GSEA Graphs (Intranasal GHK-Cu).**

**A)** **Interferon α Response	B) Interferon γ Response	C) IL2-STAT5 Signaling**

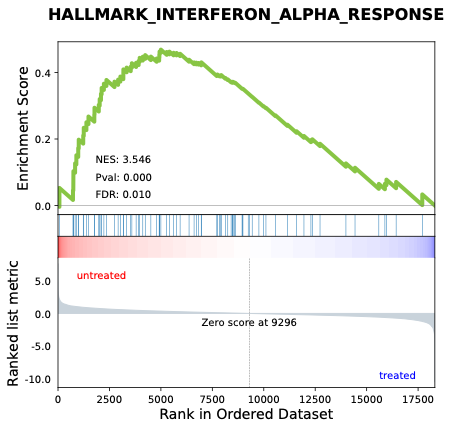

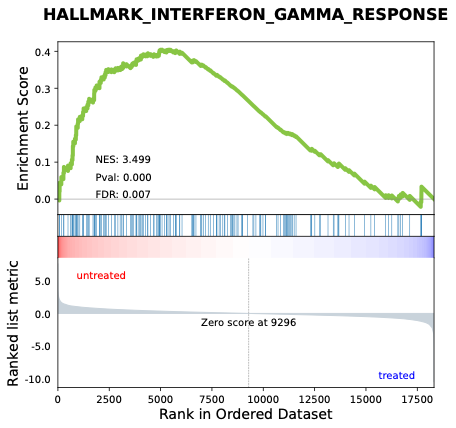

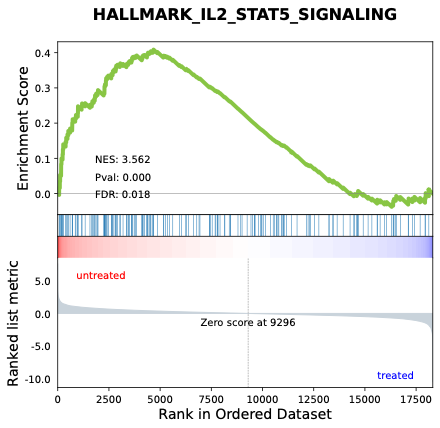

**D)** **IL6-JAK-STAT3 Signaling	E) Complement	F) Inflammatory Response**
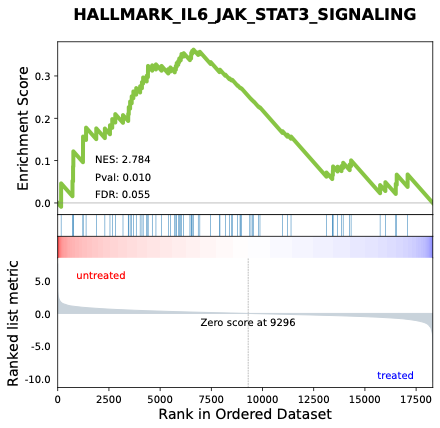

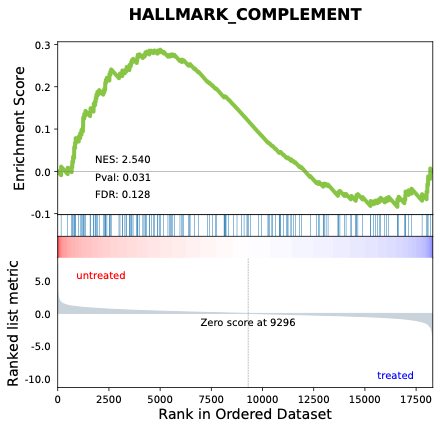

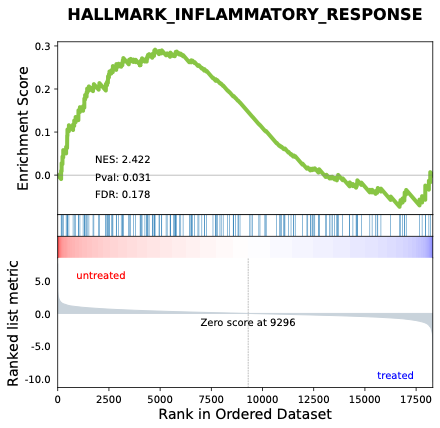

**G) Coagulation	H) TNF-α Signaling via NFκB	I) Allograft Rejection**
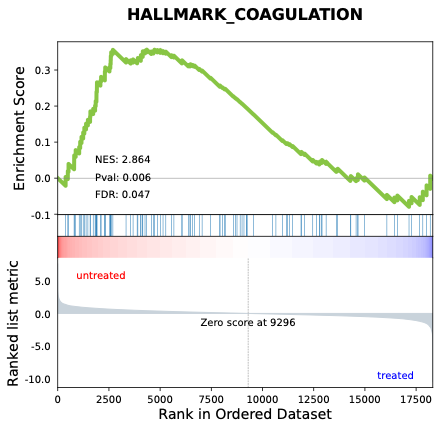

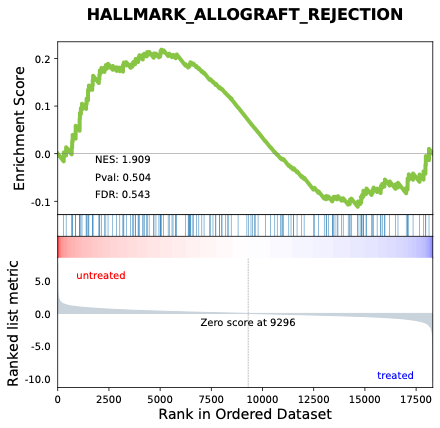

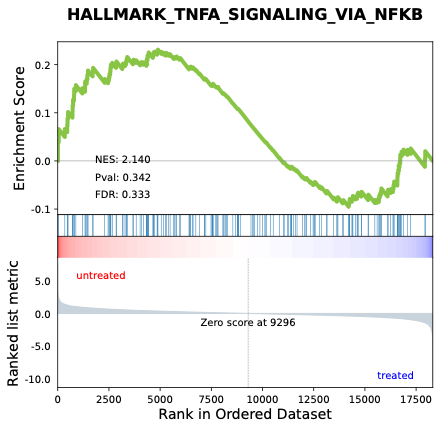

**J) Angiogenesis	K) Hypoxia	L) NOTCH Signaling**

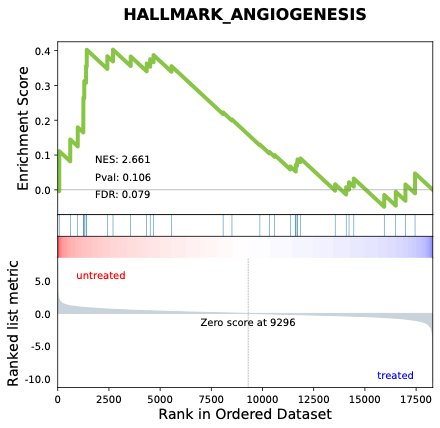

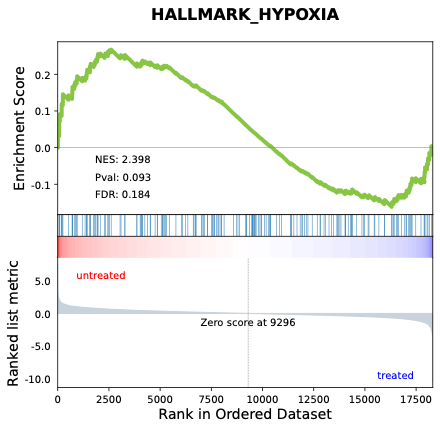

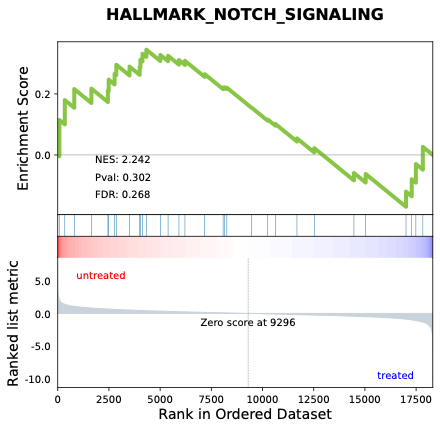

**M) WNT β-catenin signaling	N) Apoptosis**

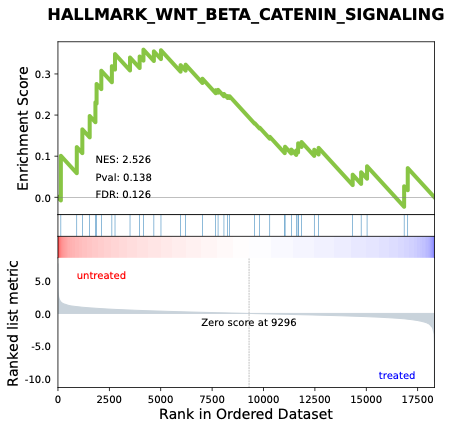

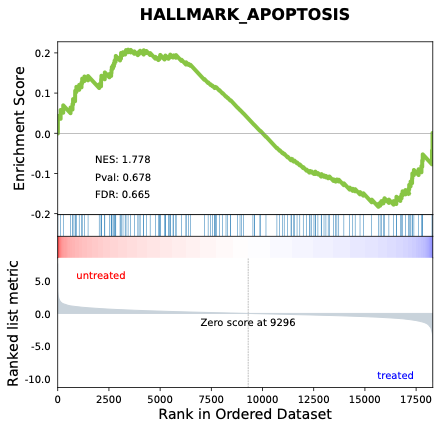

### **Supplementary Table S2. Metabolic & Detoxification Pathways GSEA Profiles (Intranasal GHK-Cu).**

| **Pathway** | **Sex** | **Enrichment Score** | **NES** | ***P*-value** | **FDR** | **Regulation** |
| --- | --- | --- | --- | --- | --- | --- |
| **Androgen Response** | Male | 0.361 | 3.234 | 0.002 | 0.032 | ↑ |
|  | Female | 0.318 | 2.789 | 0.023 | 0.058 | ↑ |
| **Oxidative Phosphorylation** | Male | -0.548 | -5.438 | <0.0001 | <0.0001 | ↓ |
|  | Female | -0.451 | -4.203 | <0.0001 | <0.0001 | ↓ |
| **Reactive Oxygen Species** | Male | -0.372 | -3.409 | 0.006 | 0.015 | ↓ |
|  | Female | -0.213 | -2.028 | 0.611 | 0.635 | NS |
| **MYC Targets V1** | Male | -0.287 | -2.809 | 0.006 | 0.104 | ↓ |
|  | Female | -0.472 | -4.312 | <0.0001 | <0.0001 | ↓ |
| **PI3K-AKT-mTOR Signaling** | Male | -0.219 | -1.983 | 0.482 | 0.673 | NS |
|  | Female | -0.318 | -3.150 | 0.013 | 0.062 | ↓ |
| **MTORC1 Signaling** | Male | -0.191 | -1.806 | 0.785 | 0.805 | NS |
|  | Female | -0.284 | -2.637 | 0.035 | 0.229 | ↓ |
| **Adipogenesis** | Male | -0.327 | -3.087 | 0.002 | 0.050 | ↓ |
|  | Female | 0.223 | 1.961 | 0.321 | 0.491 | NS |
| **Fatty Acid Metabolism** | Male | -0.329 | -2.806 | 0.017 | 0.092 | ↓ |
|  | Female | 0.247 | 2.073 | 0.266 | 0.398 | NS |
| **Cholesterol Homeostasis** | Male | -0.341 | -2.800 | 0.047 | 0.083 | ↓ |
|  | Female | -0.219 | -1.857 | 0.673 | 0.761 | NS |
| **Xenobiotic Metabolism** | Male | -0.318 | -3.047 | 0.006 | 0.052 | ↓ |
|  | Female | 0.334 | 3.012 | <0.0001 | 0.027 | ↑ |
| **Bile Acid Metabolism** | Male | -0.289 | -2.644 | 0.078 | 0.139 | NS |
|  | Female | 0.398 | 3.102 | 0.002 | 0.024 | ↑ |
| **Glycolysis** | Male | -0.266 | -2.487 | 0.108 | 0.235 | NS |
|  | Female | 0.231 | 2.166 | 0.269 | 0.333 | NS |
| **Heme Metabolism** | Male | -0.148 | -1.623 | 0.899 | 0.910 | NS |
|  | Female | 0.224 | 1.985 | 0.377 | 0.478 | NS |
| **Peroxisome** | Male | -0.284 | -2.392 | 0.161 | 0.251 | NS |
|  | Female | 0.257 | 2.012 | 0.304 | 0.459 | NS |
| **Unfolded Protein Response** | Male | 0.229 | 1.941 | 0.565 | 0.897 | NS |
|  | Female | -0.231 | -2.234 | 0.353 | 0.544 | NS |
| **Protein Secretion** | Male | 0.296 | 2.520 | 0.115 | 0.282 | NS |
|  | Female | -0.243 | -2.284 | 0.376 | 0.611 | NS |
| **Pancreatic β-cell signaling** | Male | -0.248 | -1.791 | 0.525 | 0.797 | NS |
|  | Female | -0.258 | -2.124 | 0.335 | 0.656 | NS |

**Supplemental Figure S3.** GSEA Graphs for Male **Metabolic and Detoxification Pathways** **(Intranasal GHK-Cu).**

**A) Androgen Response	B) Oxidative Phosphorylation E) Reactive Oxygen Species**

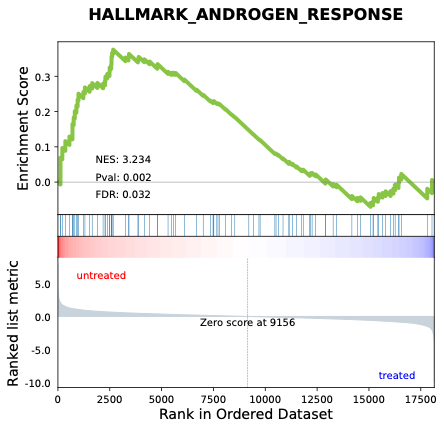

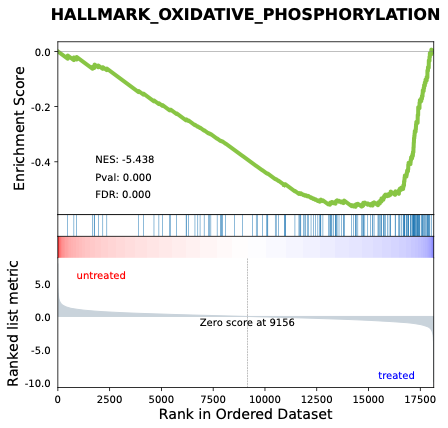

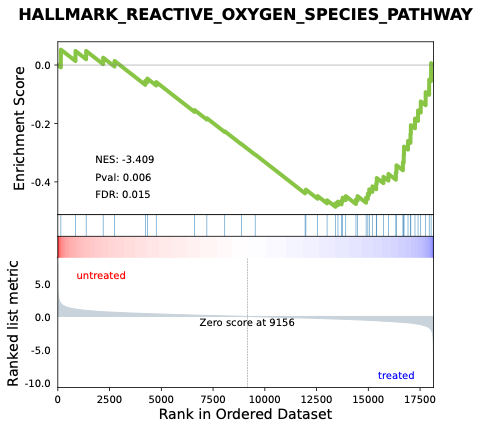

**D) MYC Targets V1	E) PI3K-AKT-mTOR Signaling	F) MTORC1 Signaling**

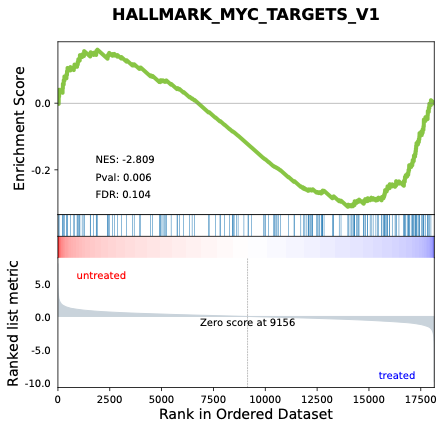

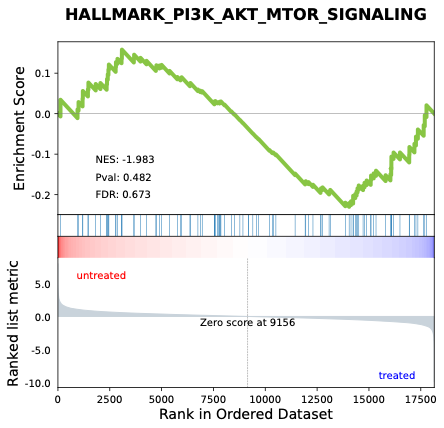

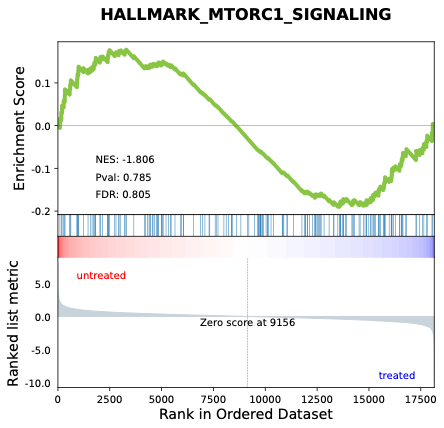

**G) Adipogenesis	H) Fatty Acid Metabolism	I) Cholesterol Homeostasis**
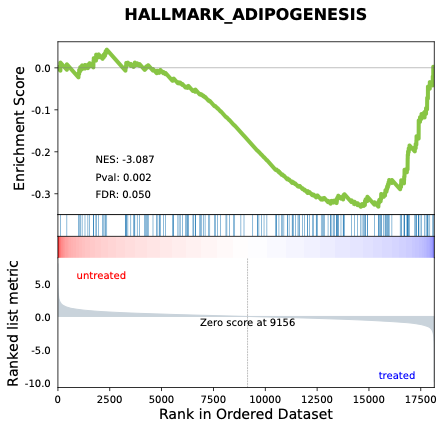

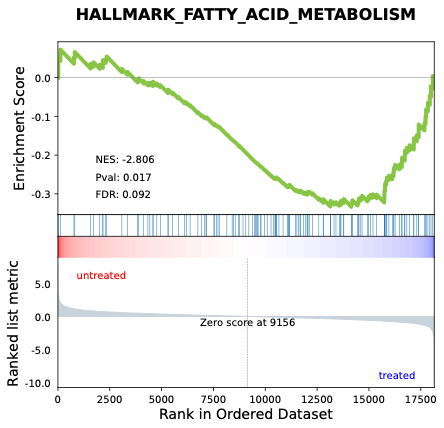

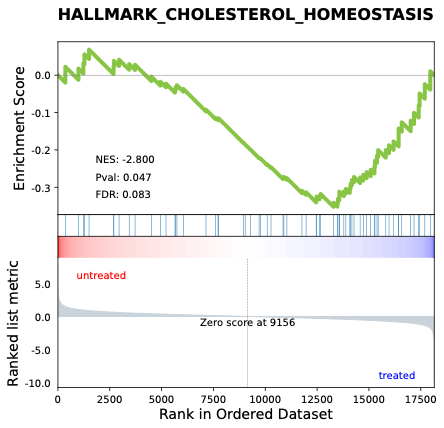

**J) Xenobiotic Metabolism	K) Bile Acid Metabolism	L) Glycolysis**

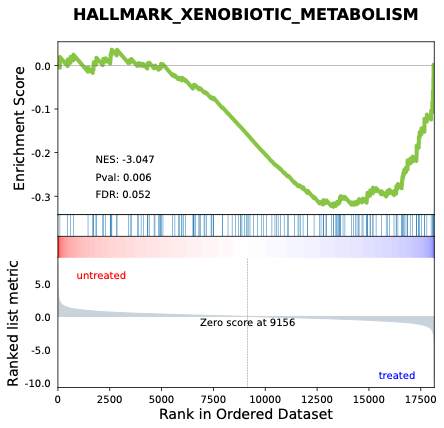

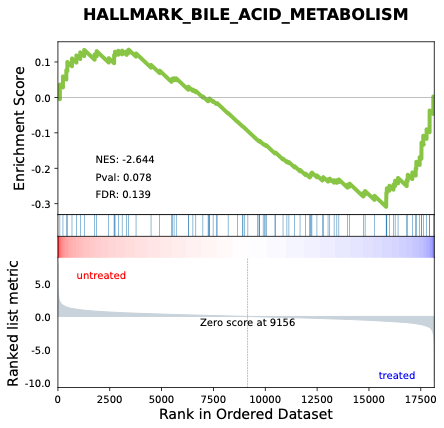

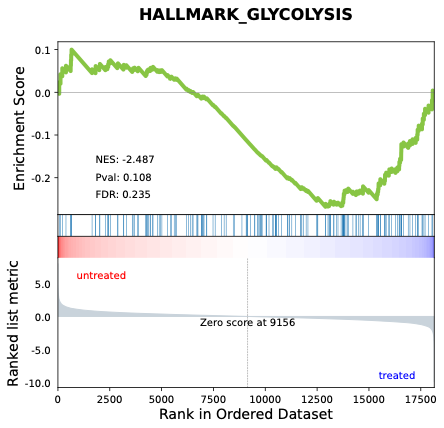

**M) Heme Metabolism	N) Peroxisome	O) Unfolded Protein Response**

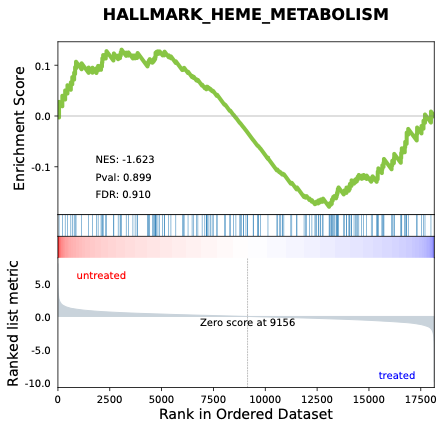

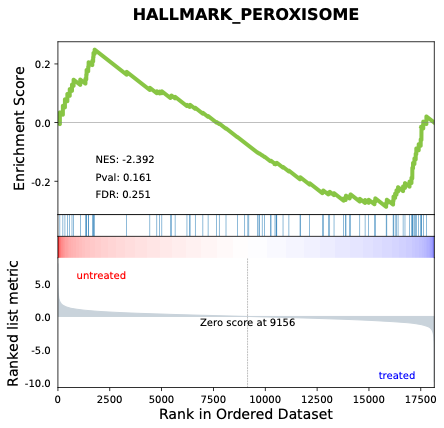

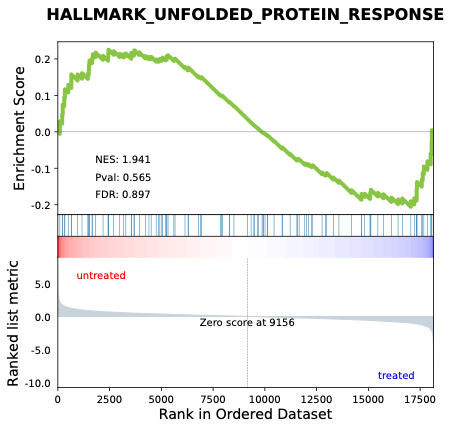

**P) Protein Secretion	Q)** **Pancreatic β-cell signaling**

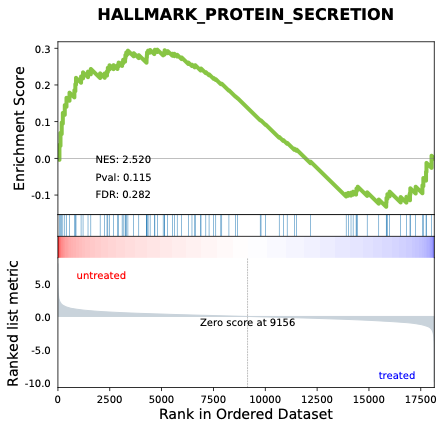

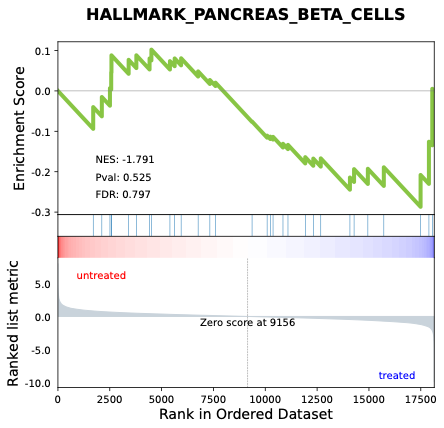

**Supplemental Figure S4.** GSEA Graphs for Female **Metabolic and Detoxification Pathways** **(Intranasal GHK-Cu).**

**A) Androgen Response	B) Oxidative Phosphorylation	C) Reactive Oxygen Species**

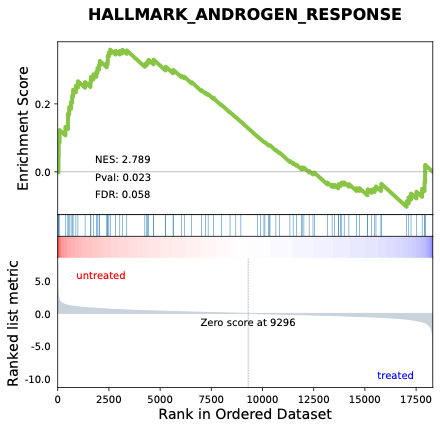

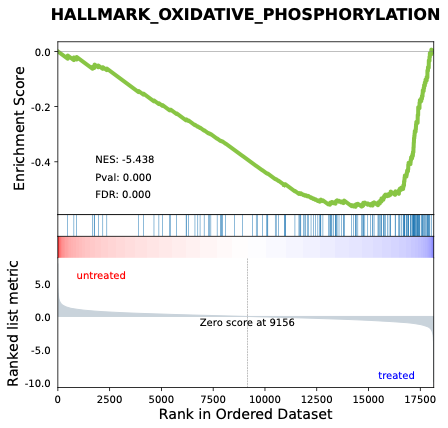

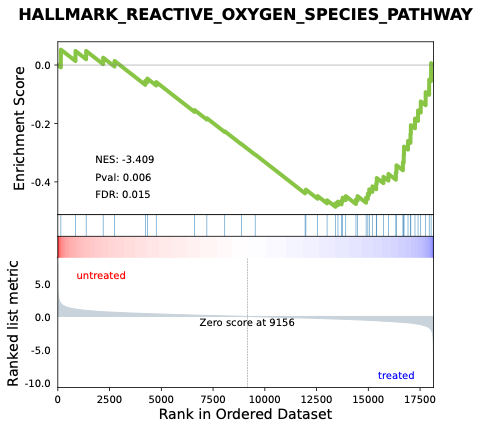

**D) MYC Targets V1	E) PI3K-AKT-mTOR Signaling	F) MTORC1 Signaling**

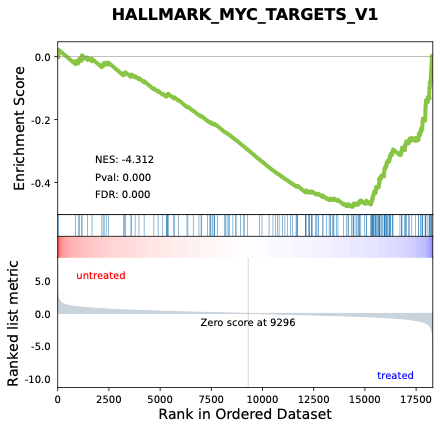

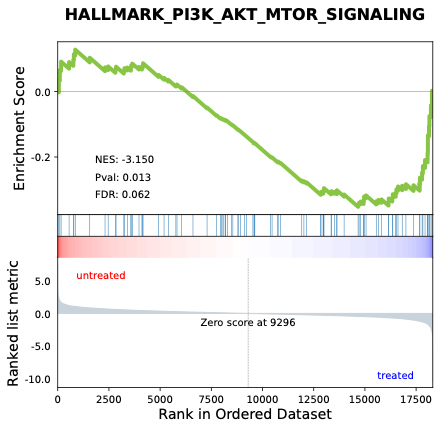

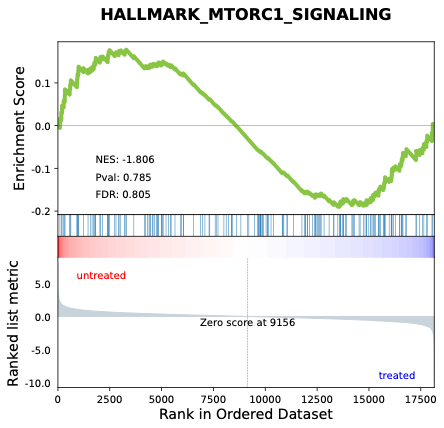

**G) Adipogenesis	H) Fatty Acid Metabolism	I) Cholesterol Homeostasis**

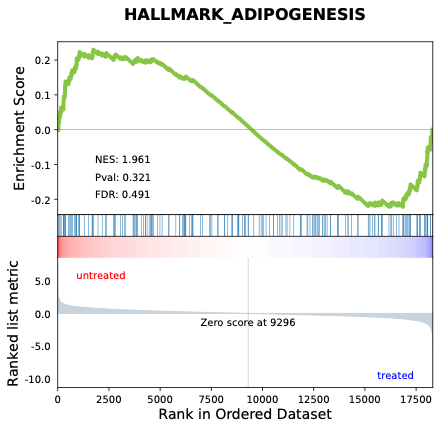

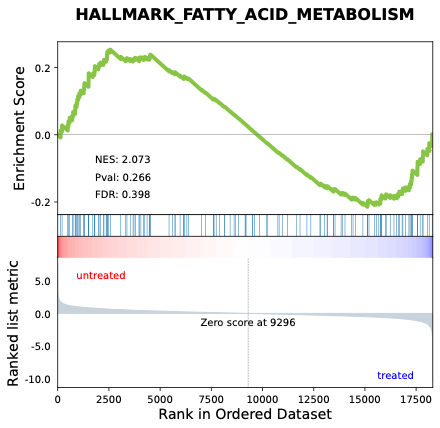

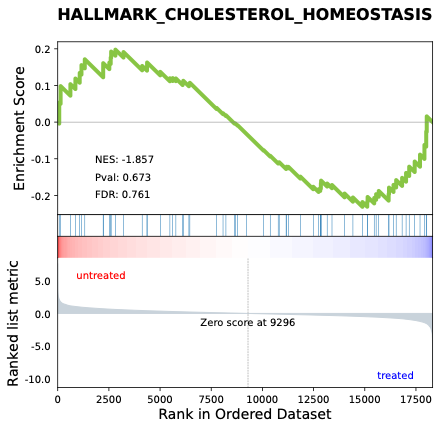

**J) Xenobiotic Metabolism	K) Bile Acid Metabolism	L) Glycolysis**

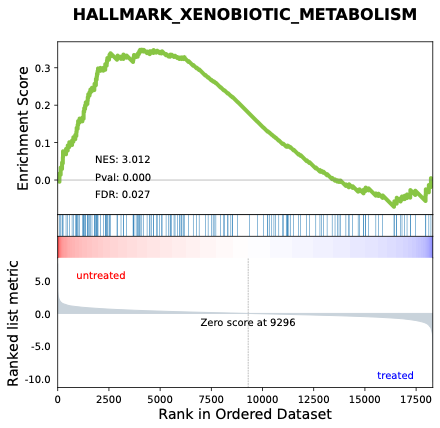

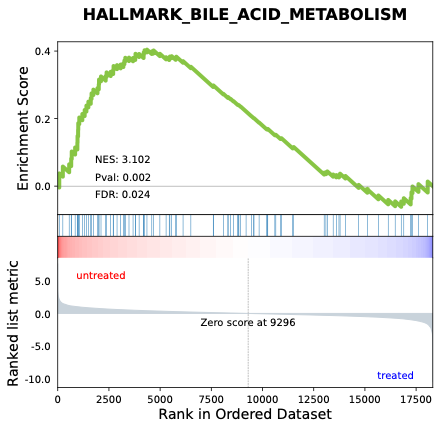

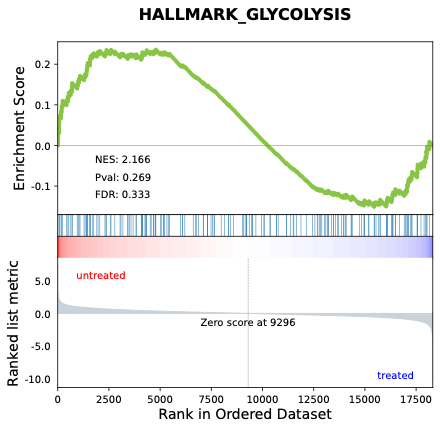

**M) Heme Metabolism	N) Peroxisome	O) Unfolded Protein Response**

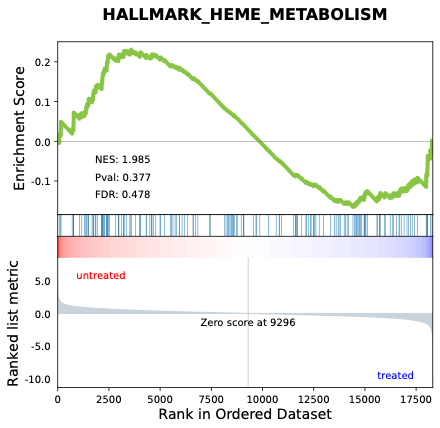

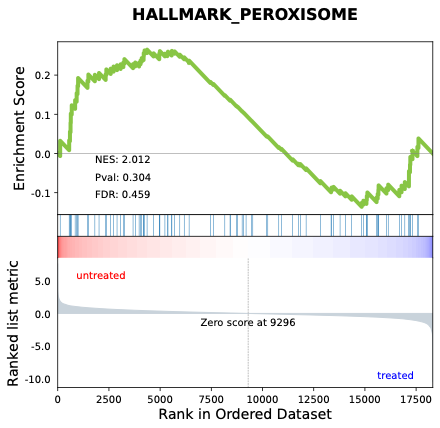

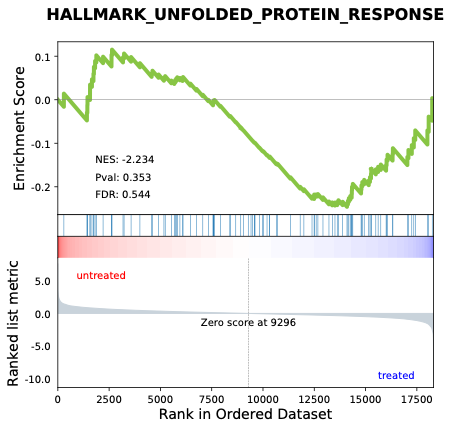

**P) Protein Secretion	Q) Pancreatic β-cell signaling**

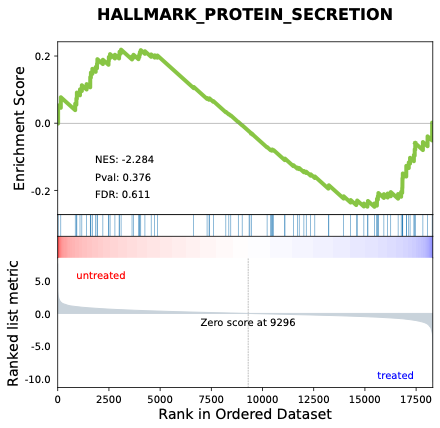

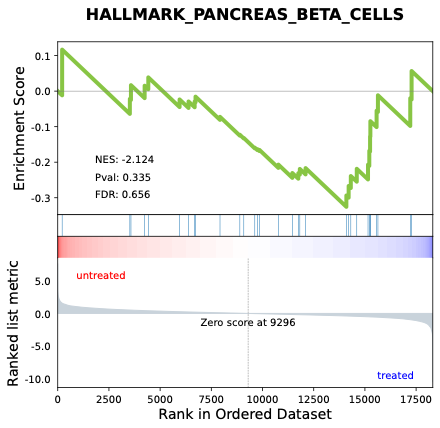
