## Supplementary material for "Middle-aged mice treated with GHK-Cu peptide administered intraperitoneally or intranasally show behavioral rescue but divergent hippocampal aging programs": Supp. Table 3-4 & Fig. 5-8

### **Supplementary Table S3. Cell Cycle, DNA Damage, and Tissue Remodeling Pathways GSEA Profiles (Intranasal GHK-Cu).**

| **Pathway** | **Sex** | **Enrichment Score** | **NES** | ***P*-value** | **FDR** | **Regulation** |
| --- | --- | --- | --- | --- | --- | --- |
| **DNA Repair** | Male | -0.456 | -4.076 | <0.0001 | <0.0001 | ↓ |
|  | Female | -0.259 | -2.663 | 0.122 | 0.251 | NS |
| **G2M Checkpoint** | Male | 0.338 | 3.170 | 0.002 | 0.024 | ↑ |
|  | Female | -0.132 | -1.407 | 1.000 | 0.983 | NS |
| **Mitotic Spindle** | Male | 0.352 | 3.413 | <0.0001 | 0.024 | ↑ |
|  | Female | -0.207 | -2.015 | 0.502 | 0.606 | NS |
| **Hedgehog Signaling** | Male | 0.418 | 3.192 | 0.027 | 0.028 | ↑ |
|  | Female | 0.142 | 1.222 | 0.967 | 0.992 | NS |
| **UV Response DN** | Male | 0.482 | 4.932 | <0.0001 | <0.0001 | ↑ |
|  | Female | 0.259 | 2.254 | 0.170 | 0.270 | NS |
| **UV Response UP** | Male | -0.322 | -2.766 | 0.004 | 0.090 | ↓ |
|  | Female | -0.218 | -2.108 | 0.499 | 0.571 | NS |
| **Epithelial Mesenchymal Transition** | Male | 0.221 | 2.258 | 0.260 | 0.461 | NS |
|  | Female | 0.392 | 3.425 | <0.0001 | 0.007 | ↑ |
| **TGF-β Signaling** | Male | 0.214 | 2.503 | 0.141 | 0.258 | NS |
|  | Female | 0.389 | 3.112 | <0.0001 | 0.028 | ↑ |
| **p53 Pathway** | Male | -0.226 | -2.306 | 0.130 | 0.297 | NS |
|  | Female | 0.302 | 2.732 | 0.006 | 0.062 | ↑ |
| **E2F Targets** | Male | 0.238 | 2.332 | 0.108 | 0.408 | NS |
|  | Female | -0.208 | -2.118 | 0.470 | 0.606 | NS |
| **MYC Targets V2** | Male | -0.243 | -2.398 | 0.184 | 0.282 | NS |
|  | Female | -0.272 | -2.716 | 0.1234 | 0.260 | NS |
| **KRAS Signaling UP** | Male | 0.207 | 1.923 | 0.632 | 0.857 | NS |
|  | Female | 0.204 | 1.827 | 0.667 | 0.640 | NS |
| **KRAS Signaling DN** | Male | 0.154 | 1.359 | 0.998 | 0.987 | NS |
|  | Female | 0.198 | 1.695 | 0.686 | 0.750 | NS |
| **Myogenesis** | Male | -0.204 | -1.896 | 0.519 | 0.790 | NS |
|  | Female | 0.228 | 2.147 | 0.229 | 0.339 | NS |

### **Supplementary Figure S5. GSEA Graphs for Male Cell Cycle, DNA Damage, and Tissue Remodeling Programs (Intranasal GHK-Cu).**

**A) DNA Repair	B) G2M Checkpoint	C) Mitotic Spindle**

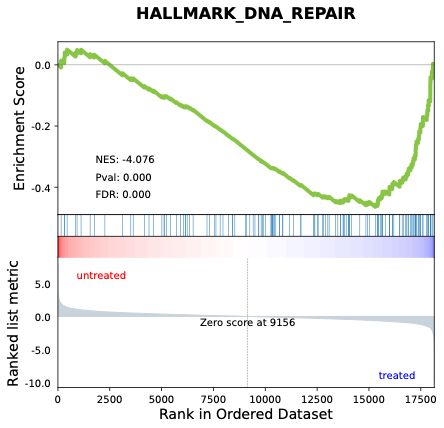

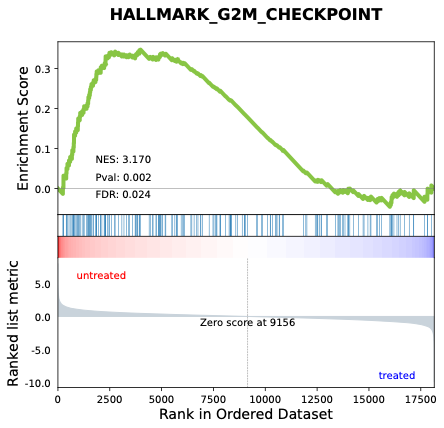

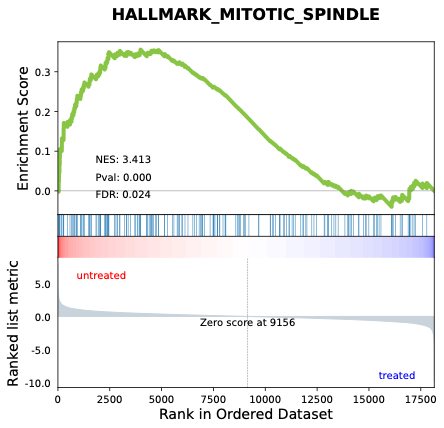

**D) Hedgehog Signaling	E) UV Response DN	F) UV Response UP**

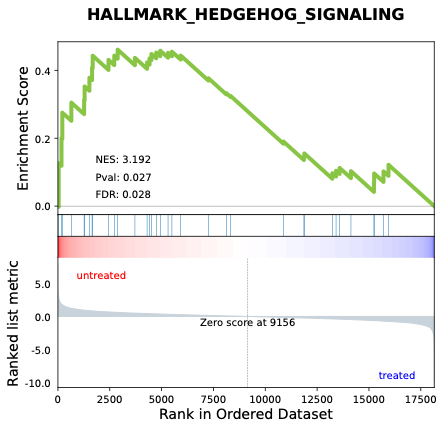

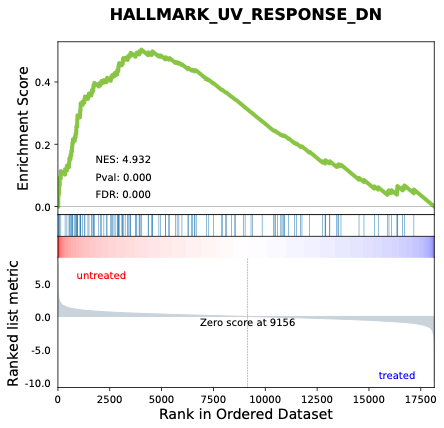

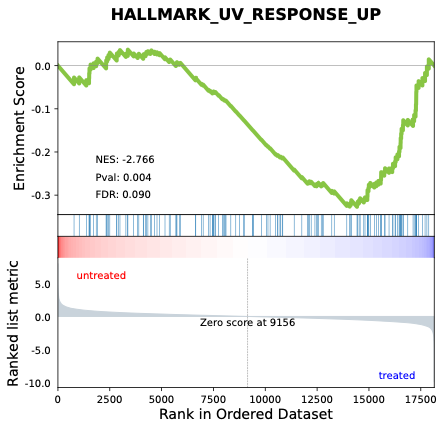

**G) Epithelial Mesenchymal Transition	H) TGF-β Signaling	I) p53 Pathway**

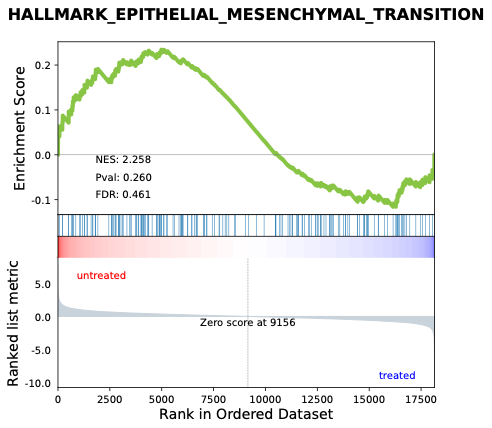

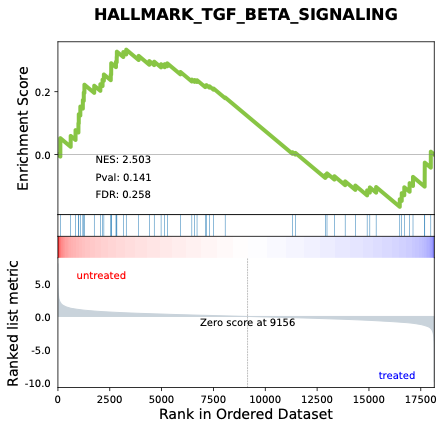

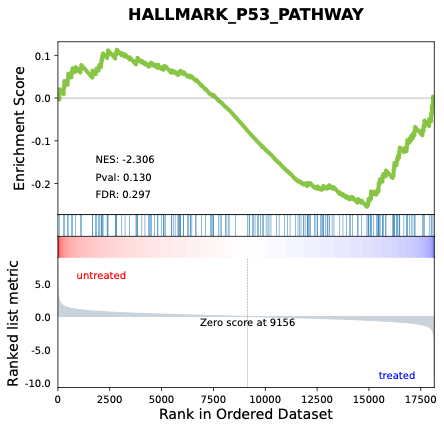

**J) E2F Targets	K) MYC Targets V2	L) KRAS Signaling UP**

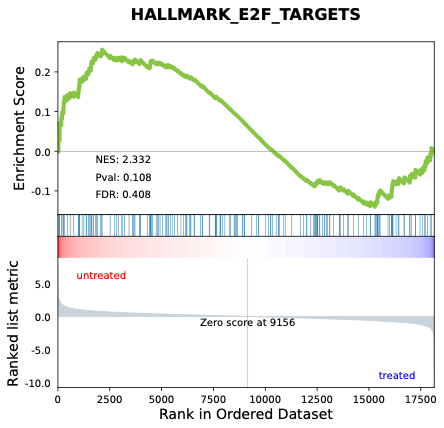

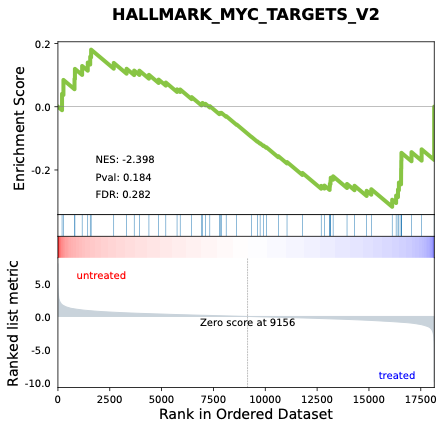

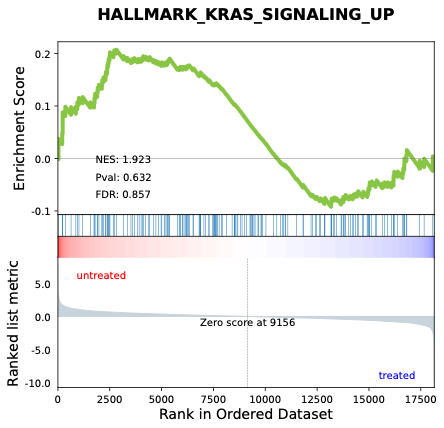

**M) KRAS Signaling DN	N) Myogenesis**

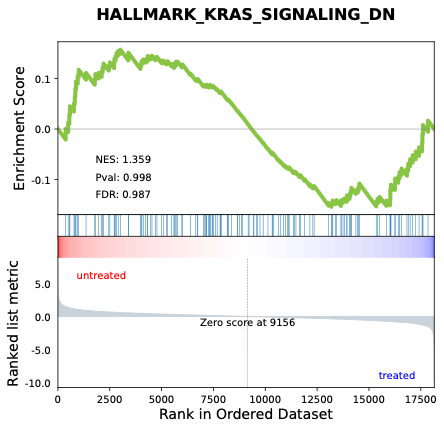

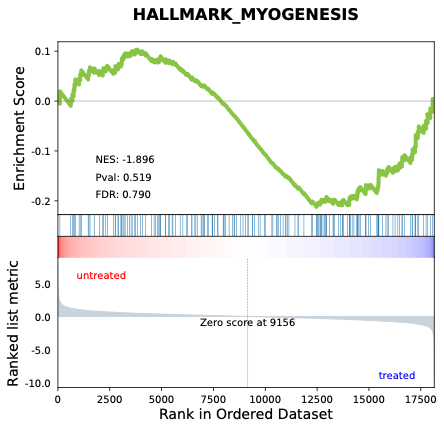

### **Supplementary Figure S6. GSEA Graphs for Female Cell Cycle, DNA Damage, and Tissue Remodeling Programs**

### **Supplementary Figure S6. GSEA Graphs for Female Cell Cycle, DNA Damage, and Tissue Remodeling Programs (Intranasal GHK-Cu).**

**A) DNA Repair	B) G2M Checkpoint	C) Mitotic Spindle**

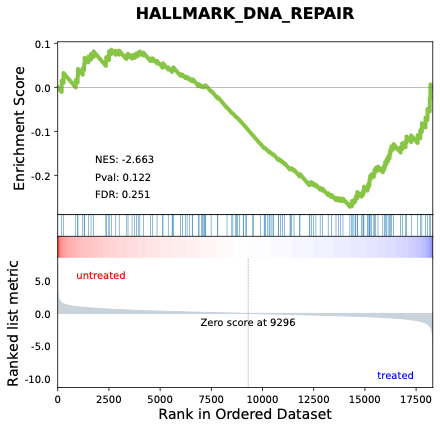

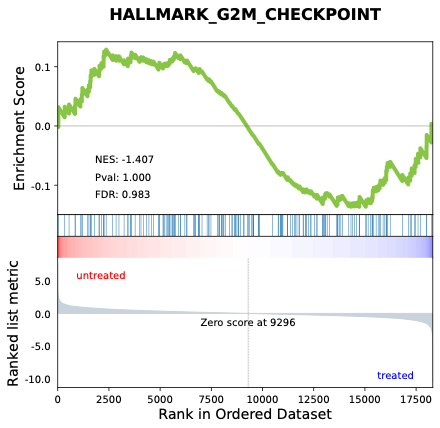

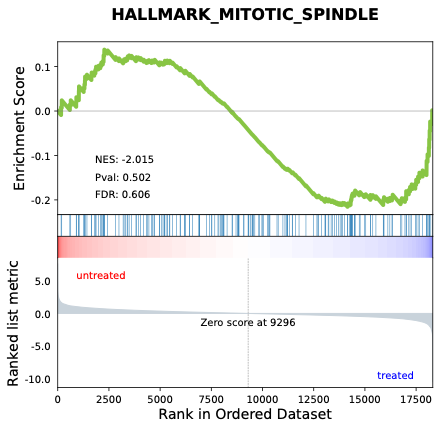

**D) Hedgehog Signaling	E) UV Response DN	F) UV Response UP**

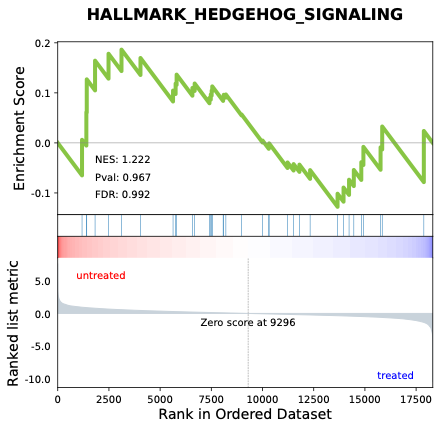

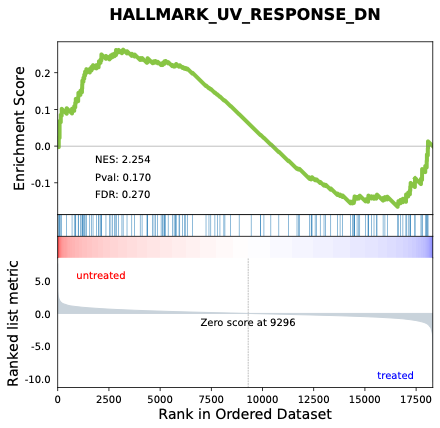

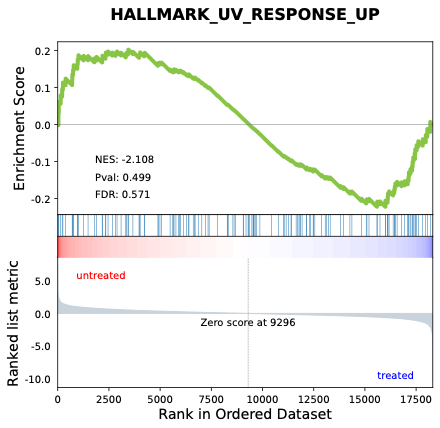

**G) Epithelial Mesenchymal Transition	H) TGF-β Signaling	I) p53 Pathway**

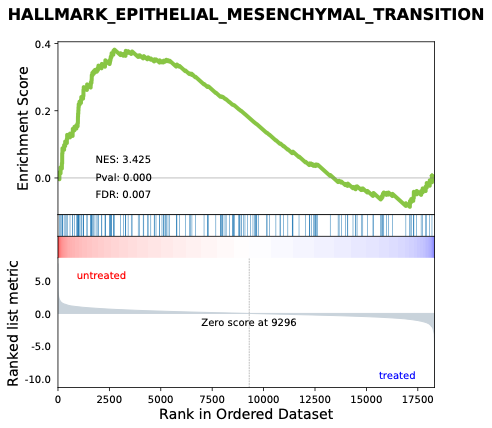

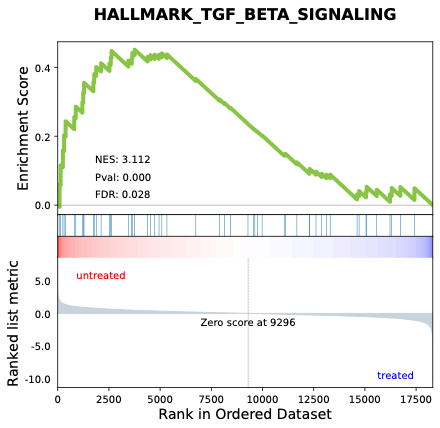

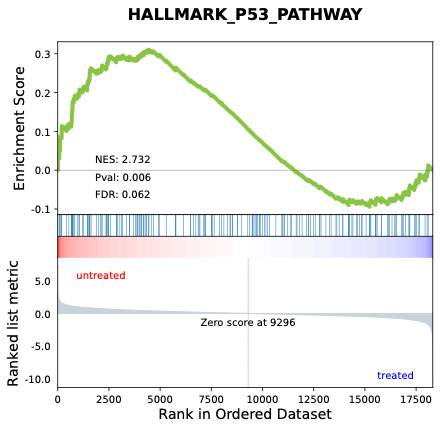

**J) E2F Targets	K) MYC Targets V2	L) KRAS Signaling UP**

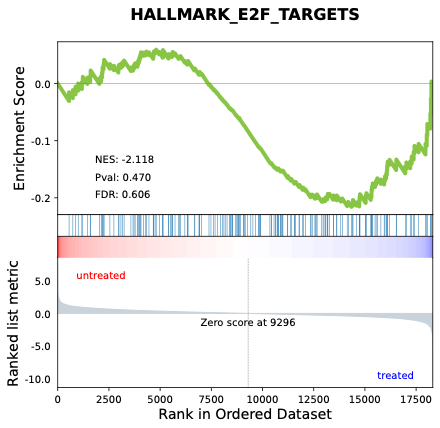

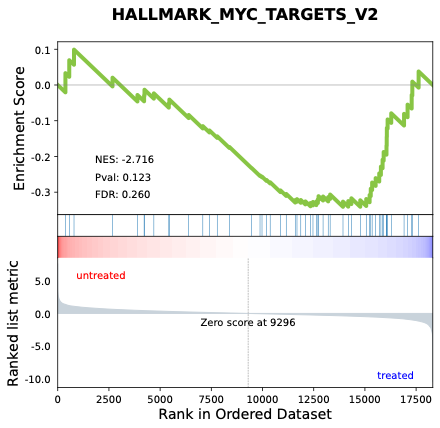

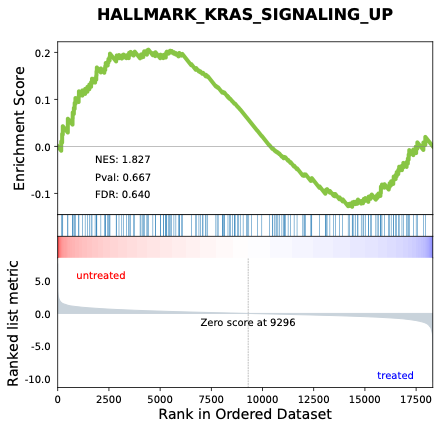

**M) KRAS Signaling DN	N) Myogenesis**

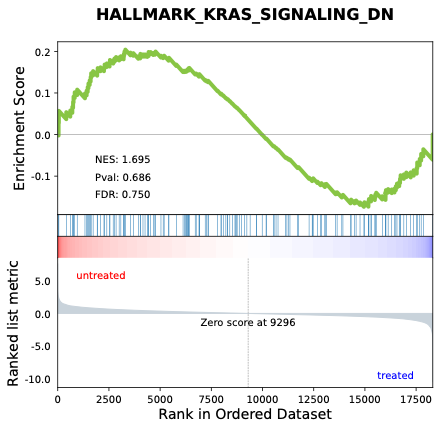

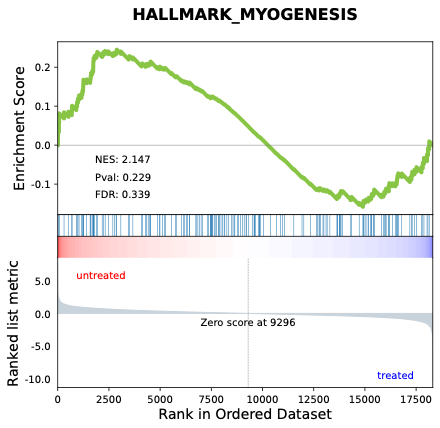

##### **Supplementary Table S4.** Additional hallmark pathways in male and female hippocampi following intranasal GHK-Cu treatment.

| **Pathway** | **Sex** | **Enrichment Score** | **NES** | ***P*-value** | **FDR** | **Regulation** |
| --- | --- | --- | --- | --- | --- | --- |
| **Estrogen Response Early** | Male | -0.156 | -1.628 | 0.895 | 0.937 | NS |
|  | Female | 0.284 | 2.377 | 0.066 | 0.188 | NS |
| **Estrogen Response Late** | Male | -0.258 | -2.398 | 0.106 | 0.263 | NS |
|  | Female | 0.336 | 3.022 | 0.002 | 0.030 | ↑ |
| **Spermatogenesis** | Male | -0.206 | -1.832 | 0.627 | 0.831 | NS |
|  | Female | -0.239 | -2.264 | 0.334 | 0.565 | NS |
| **Apical Junction** | Male | -0.189 | -1.660 | 0.800 | 0.936 | NS |
|  | Female | 0.256 | 2.372 | 0.074 | 0.181 | NS |
| **Apical Surface** | Male | 0.164 | 1.898 | 0.569 | 0.832 | NS |
|  | Female | 0.181 | 1.821 | 0.575 | 0.627 | NS |

### **Supplementary Figure S7. GSEA Graphs for** additional hallmark pathways in male hippocampi following intranasal GHK-Cu treatment.

**A) Estrogen Response Early	B) Estrogen Response Late	C) Spermatogenesis**

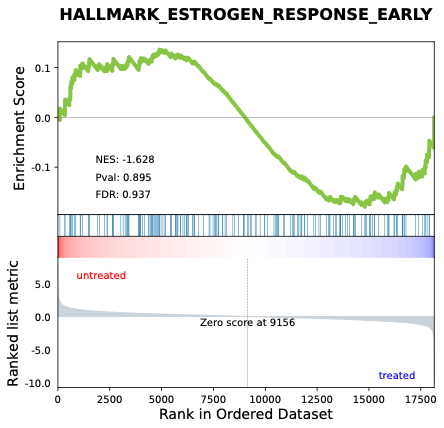

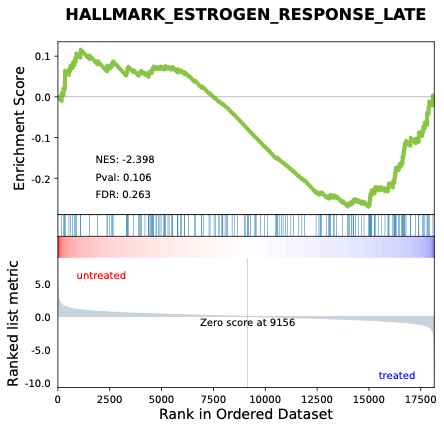

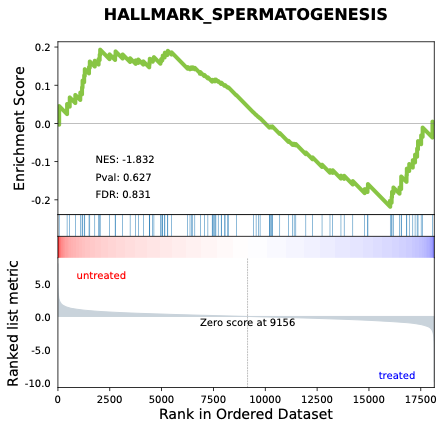

**D) Apical Junction	E) Apical Surface**

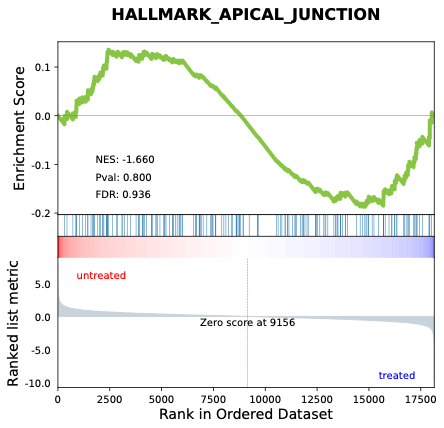

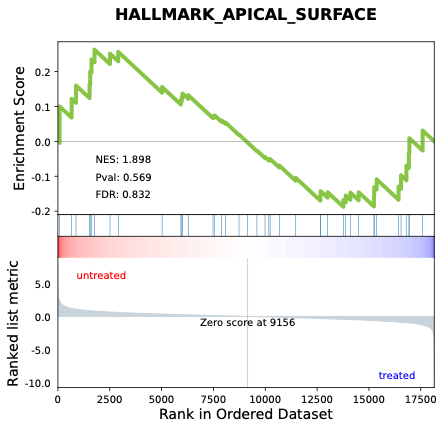

### **Supplementary Figure S8. GSEA Graphs Female** Additional Pathways (Intranasal GHK-Cu).

**A) Estrogen Response Early	B) Estrogen Response Late	C) Spermatogenesis**

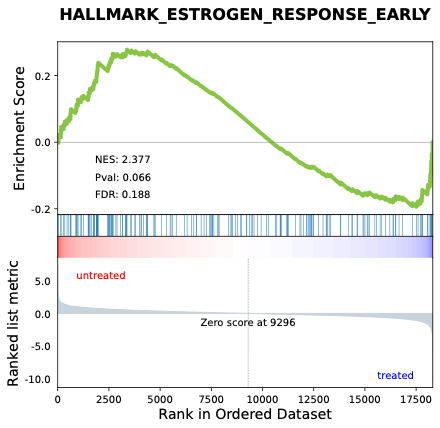

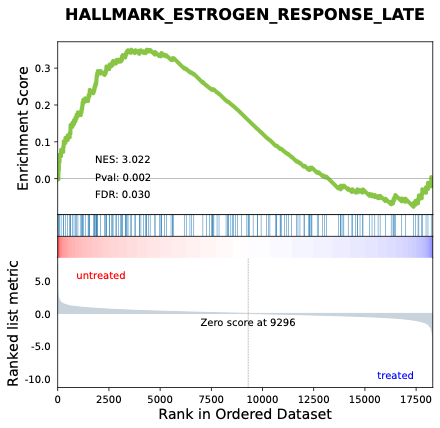

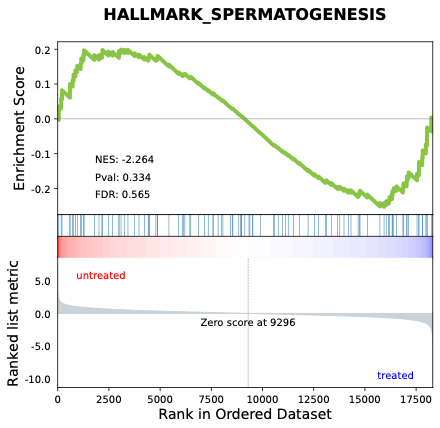
 **D) Apical Junction	E) Apical Surface**

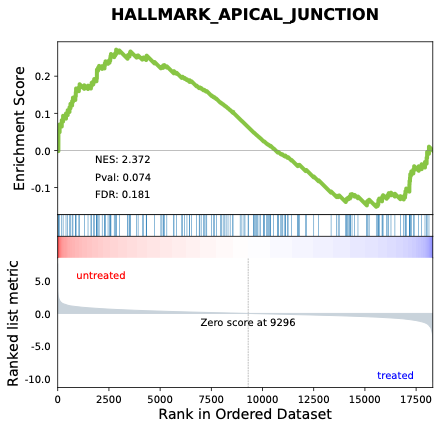

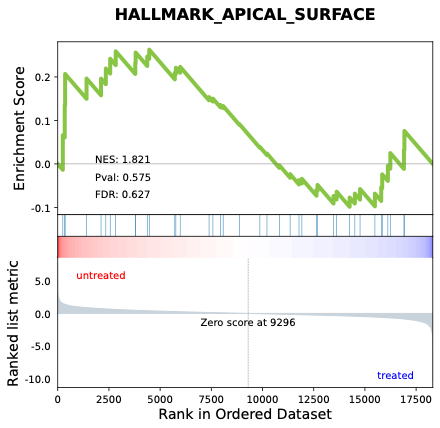
