## Supplementary material for "Middle-aged mice treated with GHK-Cu peptide administered intraperitoneally or intranasally show behavioral rescue but divergent hippocampal aging programs": Supp. Table 5-6 & Fig. 9-12

### **Supplementary Table S5. Immune & Inflammatory Pathways GSEA Profiles (Intraperitoneal GHK-Cu).**

| **Pathway** | **Sex** | **ES** | **NES** | **P-value** | **FDR** | **Regulation** |
| --- | --- | --- | --- | --- | --- | --- |
| **Interferon α Response** | Male | 0.241 | 1.981 | 0.478 | 0.845 | NS |
|  | Female | 0.205 | 2.110 | 0.554 | 0.788 | NS |
| **Interferon γ Response** | Male | 0.194 | 1.752 | 0.858 | 0.864 | NS |
|  | Female | 0.271 | 2.845 | 0.116 | 0.195 | NS |
| **IL2-STAT5 Signaling** | Male | 0.274 | 2.705 | 0.041 | 0.302 | NS |
|  | Female | -0.221 | -1.759 | 0.493 | 0.787 | NS |
| **IL6-JAK-STAT3 Signaling** | Male | 0.216 | 1.874 | 0.685 | 0.920 | NS |
|  | Female | 0.227 | 2.015 | 0.526 | 0.782 | NS |
| **Complement** | Male | -0.201 | -1.842 | 0.573 | 0.802 | NS |
|  | Female | 0.233 | 2.471 | 0.264 | 0.413 | NS |
| **Inflammatory Response** | Male | -0.236 | -2.113 | 0.183 | 0.600 | NS |
|  | Female | 0.214 | 2.235 | 0.523 | 0.712 | NS |
| **Coagulation** | Male | 0.223 | 2.183 | 0.528 | 0.635 | NS |
|  | Female | 0.245 | 2.489 | 0.262 | 0.419 | NS |
| **TNF-α Signaling via NFκB** | Male | 0.184 | 1.824 | 0.843 | 0.939 | NS |
|  | Female | 0.414 | 4.133 | <0.001 | 0.004 | ↑ |
| **Allograft Rejection** | Male | 0.244 | 2.194 | 0.330 | 0.711 | NS |
|  | Female | -0.156 | -1.385 | 0.951 | 1.000 | NS |
| **Angiogenesis** | Male | -0.179 | -1.618 | 0.730 | 0.876 | NS |
|  | Female | -0.148 | -1.335 | 0.905 | 1.000 | NS |
| **Hypoxia** | Male | -0.224 | -2.013 | 0.217 | 0.627 | NS |
|  | Female | 0.289 | 3.572 | 0.010 | 0.022 | ↑ |
| **NOTCH Signaling** | Male | -0.262 | -1.987 | 0.375 | 0.628 | NS |
|  | Female | 0.286 | 2.048 | 0.427 | 0.772 | NS |
| **WNT β-catenin signaling** | Male | -0.204 | -2.408 | 0.131 | 0.420 | NS |
|  | Female | 0.311 | 2.835 | 0.103 | 0.186 | NS |
| **Apoptosis** | Male | 0.224 | 1.921 | 0.478 | 0.897 | NS |
|  | Female | -0.204 | -1.544 | 0.774 | 0.986 | NS |

### **Supplementary Figure S9. Male Immune & Inflammatory Pathways GSEA Graphs (Intraperitoneal GHK-Cu).**

**A)** **Interferon α Response	B) Interferon γ Response	C) IL2-STAT5 Signaling**

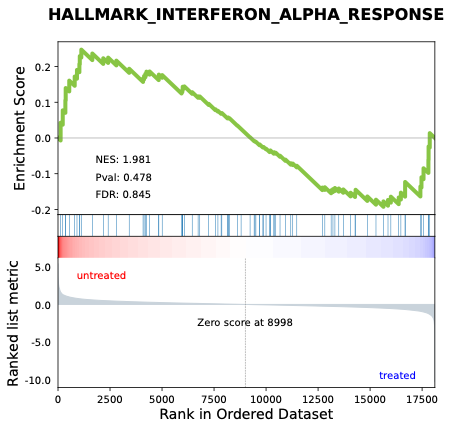

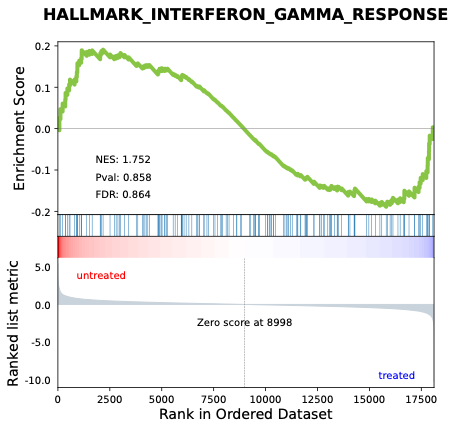

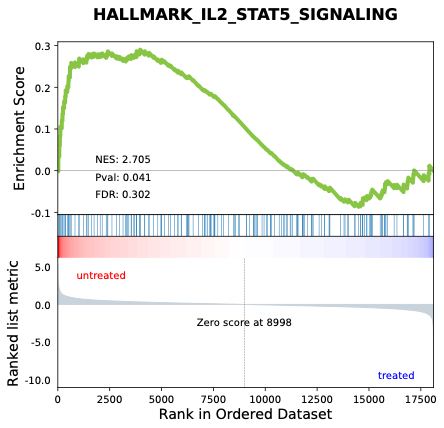

**D) IL6-JAK-STAT3 Signaling	E) Complement	F) Inflammatory Response**

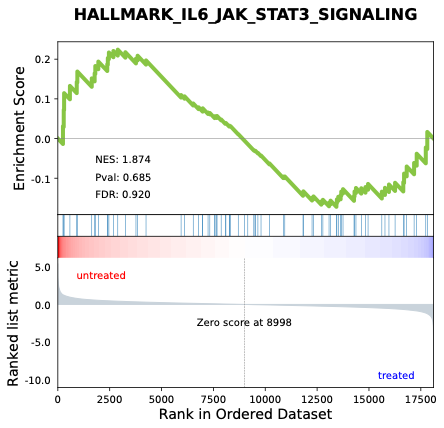

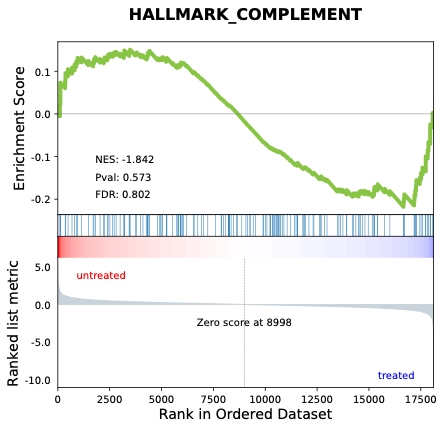

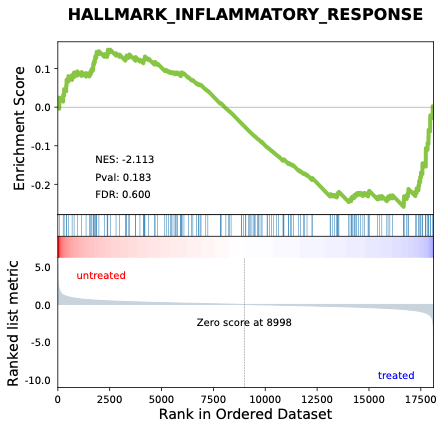

**G) Coagulation	H) TNF-α Signaling via NFκB	I) Allograft Rejection**

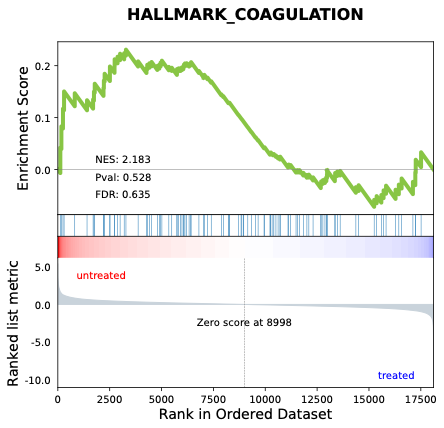

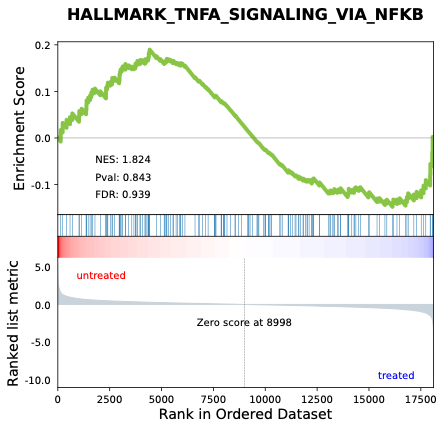

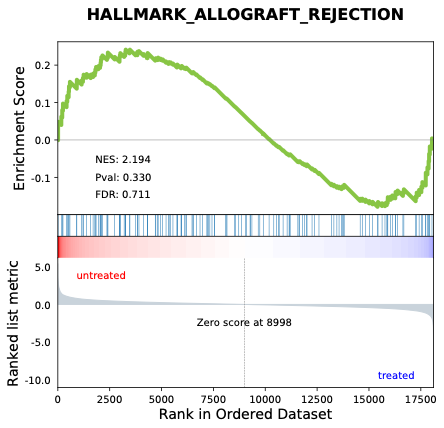

**J) Angiogenesis	K) Hypoxia	L) NOTCH Signaling**

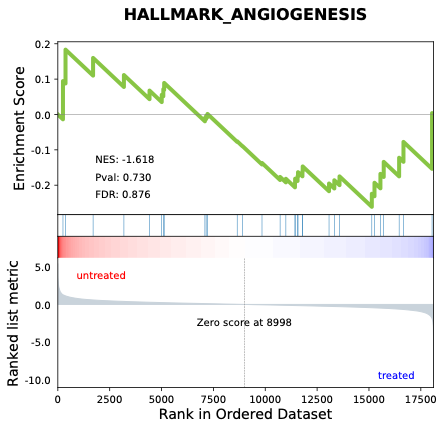

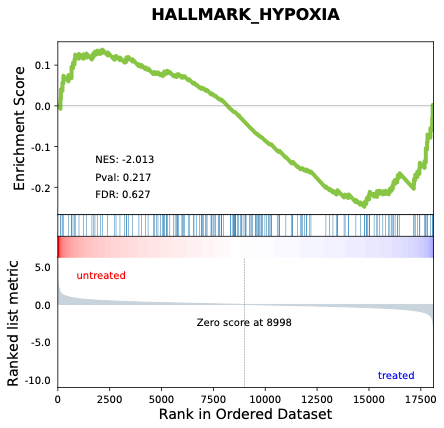

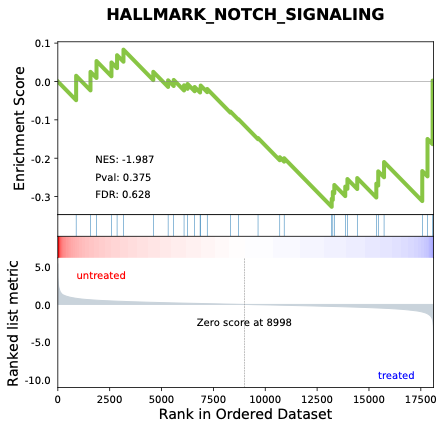

**M) WNT β-catenin signaling	N) Apoptosis**

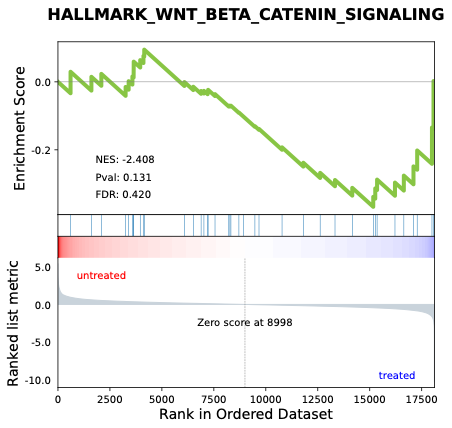

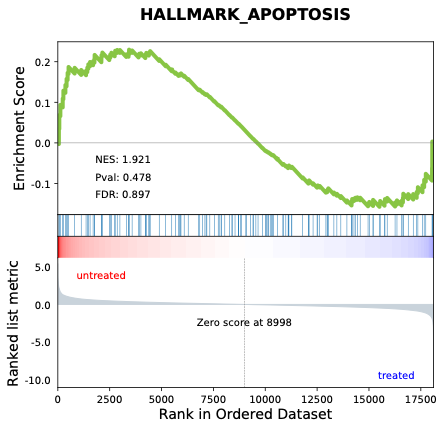

### **Supplementary Figure S10. Female Immune & Inflammatory Pathways GSEA Graphs (Intraperitoneal GHK-Cu).**

**A)** **Interferon α Response	B) Interferon γ Response	C) IL2-STAT5 Signaling**

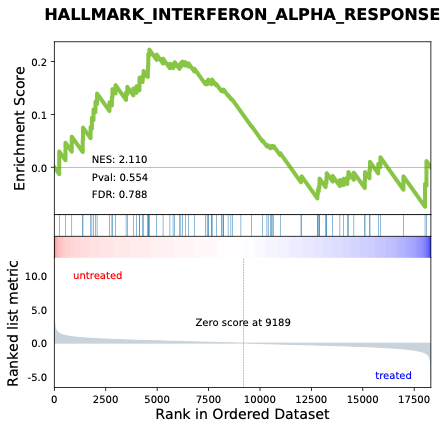

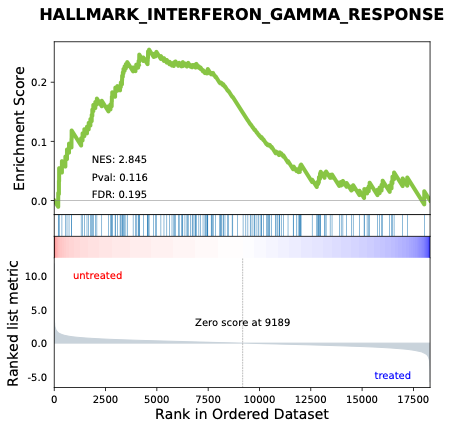

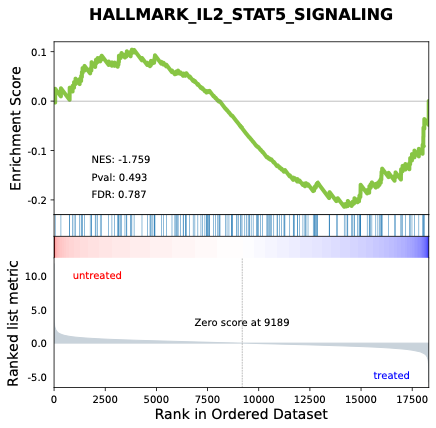

**D) IL6-JAK-STAT3 Signaling	E) Complement	F) Inflammatory Response**

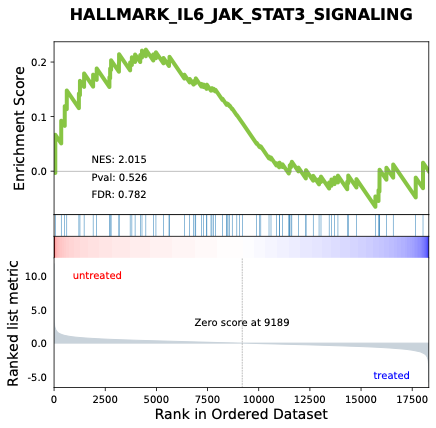

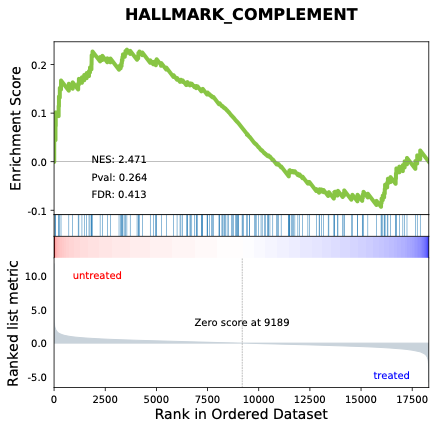

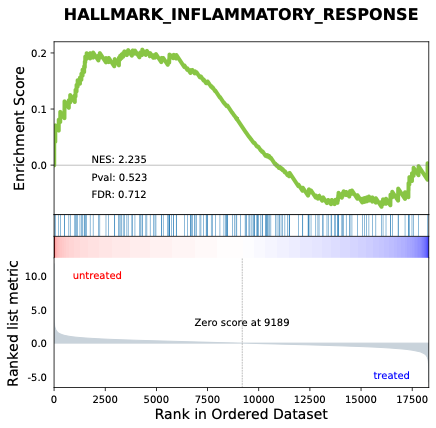

**G) Coagulation	H) TNF-α Signaling via NFκB	I) Allograft Rejection**

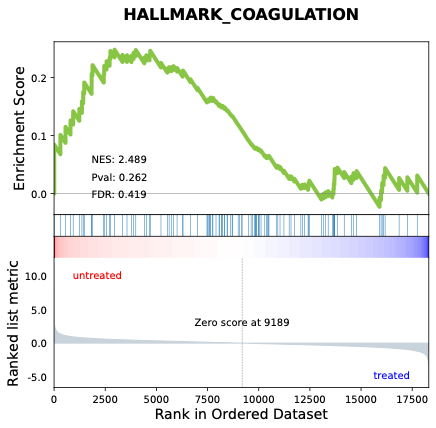

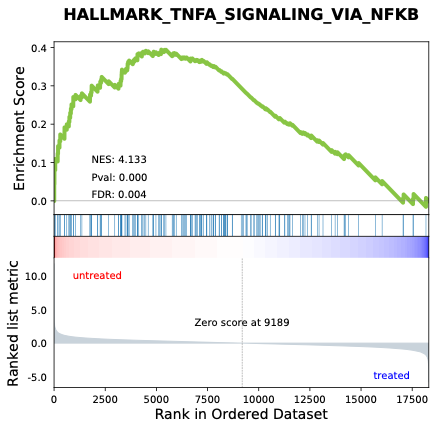

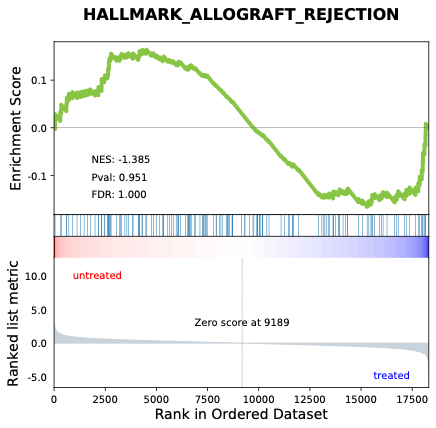

**J) Angiogenesis	K) Hypoxia	L) NOTCH Signaling**

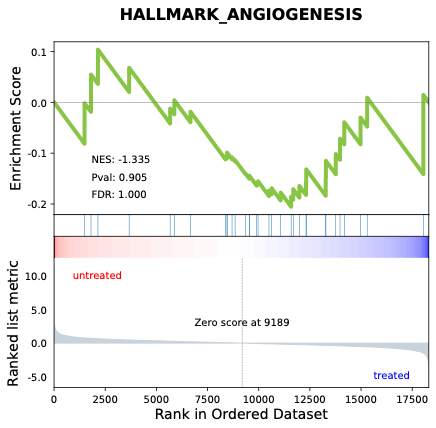

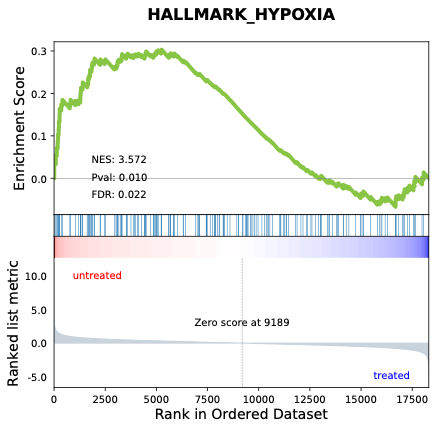

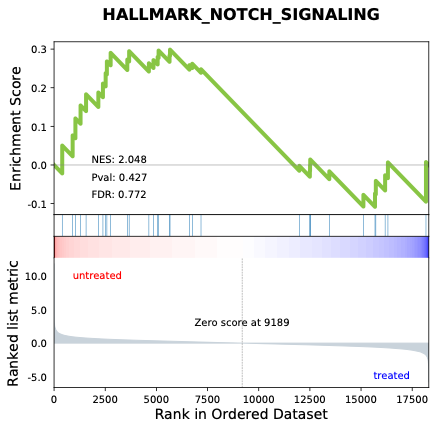

**M) WNT β-catenin signaling	N) Apoptosis**

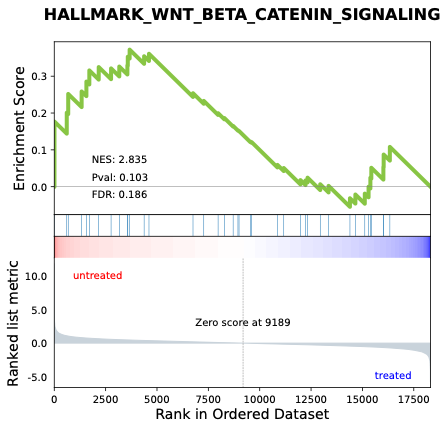

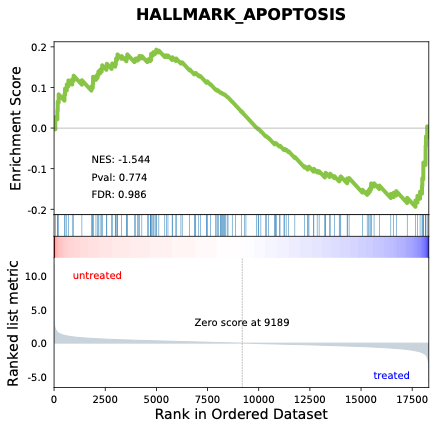

### **Supplementary Table S6. Metabolic & Detoxification Pathways GSEA Profiles (Intraperitoneal GHK-Cu).**

| **Pathway** | **Sex** | **Enrichment Score** | **NES** | ***P*-value** | **FDR** | **Regulation** |
| --- | --- | --- | --- | --- | --- | --- |
| **Androgen Response** | Male | -0.342 | -2.811 | 0.014 | 0.129 | ↓ |
|  | Female | -0.214 | -1.847 | 0.468 | 0.807 | NS |
| **Oxidative Phosphorylation** | Male | 0.298 | 2.935 | 0.019 | 0.247 | NS |
|  | Female | 0.471 | 4.965 | <0.001 | <0.001 | ↑ |
| **Reactive Oxygen Species** | Male | 0.241 | 2.470 | 0.257 | 0.449 | NS |
|  | Female | 0.392 | 3.401 | 0.021 | 0.037 | ↑ |
| **MYC Targets V1** | Male | 0.246 | 2.467 | 0.173 | 0.409 | NS |
|  | Female | 0.421 | 4.344 | <0.001 | 0.002 | ↑ |
| **PI3K-AKT-mTOR Signaling** | Male | -0.284 | -2.393 | 0.071 | 0.372 | NS |
|  | Female | 0.185 | 2.095 | 0.652 | 0.774 | NS |
| **MTORC1 Signaling** | Male | 0.217 | 2.115 | 0.416 | 0.704 | NS |
|  | Female | 0.246 | 2.999 | 0.023 | 0.136 | ↑ |
| **Adipogenesis** | Male | -0.192 | -1.693 | 0.770 | 0.907 | NS |
|  | Female | 0.231 | 2.784 | 0.106 | 0.207 | NS |
| **Fatty Acid Metabolism** | Male | 0.279 | 2.520 | 0.113 | 0.429 | NS |
|  | Female | 0.205 | 2.212 | 0.508 | 0.709 | NS |
| **Cholesterol Homeostasis** | Male | 0.286 | 2.309 | 0.226 | 0.558 | NS |
|  | Female | 0.217 | 1.971 | 0.520 | 0.804 | NS |
| **Xenobiotic Metabolism** | Male | 0.191 | 1.788 | 0.743 | 0.900 | NS |
|  | Female | 0.281 | 2.971 | 0.011 | 0.136 | ↑ |
| **Bile Acid Metabolism** | Male | -0.213 | -1.720 | 0.541 | 0.968 | NS |
|  | Female | -0.224 | -1.968 | 0.336 | 0.713 | NS |
| **Glycolysis** | Male | 0.152 | 1.596 | 0.951 | 0.934 | NS |
|  | Female | 0.218 | 2.592 | 0.263 | 0.331 | NS |
| **Heme Metabolism** | Male | -0.251 | -2.300 | 0.129 | 0.396 | NS |
|  | Female | -0.214 | -1.842 | 0.326 | 0.744 | NS |
| **Peroxisome** | Male | -0.214 | -1.920 | 0.426 | 0.703 | NS |
|  | Female | 0.189 | 1.890 | 0.791 | 0.856 | NS |
| **Unfolded Protein Response** | Male | 0.236 | 2.355 | 0.191 | 0.533 | NS |
|  | Female | 0.215 | 2.174 | 0.447 | 0.732 | NS |
| **Protein Secretion** | Male | -0.236 | -2.113 | 0.266 | 0.546 | NS |
|  | Female | -0.401 | -3.140 | <0.001 | 0.024 | ↓ |
| **Pancreatic β-cell signaling** | Male | -0.118 | -1.360 | 0.873 | 0.968 | NS |
|  | Female | 0.321 | 2.626 | 0.141 | 0.320 | NS |

### **Supplementary Figure S11. Male Metabolic & Detoxification Pathways GSEA Graphs (Intraperitoneal GHK-Cu).**

**A) Androgen Response	B) Oxidative Phosphorylation	C) Reactive Oxygen Species**

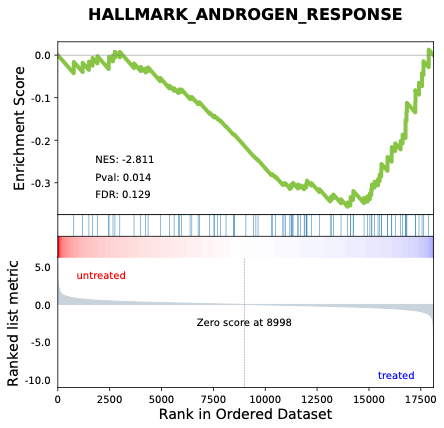

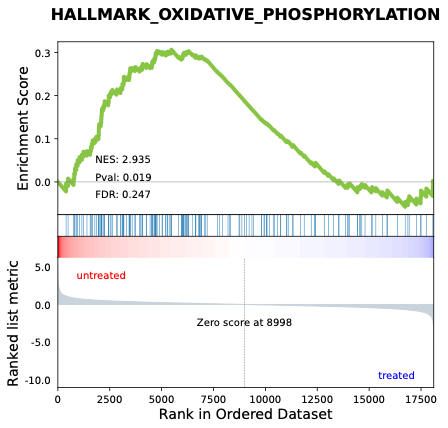

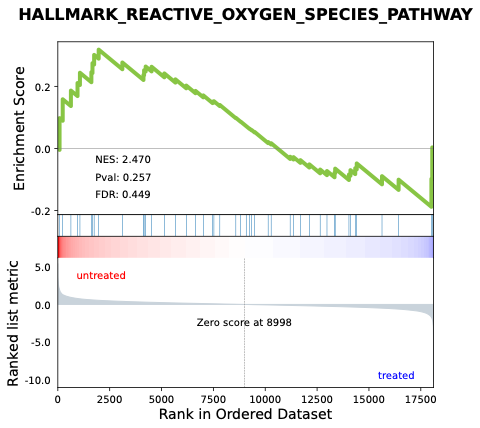

**D) MYC Targets V1	E) PI3K-AKT-mTOR Signaling	F) MTORC1 Signaling**

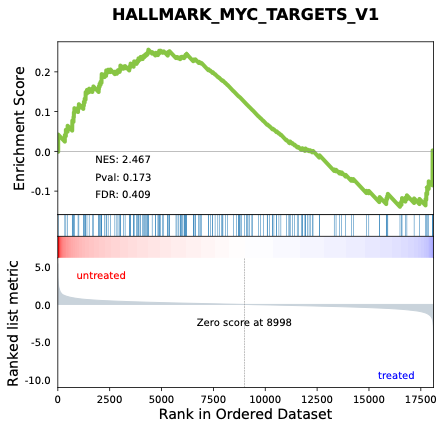

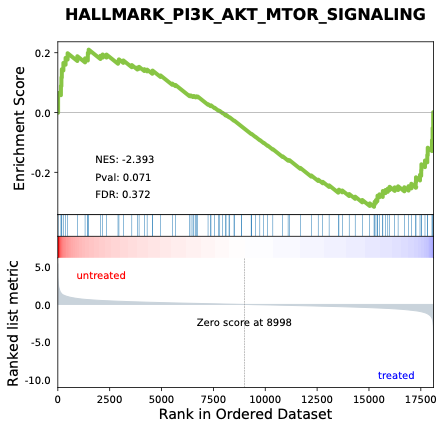

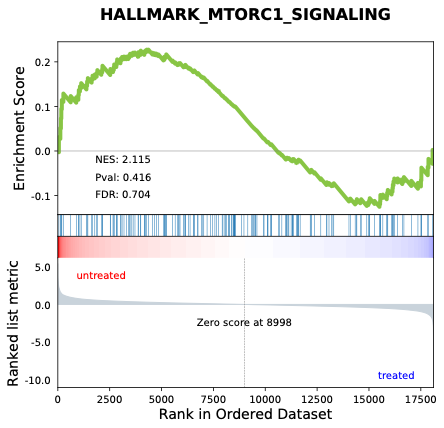

**G) Adipogenesis	H) Fatty Acid Metabolism	I) Cholesterol Homeostasis**

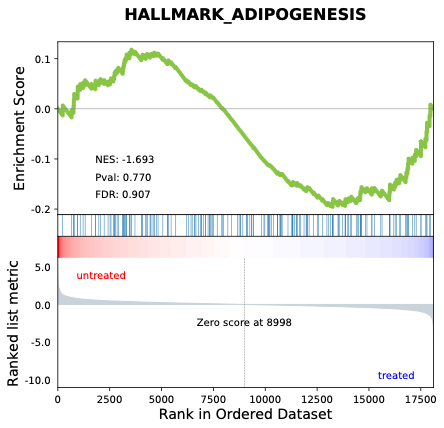

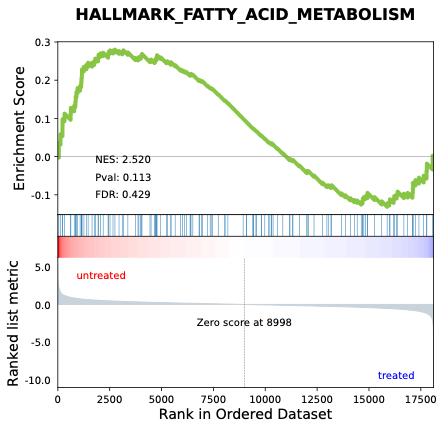

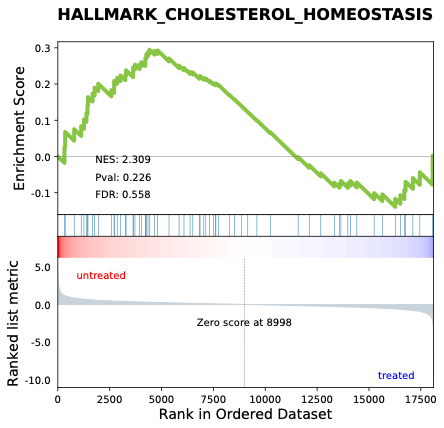

**J) Xenobiotic Metabolism	K) Bile Acid Metabolism	L) Glycolysis**

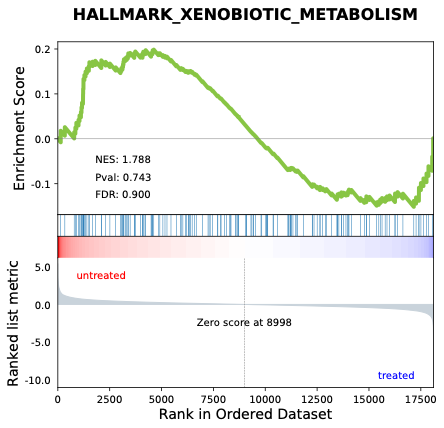

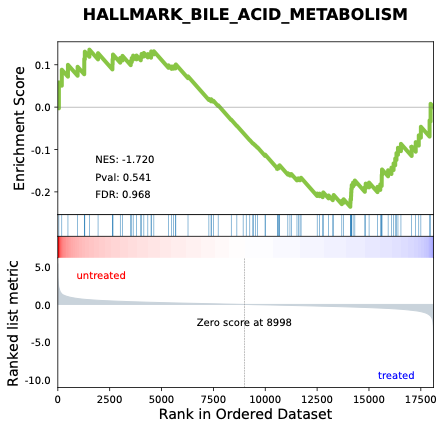

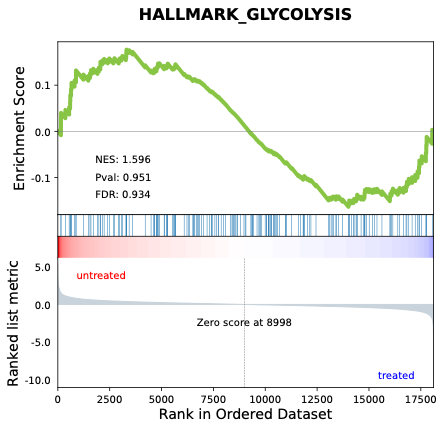

**M) Heme Metabolism	N) Peroxisome	O) Unfolded Protein Response**

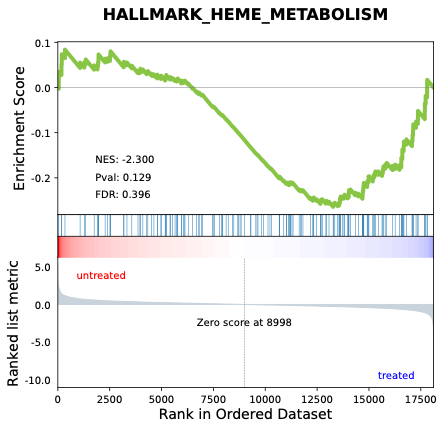

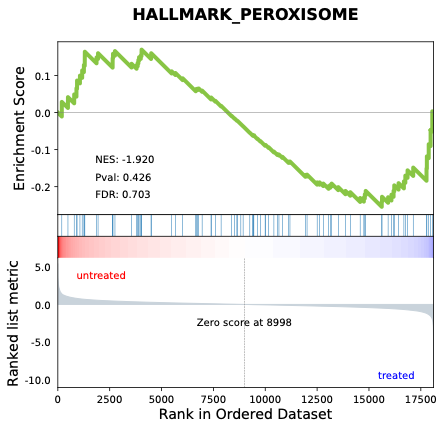

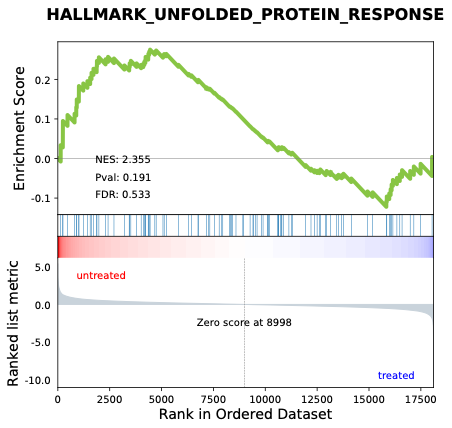

**P) Protein Secretion	Q) Pancreatic β-cell signaling**

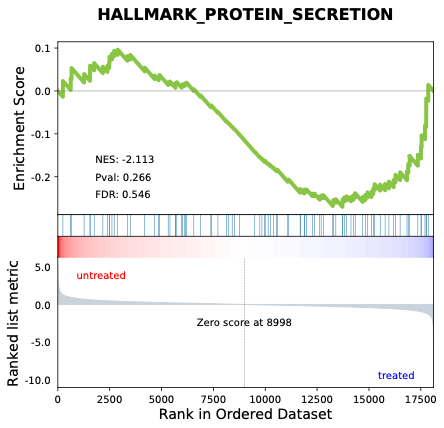

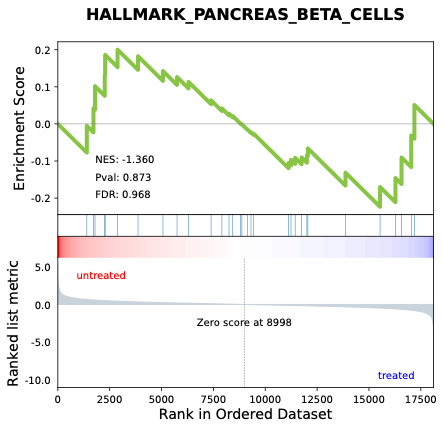

### **Supplementary Figure S12. Female Metabolic & Detoxification Pathways GSEA Graphs (Intraperitoneal GHK-Cu).**

**A) Androgen Response	B) Oxidative Phosphorylation	C) Reactive Oxygen Species**

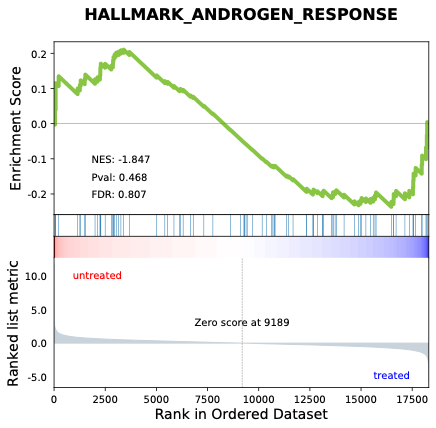

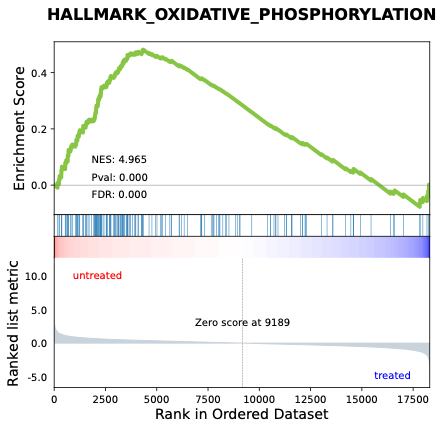

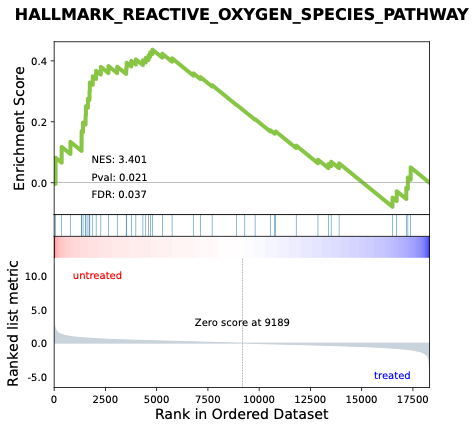

**D) MYC Targets V1	E) PI3K-AKT-mTOR Signaling	F) MTORC1 Signaling**

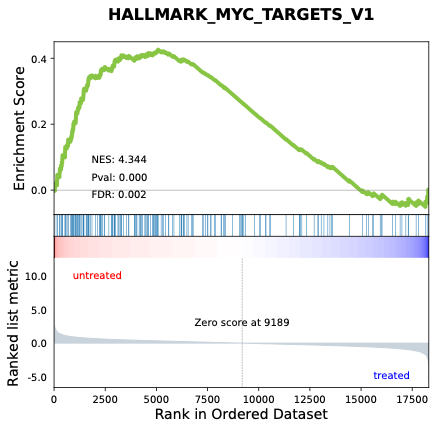

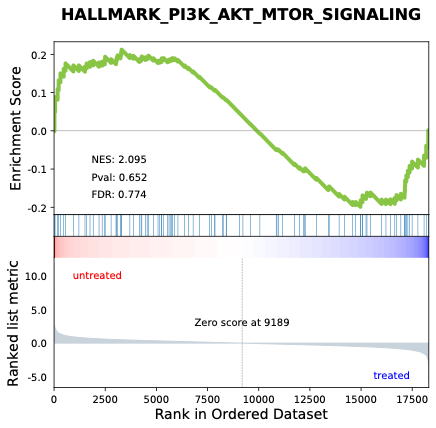

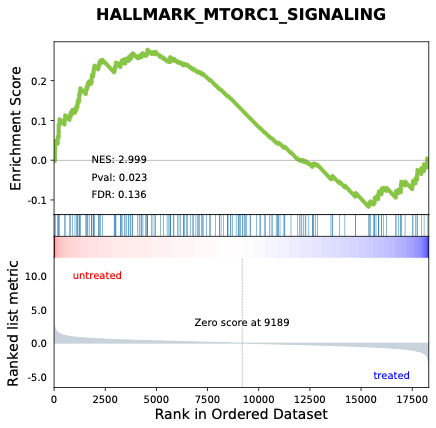

**G) Adipogenesis	H) Fatty Acid Metabolism	I) Cholesterol Homeostasis**

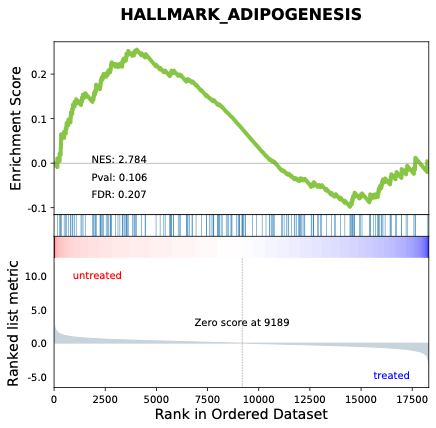

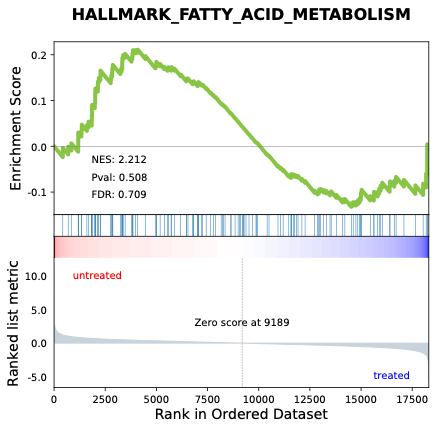

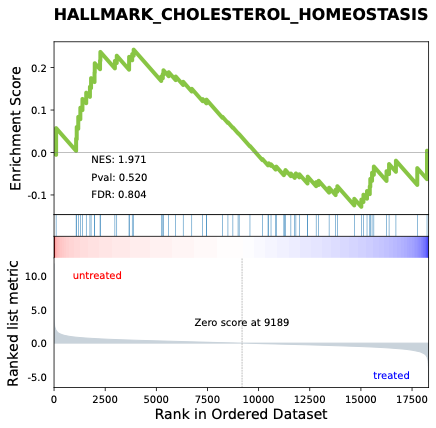

**J) Xenobiotic Metabolism	K) Bile Acid Metabolism	L) Glycolysis**

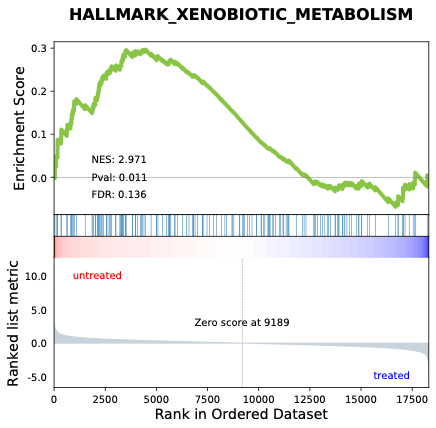

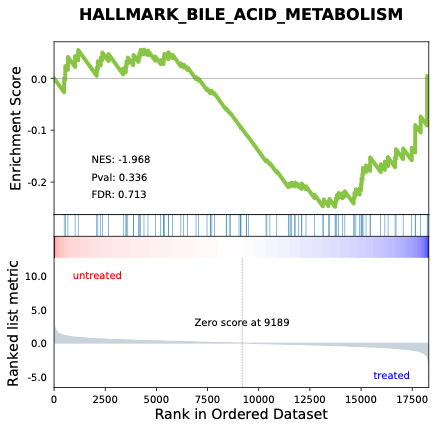

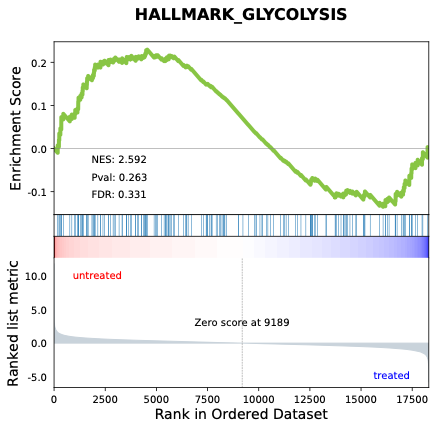

**M) Heme Metabolism	N) Peroxisome	O) Unfolded Protein Response**

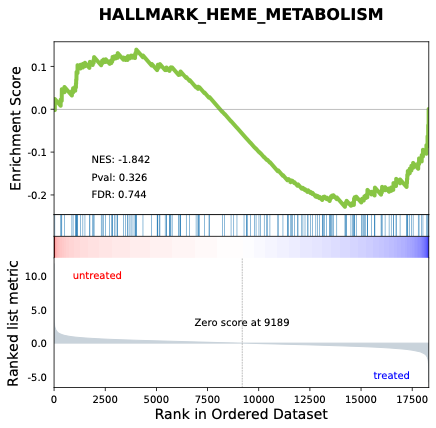

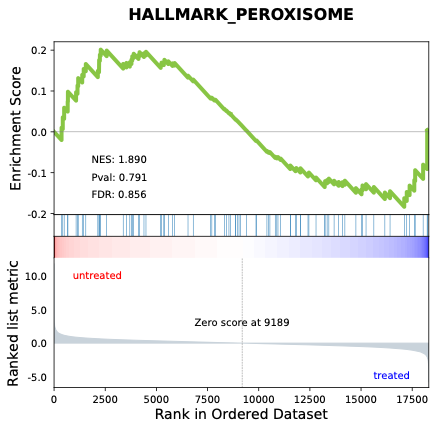

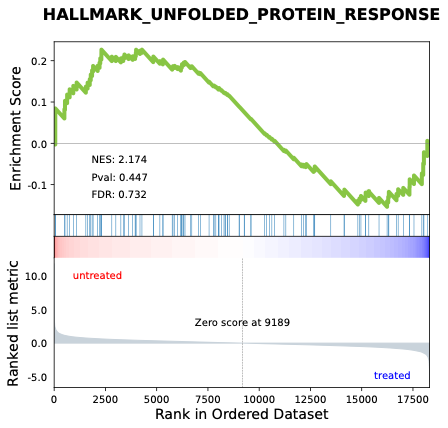

**P) Protein Secretion	Q) Pancreatic β-cell signaling**

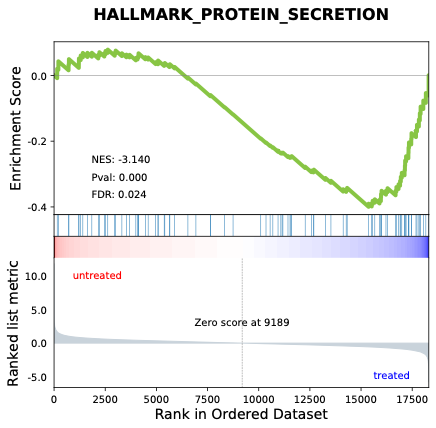

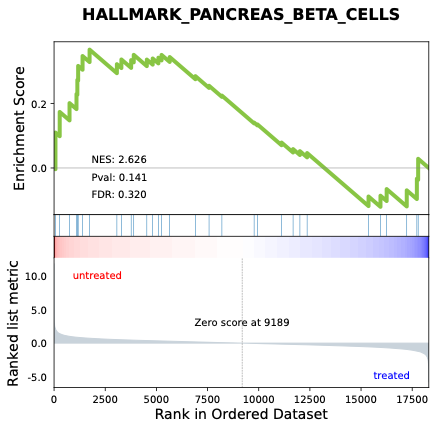
