## Supplementary material for "Middle-aged mice treated with GHK-Cu peptide administered intraperitoneally or intranasally show behavioral rescue but divergent hippocampal aging programs": Supp. Table 7-8 & Fig. 13-16

### **Supplementary Table S7. Cell Cycle, DNA Damage, and Tissue Remodeling Pathways GSEA Profiles (Intraperitoneal GHK-Cu).**

| **Pathway** | **Sex** | **Enrichment Score** | **NES** | ***P*-value** | **FDR** | **Regulation** |
| --- | --- | --- | --- | --- | --- | --- |
| **DNA Repair** | Male | 0.331 | 3.294 | 0.007 | 0.154 | NS |
|  | Female | 0.456 | 5.575 | <0.001 | <0.001 | ↑ |
| **G2M Checkpoint** | Male | 0.196 | 1.795 | 0.925 | 0.932 | NS |
|  | Female | -0.224 | -1.921 | 0.257 | 0.730 | NS |
| **Mitotic Spindle** | Male | -0.389 | -3.351 | <0.001 | 0.033 | ↓ |
|  | Female | -0.402 | -3.385 | <0.001 | 0.006 | ↓ |
| **Hedgehog Signaling** | Male | -0.141 | -1.638 | 0.827 | 0.894 | NS |
|  | Female | -0.198 | -1.657 | 0.639 | 0.858 | NS |
| **UV Response DN** | Male | -0.312 | -2.724 | 0.008 | 0.141 | ↓ |
|  | Female | -0.298 | -3.029 | <0.001 | 0.024 | ↓ |
| **UV Response UP** | Male | 0.208 | 1.994 | 0.470 | 0.870 | NS |
|  | Female | 0.332 | 3.220 | 0.011 | 0.069 | ↑ |
| **Epithelial Mesenchymal Transition** | Male | -0.183 | -1.583 | 0.846 | 0.874 | NS |
|  | Female | -0.218 | -1.807 | 0.345 | 0.754 | NS |
| **TGF-β Signaling** | Male | -0.146 | -1.704 | 0.617 | 0.941 | NS |
|  | Female | -0.285 | -2.200 | 0.191 | 0.369 | NS |
| **p53 Pathway** | Male | 0.291 | 2.967 | 0.020 | 0.325 | NS |
|  | Female | 0.318 | 3.754 | <0.001 | 0.012 | ↑ |
| **E2F Targets** | Male | 0.289 | 2.825 | 0.041 | 0.285 | NS |
|  | Female | 0.338 | 3.790 | 0.002 | 0.013 | ↑ |
| **MYC Targets V2** | Male | 0.327 | 2.797 | 0.050 | 0.253 | NS |
|  | Female | 0.233 | 2.057 | 0.477 | 0.793 | NS |
| **KRAS Signaling UP** | Male | 0.232 | 2.190 | 0.391 | 0.667 | NS |
|  | Female | -0.158 | -1.196 | 0.997 | 0.990 | NS |
| **KRAS Signaling DN** | Male | -0.241 | -2.164 | 0.205 | 0.567 | NS |
|  | Female | -0.207 | -1.698 | 0.562 | 0.842 | NS |
| **Myogenesis** | Male | -0.198 | -1.655 | 0.749 | 0.915 | NS |
|  | Female | 0.178 | 1.838 | 0.964 | 0.840 | NS |

### **Supplementary Figure S13. Male Cell Cycle, DNA Damage, and Tissue Remodeling Pathways GSEA Graphs (Intraperitoneal GHK-Cu).**

**A) DNA Repair	B) G2M Checkpoint	C) Mitotic Spindle**

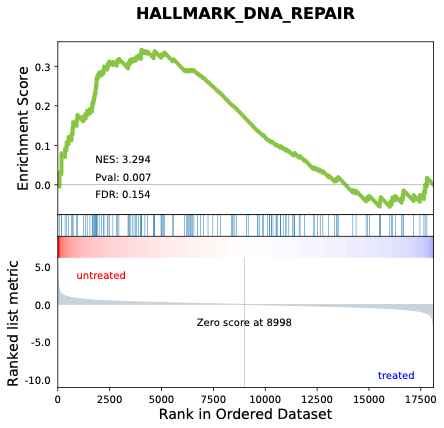

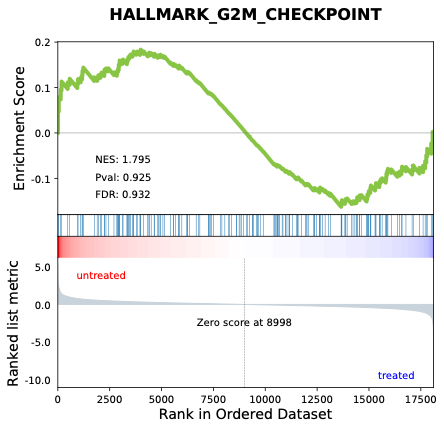

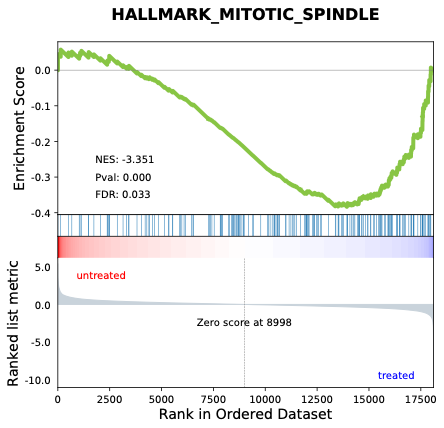

**D) Hedgehog Signaling	E) UV Response DN	F) UV Response UP**

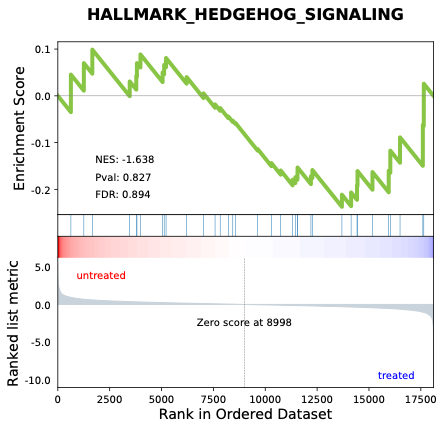

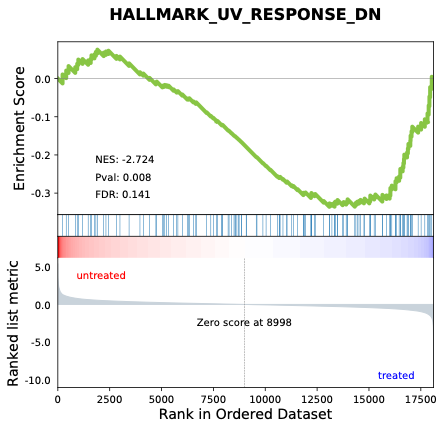

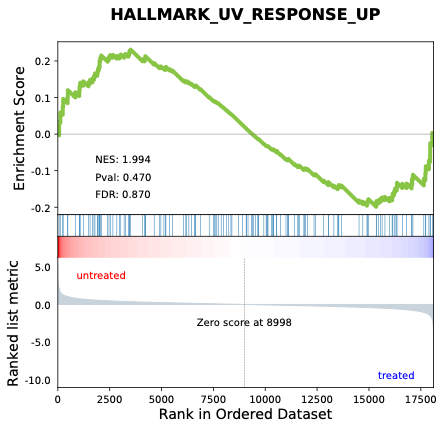

**G) Epithelial Mesenchymal Transition	H) TGF-β Signaling	I) p53 Pathway**

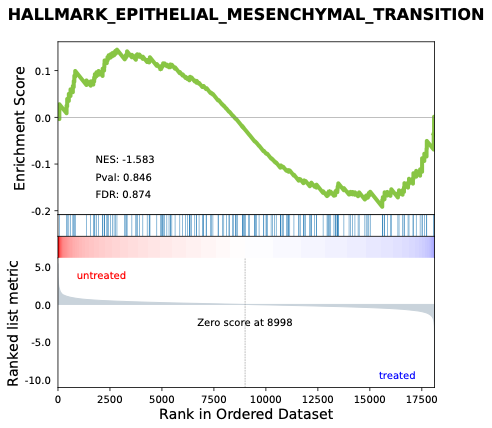

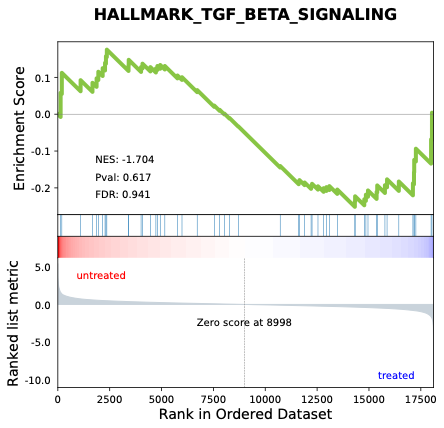

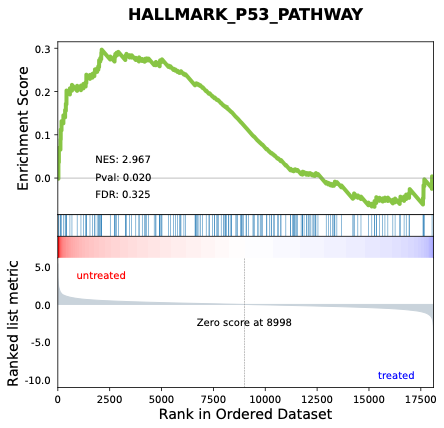

**J) E2F Targets	K) MYC Targets V2	L) KRAS Signaling UP**

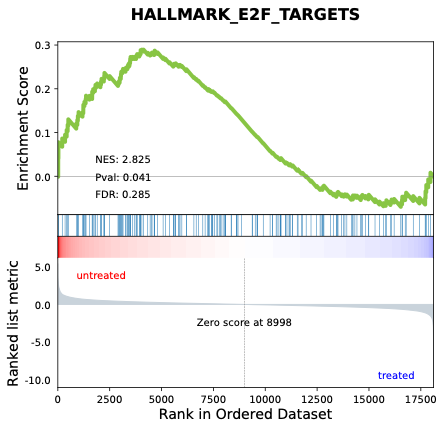

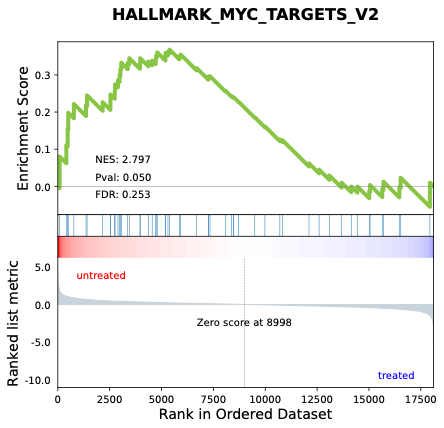

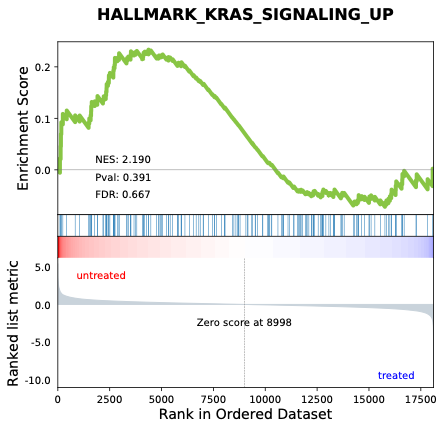

**M) KRAS Signaling DN	N) Myogenesis**

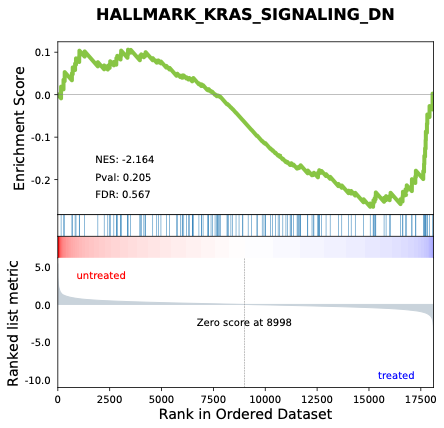

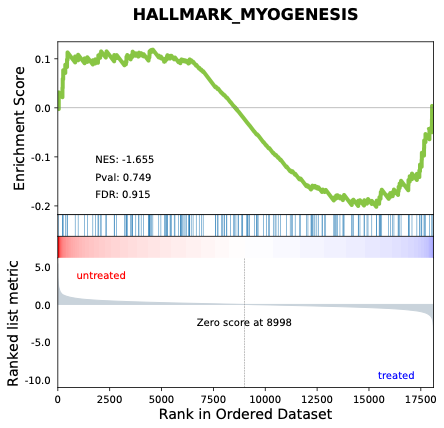

### **Supplementary Figure S14. Female Cell Cycle, DNA Damage, and Tissue Remodeling Pathways GSEA Graphs (Intraperitoneal GHK-Cu).**

**A) DNA Repair	B) G2M Checkpoint	C) Mitotic Spindle**

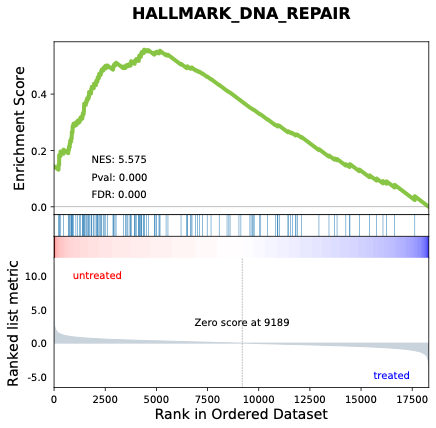

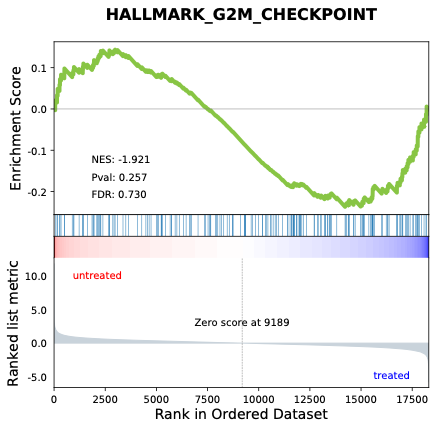

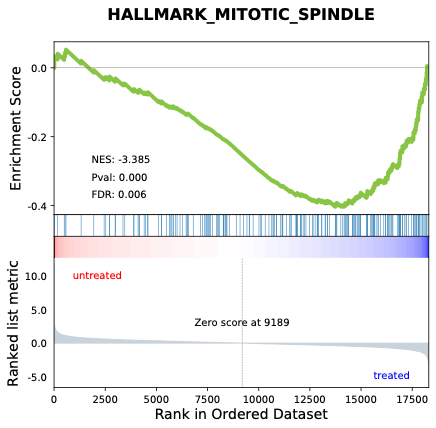

**D) Hedgehog Signaling	E) UV Response DN	F) UV Response UP**

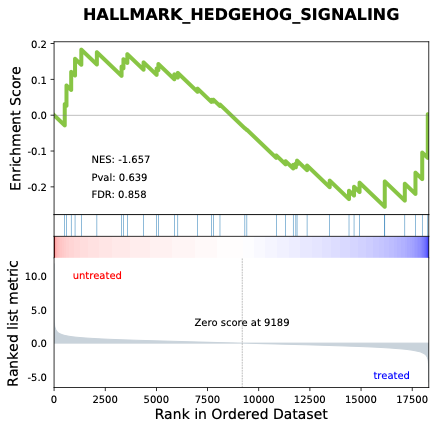

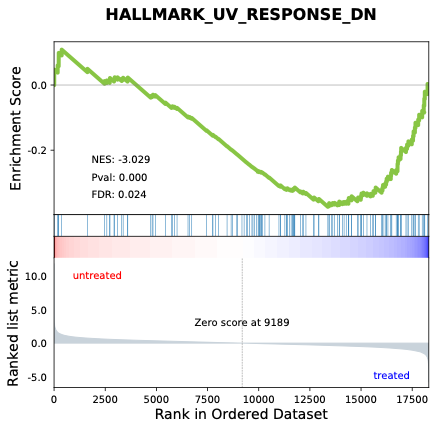

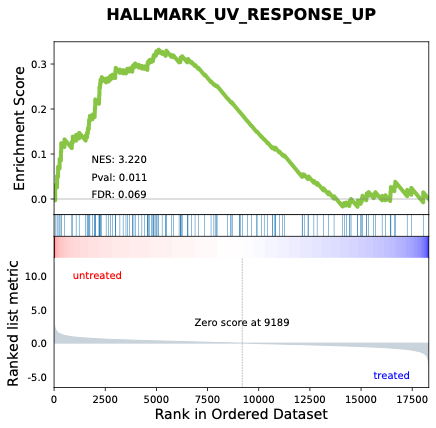

**G) Epithelial Mesenchymal Transition	H) TGF-β Signaling	I) p53 Pathway**

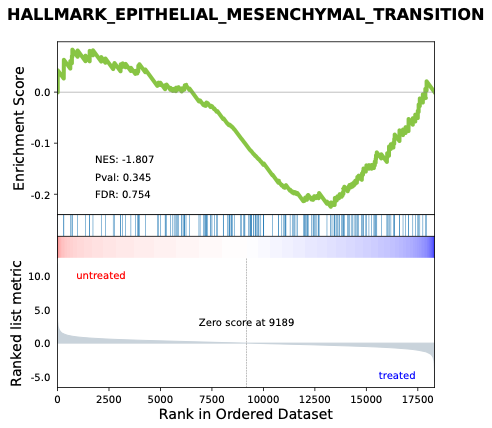

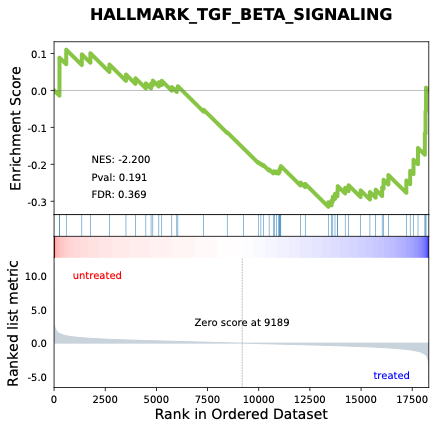

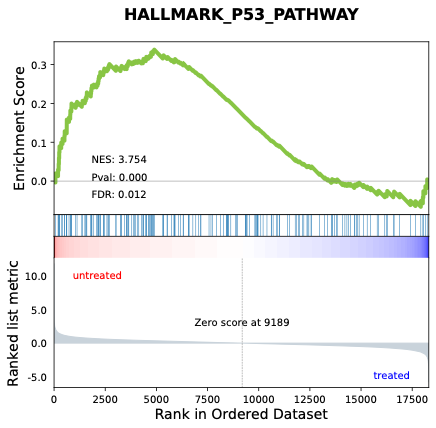

**J) E2F Targets	K) MYC Targets V2	L) KRAS Signaling UP**

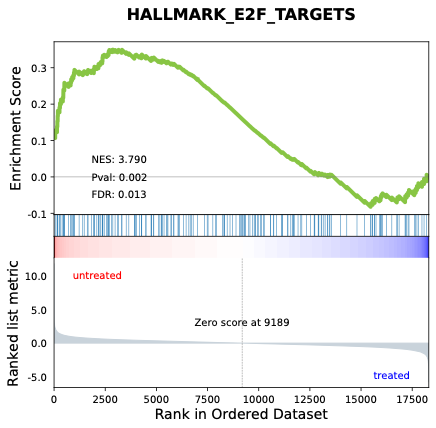

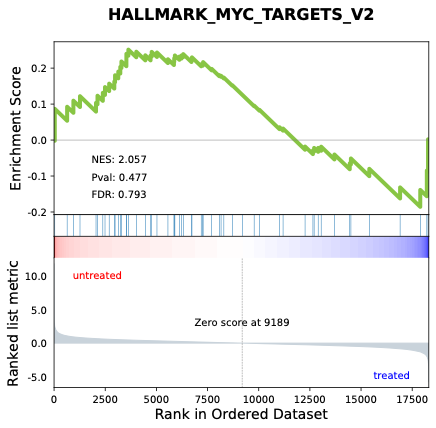

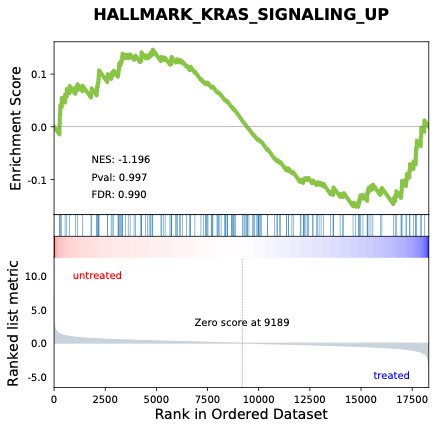

**M) KRAS Signaling DN	N) Myogenesis**

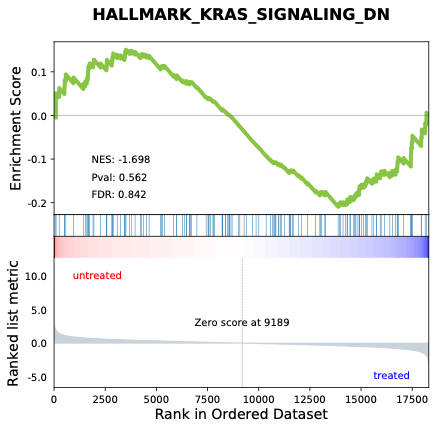

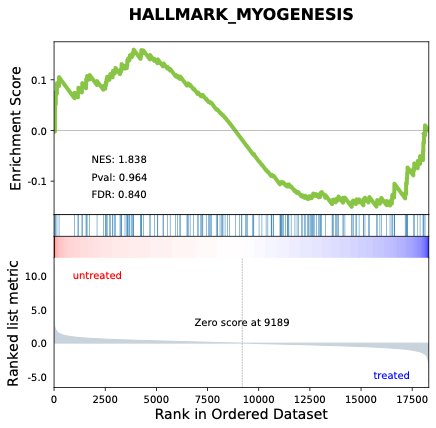

### **Supplementary Table S8. Additional Pathways GSEA Profiles (Intraperitoneal GHK-Cu).**

| **Pathway** | **Sex** | **Enrichment Score** | **NES** | ***P*-value** | **FDR** | **Regulation** |
| --- | --- | --- | --- | --- | --- | --- |
| **Estrogen Response Early** | Male | -0.301 | -2.819 | 0.023 | 0.188 | NS |
|  | Female | -0.281 | -2.378 | 0.030 | 0.217 | NS |
| **Estrogen Response Late** | Male | 0.188 | 1.757 | 0.850 | 0.895 | NS |
|  | Female | -0.167 | -1.508 | 0.860 | 0.975 | NS |
| **Spermatogenesis** | Male | 0.291 | 2.637 | 0.130 | 0.328 | NS |
|  | Female | 0.222 | 1.867 | 0.706 | 0.847 | NS |
| **Apical Junction** | Male | -0.236 | -2.376 | 0.085 | 0.337 | NS |
|  | Female | -0.312 | -2.568 | 0.002 | 0.147 | ↓ |
| **Apical Surface** | Male | -0.207 | -2.086 | 0.358 | 0.544 | NS |
|  | Female | -0.286 | -2.470 | 0.133 | 0.174 | NS |

### **Supplementary Figure S15. Male Additional Pathways GSEA Graphs (Intraperitoneal GHK-Cu).**

**A) Estrogen Response Early	B) Estrogen Response Late	C) Spermatogenesis**

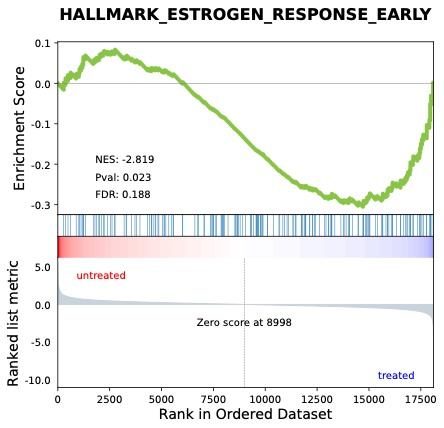

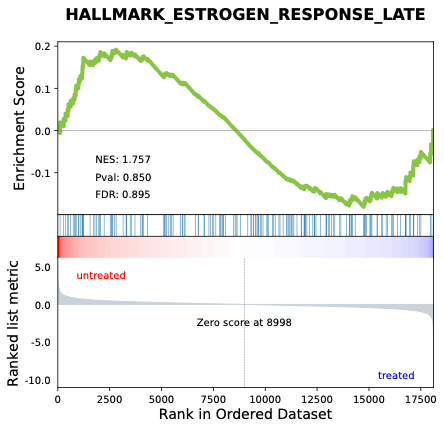

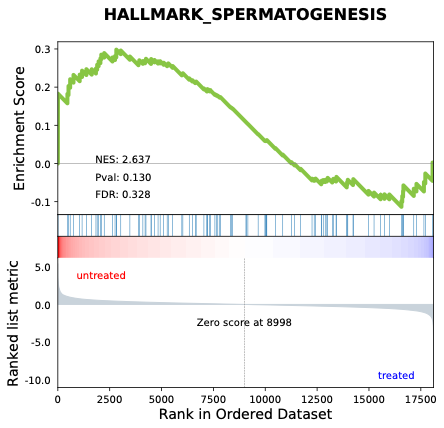

**D) Apical Junction	E) Apical Surface**

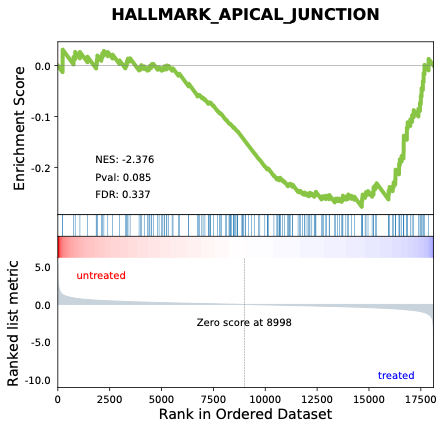

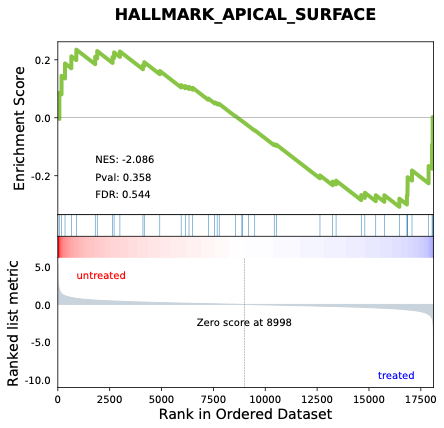

### **Supplementary Figure S16. Female Additional Pathways GSEA Graphs (Intraperitoneal GHK-Cu).**

**A) Estrogen Response Early	B) Estrogen Response Late	C) Spermatogenesis**

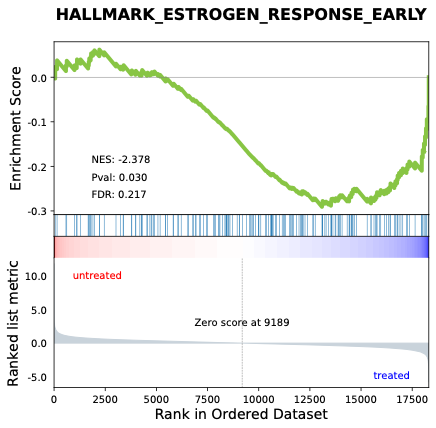

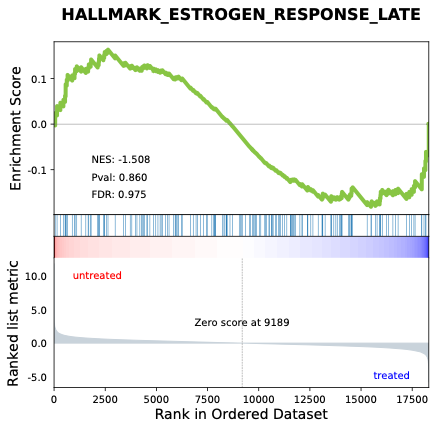

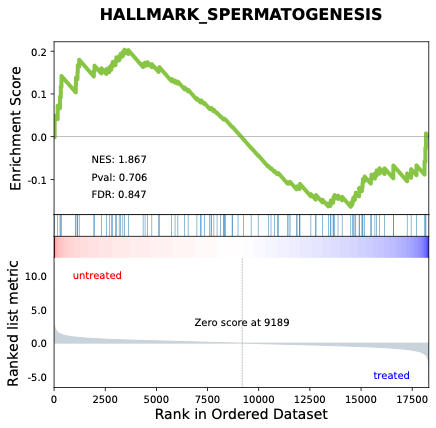

**D) Apical Junction	E) Apical Surface**

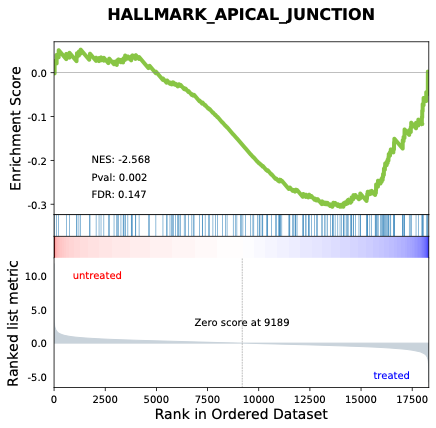

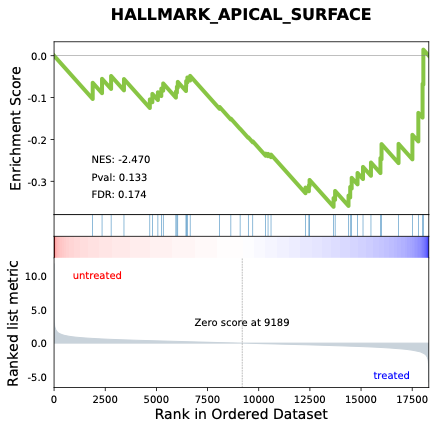
